# Evolutionary dynamics of transposable elements in bdelloid rotifers

**DOI:** 10.1101/2020.09.15.297952

**Authors:** Reuben W. Nowell, Christopher G. Wilson, Pedro Almeida, Philipp H. Schiffer, Diego Fontaneto, Lutz Becks, Fernando Rodriguez, Irina R. Arkhipova, Timothy G. Barraclough

## Abstract

Transposable elements (TEs) are selfish genomic parasites whose ability to spread autonomously is facilitated by sexual reproduction in their hosts. If hosts become obligately asexual, TE frequencies and dynamics are predicted to change dramatically, but the long-term outcome is unclear. Here, we test current theory using whole-genome sequence data from eight species of bdelloid rotifers, a class of invertebrates where males are thus far unknown. Contrary to expectations, we find a diverse range of active TEs in bdelloid genomes, at an overall frequency within the range seen in sexual species. We find no evidence that TEs are spread by cryptic recombination or restrained by unusual DNA repair mechanisms, but we report that bdelloids share a large and unusual expansion of genes involved in RNAi-mediated TE suppression. This suggests that enhanced cellular defence mechanisms might mitigate the deleterious effects of active TEs and compensate for the consequences of long-term asexuality.

## Introduction

Transposable elements (TEs) are repeated sequences of DNA that can mobilize and replicate themselves within genomes [1–3]. TEs are divided into two major categories: class I retrotransposons, which use a ‘copy-and-paste’ replication mechanism via a reverse-transcribed RNA intermediate, and class II DNA transposons, which use ‘cut-and-paste’ replication with a DNA intermediate. Both classes are ancient and diverse—retrotransposons are found in some bacteria and nearly all eukaryotes, while DNA transposons are found across the tree of life [4–6]. Although TE replications are occasionally beneficial [7], the vast majority are deleterious for the host [8,9]. Costs include insertional mutations that disrupt genes [10], cellular costs of replicating and expressing excess DNA [11], and increased risk of chromosomal abnormalities due to ectopic recombination between homologous TE sequences interspersed through the genome [12,13]. Despite this, by replicating autonomously as selfish elements, TEs can accumulate to large numbers within genomes—for example, TEs comprise 46% of the human genome, including over 1 million (∼11%) nonautonomous *Alu* retroelements [14,15]. TE numbers vary greatly, however, even between closely related species. In vertebrates, for example, TEs span an order of magnitude, from below 6% to over 50% of the genome [16], with similarly large variation observed within and between other groups such as arthropods [17], nematodes [18], and fungi [19]. Explaining this variation is vital to understanding the mechanisms affecting TE spread and control.

Sexual reproduction has long been thought to play a major role in TE dynamics within eukaryotes. On the one hand, sexual reproduction and outcrossing decouples the fate of TEs from other host genes, allowing them to jump into new genomic backgrounds and to behave as selfish genomic parasites [1,2,20]. On the other hand, sex enables the efficient removal of deleterious insertions from populations through recombination and segregation [21–23]. The risk of chromosome abnormalities due to ectopic recombination, arguably the main cost of high TE loads in eukaryotes [9,24], also occurs during chromosome pairing at meiosis. Sex therefore plays opposing roles—it permits spread and selfish behaviour of TEs, and yet it facilitates and strengthens selection against high loads. Variation in TE content among taxa might thus result from shifts in the balance of these different opposing forces.

By this logic, the loss of sexual reproduction should affect TE dynamics dramatically. Since asexual lineages generally arise from sexual species [25], it is likely that they initially harbor many active TEs [26,27]. All else being equal, the loss of recombination will limit the ability of selection to remove deleterious insertions from a fully linked host genome, and so the load of TEs should accumulate. At the same time, the fate of TEs is immediately coupled to that of the host genome, resulting in intensified selection for inactivation, excision or domestication of the elements [2,22,27–29]. The genomes of asexual lineages whose TEs continued to replicate unchecked would become overrun, potentially leading to extinction of the lineage and the TEs themselves. While some TEs could be maintained by horizontal transfer between species (especially class II DNA elements [6,30]) or by having beneficial effects (as in bacteria, [31,32]), other TEs—particularly the class I LINE-like (i.e. non–long terminal repeat [LTR]) retrotransposons—are thought to be transmitted almost exclusively vertically [5,33,34], and therefore depend strongly on sex for their persistence.

Models of the population genetics of vertically transmitted TEs in asexuals predict one of two outcomes: either TEs accumulate within lineages faster than they can be removed, overrunning each lineage in turn and driving the population extinct, or, conversely, TE removal outweighs proliferation and the population eventually purges itself entirely of deleterious TEs [27,35,36]. These predictions are difficult to test empirically, however, because the time required for a population to arrive at either extinction or redemption is expected to be on the order of millions of generations [27], too long to observe directly and beyond the lifespan of most asexual lineages [37,38].

Here, we test these ideas in a well-known group of asexual animals, the bdelloid rotifers. These microscopic invertebrates appear to have reproduced without males or meiosis for tens of millions of years, diversifying into hundreds of species within limno-terrestrial and freshwater habitats globally [39,40]. Bdelloids sampled from nature (and those reared in the laboratory) consist entirely of parthenogenetic females, and neither males nor hermaphrodites are described for any species despite centuries of close observation by naturalists [39,41,42]. Genetic and genomic evidence for their proposed ancient and obligate asexuality remains uncertain, however. Initial evidence of long-term asexuality [43,44] has been refuted by later studies or confounded by alternative explanations [45–47]. Some recent studies have proposed alternative modes of inter-individual genetic exchange, but these suggestions would require exotic mechanisms unknown in other animals [44,48], or rates of sex that are difficult to reconcile with the lack of observed males [49]. While the precise nature of reproduction in bdelloids remains an open question, nonetheless they provide a unique test-case for models of TE evolution when conventional sex is absent or strikingly rare.

Initial PCR-based surveys of five bdelloid genomes found no evidence of class I retrotransposons from either the LTR or LINE-like superfamilies, but did reveal a diverse array of class II DNA transposons, mostly at low copy number [50]. The presence of class II TEs in bdelloids might be explained by horizontal transfer, which is thought to occur more frequently for class II TEs with DNA intermediates [6,30,33,34] (but see [51]). The apparent lack of retrotransposons contrasted sharply, however, with their near ubiquity in other taxa. At the time, the absence of class I TEs appeared consistent with the view that long-term asexual evolution in bdelloids had caused the loss of parasitic elements that depended on sexual transmission [22,27,50,52].

Another unusual aspect of bdelloid physiology was suggested to contribute to their seemingly low TE complement. In most bdelloid species (but not all), individuals can survive complete desiccation at any life stage via a process called anhydrobiosis (‘life without water’). Desiccation causes double-strand breakages (DSBs) in DNA, but bdelloids are able to repair these and recover to an unusual degree [53–55]. It was proposed that anhydrobiosis might influence TE evolution in two ways [27,52,56]. First, DSB repair could aid TE removal, either via gene conversion from a homologous chromosome lacking the TE insertion, or excision of mis-paired regions. Second, the pairing of homologous chromosomes, if required during DSB-repair, could provide a context for ongoing selection against chromosomal abnormalities caused by ectopic recombination. In either case, anhydrobiosis would decrease the number of TEs, potentially helping to explain the low overall TE content encoded in bdelloid genomes.

These early ideas were transformed by more detailed studies of the model bdelloid species *Adineta vaga*, which used refined methods and genome-scale data to discover a variety of retrotransposon families. These include an endonuclease-deficient *Penelope*-like element (PLE) designated *Athena* [57,58], which is itself incorporated within much larger and highly unusual retroelements called *Terminons* [59]; another PLE that has retained its endonuclease [60], LTR retrotransposons (*Juno, Vesta, TelKA* and *Mag* [61,62]), and LINE-like retrotransposons (*R4, R9, Hebe, RTE, Tx1* and *Soliton* [44,63,64]). In total, TEs accounted for 2.2% of the 217 Mb genome (∼4.8 Mb) [44], rising to ∼4% on inclusion of the recently discovered giant *Terminon* elements [59]. The conclusion that bdelloids lack class I TEs therefore no longer holds, and the predicted effects of asexuality and anhydrobiosis on TE evolution remain open questions that are amenable to testing with comparative genomics. Specifically, the comparison of desiccating species with those few bdelloid rotifer lineages that are unable to survive desiccation would indicate whether anhydrobiosis does limit TE numbers as hypothesized. Also, comparisons within populations could shed light on the activity of TEs and whether bdelloids possess cryptic forms of recombination that could aid in TE removal or facilitate TE spread.

Here, we test the predicted effects of asexuality and anhydrobiosis on TE evolution by comparing 42 rotifer genomes belonging to 15 taxonomic species. Our sample includes both desiccating and nondesiccating bdelloids, and eight monogonont rotifers [65–67], a separate class that alternates sexual and asexual reproduction and cannot survive desiccation as adults. Results are set in context by comparison to published genomes from an acanthocephalan [68] (now classified with rotifers in the Phylum Syndermata) and a range of other animal phyla. We ask six questions raised by theory. (1) How does the abundance and diversity of TEs in bdelloids differ from that in other animals, including sexual rotifers? (2) Are TEs recently and currently active in all bdelloids? (3) Can we detect signatures of recombination affecting bdelloid TEs, as might be expected if bdelloids do have cryptic forms of sexual reproduction? (4) Do desiccating species contain fewer TEs than nondesiccating species, as previously theorized? (5) Are TEs in bdelloids under the same selective constraints as in other taxa? (6) Finally, do bdelloids possess alternative molecular mechanisms that might help to keep their TEs under control, as might be expected if the ability to control TEs by sexual reproduction is absent or reduced?

## Results and Discussion

### High-quality population genomics data for bdelloid rotifers

To quantify variation in repeat content within and between bdelloid species, we generated *de novo* whole-genome assemblies for 31 rotifer samples encompassing nine species (**Fig 1a**; **Table 1**; S1 Data). Three of these assemblies were generated using 10x Genomics linked-read data (for *Adineta steineri, Rotaria sordida*, and *Rotaria* sp. ‘Silwood-1’), while 26 are from single-individual samples. In order to capture as many potential repeats as possible, we generated two assemblies for each sample: a ‘reference’ assembly, with a focus on quality and contiguity, and a ‘maximum haplotype’ (maxhap) assembly that included small or highly similar contigs that might be derived from recent TE duplications or other sources of copy number variation, at the expense of contiguity.

**Table 1.** Assembly statistics for one monogonont and 30 bdelloid rotifer reference assemblies presented in this study.

| Sample ID | Species name | SZ (Mb) | NN | N50 (kb) | L50 | AU (kb) | GC (%) | Gaps (kb) | Coverage (X) | Genome BUSCO score | CDS | Proteome BUSCO score | GenBank accession |
| --- | --- | --- | --- | --- | --- | --- | --- | --- | --- | --- | --- | --- | --- |
| **Bc_PSC1** | *Brachionus calyciflorus* (Monogonont) | 116.7 | 14,869 | 18.5 | 1,692 | 26.6 | 25.6 | 78 | 186 | C:96%[S:93%,D:3%],F:2% | 24,404 | C:98%[S:93%,D:5%],F:1% | GCA_904009505.1 |
| **Ar_ARIC003** | *Adineta ricciae* | 135.6 | 4,302 | 283.8 | 129 | 388 | 35.5 | 65 | 89 | C:97%[S:58%,D:39%],F:2% | 49,015 | C:97%[S:52%,D:45%],F:1% | GCA_904047305.1 |
| **As_10x_p** | *Adineta steineri* | 171.1 | 8,257 | 200.1 | 163 | 394.5 | 29 | 206 | 198 | C:95%[S:67%,D:33%],F:2% | 50,321 | C:97%[S:58%,D:38%],F:2% |  |
| **As_ASTE804** | *Adineta steineri* | 160.3 | 9,359 | 158.1 | 265 | 214.9 | 29.1 | 152 | 62 | C:95%[S:74%,D:22%],F:2% | 47,222 | C:98%[S:74%,D:24%],F:2% | GCA_904047245.1 |
| **As_ASTE805** | *Adineta steineri* | 156.3 | 9,008 | 169.6 | 245 | 226.4 | 29.2 | 129 | 65 | C:98%[S:77%,D:21%],F:1% | 43,986 | C:99%[S:72%,D:26%],F:1% | GCA_904047255.1 |
| **As_ASTE806** | *Adineta steineri* | 160.3 | 7,597 | 168.2 | 257 | 222.5 | 29.2 | 145 | 82 | C:96%[S:72%,D:24%],F:2% | 45,930 | C:98%[S:74%,D:24%],F:2% | GCA_904047275.1 |
| **Dc_DCAR505** | *Didymodactylos carnosus* | 323.6 | 87,048 | 7.8 | 11,656 | 10.5 | 33.5 | 41 | 21 | C:86%[S:69%,D:17%],F:8% | 46,286 | C:88%[S:71%,D:18%],F:9% | GCA_904047325.1 |
| **Dc_DCAR706** | *Didymodactylos carnosus* | 368.8 | 78,356 | 12 | 7,695 | 19.1 | 33.5 | 13 | 76 | C:95%[S:70%,D:25%],F:2% | 46,863 | C:95%[S:71%,D:25%],F:2% | GCA_904053135.1 |
| **Rd_10x_p** | *Rotaria sordida* | 272.5 | 16,571 | 64.5 | 843 | 193.8 | 30.8 | 395 | 91 | C:94%[S:77%,D:19%],F:2% | 44,299 | C:95%[S:69%,D:26%],F:2% |  |
| **Rd_RSOR408** | *Rotaria sordida* | 252.9 | 20,315 | 57.6 | 1,246 | 75.5 | 30.4 | 291 | 39 | C:94%[S:76%,D:19%],F:3% | 40,501 | C:97%[S:73%,D:24%],F:2% | GCA_904053145.1 |
| **Rd_RSOR410** | *Rotaria sordida* | 252.6 | 19,518 | 60.9 | 1,179 | 80 | 30.4 | 252 | 42 | C:95%[S:77%,D:18%],F:3% | 40,474 | C:98%[S:74%,D:24%],F:2% | GCA_904053495.1 |
| **Rd_RSOR504** | *Rotaria sordida* | 251.3 | 22,067 | 53.1 | 1,338 | 69.8 | 30.4 | 369 | 39 | C:94%[S:78%,D:16%],F:3% | 41,085 | C:96%[S:73%,D:23%],F:3% | GCA_904047315.1 |
| **Rg_MAG1** | *Rotaria magnacalcarata* | 178.7 | 19,184 | 42 | 1,077 | 62.4 | 32 | 402 | 58 | C:97%[S:81%,D:16%],F:1% | 40,318 | C:99%[S:76%,D:22%],F:1% | GCA_903995865.1 |
| **Rg_MAG2** | *Rotaria magnacalcarata* | 181.1 | 22,216 | 39.7 | 1,141 | 61 | 32 | 433 | 63 | C:98%[S:81%,D:17%],F:1% | 40,289 | C:99%[S:74%,D:26%],F:0% | GCA_903995855.1 |
| **Rg_MAG3** | *Rotaria magnacalcarata* | 180.9 | 22,132 | 40.7 | 1,142 | 60 | 32 | 508 | 60 | C:96%[S:80%,D:17%],F:1% | 40,740 | C:99%[S:77%,D:22%],F:0% | GCA_903995845.1 |
| **Rg_RM15** | *Rotaria magnacalcarata* | 174 | 18,391 | 46.5 | 966 | 67.3 | 32 | 430 | 55 | C:96%[S:80%,D:16%],F:2% | 38,283 | C:99%[S:77%,D:22%],F:1% | GCA_903995885.1 |
| **Rg_RM9** | *Rotaria magnacalcarata* | 173.8 | 19,520 | 44 | 999 | 64.7 | 31.9 | 594 | 51 | C:96%[S:80%,D:16%],F:1% | 38,404 | C:98%[S:76%,D:22%],F:1% | GCA_903995825.1 |
| **Rp_RPSE411** | *Rotaria* sp. 'Silwood-2' | 296.5 | 30,050 | 102.5 | 381 | 691.1 | 31 | 247 | 35 | C:93%[,S:72%,D:21%],F:4% | 48,378 | C:95%[S:72%,D:23%],F:4% | GCA_904053095.1 |
| **Rp_RPSE503** | *Rotaria* sp. 'Silwood-2' | 285.6 | 33,174 | 78.3 | 449 | 627.2 | 31.3 | 446 | 34 | C:91%[,S:75%,D:17%],F:5% | 48,269 | C:92%[S:72%,D:20%],F:7% | GCA_904053125.1 |
| **Rp_RPSE809** | *Rotaria* sp. 'Silwood-2' | 271.1 | 28,589 | 101.6 | 350 | 681.4 | 31 | 377 | 27 | C:93%[,S:76%,D:17%],F:4% | 47,010 | C:95%[S:74%,D:22%],F:4% | GCA_904053085.1 |
| **Rp_RPSE812** | *Rotaria* sp. 'Silwood-2' | 264.1 | 34,498 | 80.8 | 403 | 616.2 | 31.1 | 428 | 27 | C:89%[,S:74%,D:15%],F:8% | 47,040 | C:90%[S:73%,D:17%],F:8% | GCA_904053105.1 |
| **Rs_AK11** | *Rotaria socialis* | 149.2 | 6,303 | 111.3 | 370 | 150.2 | 31.8 | 442 | 39 | C:97%[,S:80%,D:17%],F:1% | 34,844 | C:99%[S:75%,D:24%],F:1% | GCA_903995835.1 |
| **Rs_AK15** | *Rotaria socialis* | 147.4 | 5,030 | 134.7 | 305 | 177.6 | 31.8 | 423 | 37 | C:96%[,S:79%,D:18%],F:1% | 34,140 | C:98%[S:76%,D:23%],F:1% | GCA_903995795.1 |
| **Rs_AK16** | *Rotaria socialis* | 147.4 | 4,720 | 139.5 | 296 | 180.3 | 31.8 | 332 | 43 | C:97%[,S:80%,D:18%],F:0% | 33,717 | C:99%[S:76%,D:23%],F:1% | GCA_903995805.1 |
| **Rs_AK27** | *Rotaria socialis* | 149.9 | 5,952 | 123.7 | 343 | 159.8 | 31.8 | 458 | 36 | C:97%[,S:80%,D:17%],F:0% | 34,369 | C:99%[S:75%,D:24%],F:1% | GCA_903995815.1 |
| **Rs_RS1** | *Rotaria socialis* | 151.1 | 6,254 | 124.9 | 334 | 166.2 | 31.8 | 490 | 40 | C:97%[,S:80%,D:17%],F:0% | 33,937 | C:99%[S:77%,D:22%],F:1% | GCA_903995875.1 |
| **Rw_10x_p** | *Rotaria* sp.<br>'Silwood-1' | 310.4 | 16,995 | 211.8 | 211 | 126.2 | 31.1 | 534 | 53 | C:95%,[S:76%,D:20%],F:1% | 44,241 | C:97%[S:73%,D:24%],F:1% |  |
| **Rw_RSIL801** | *Rotaria* sp.<br>'Silwood-1' | 268.4 | 28,548 | 136.5 | 288 | 687.3 | 30.8 | 472 | 45 | C:94%,[S:77%,D:17%],F:4% | 41,574 | C:95%[S:75%,D:21%],F:5% | GCA_904053155.1 |
| **Rw_RSIL802** | *Rotaria* sp.<br>'Silwood-1' | 249.9 | 21,286 | 153.4 | 238 | 702.3 | 30.7 | 451 | 42 | C:92%,[S:76%,D:16%],F:4% | 39,577 | C:94%[S:76%,D:18%],F:4% | GCA_904053505.1 |
| **Rw_RSIL804** | *Rotaria* sp.<br>'Silwood-1' | 247.6 | 25,643 | 118.3 | 287 | 660.4 | 30.8 | 667 | 34 | C:94%,[S:78%,D:16%],F:3% | 41,139 | C:96%[S:78%,D:19%],F:3% | GCA_904053115.1 |
| **Rw_RSIL806** | *Rotaria* sp.<br>'Silwood-1' | 294.1 | 29,968 | 132.4 | 333 | 681.9 | 30.8 | 500 | 31 | C:95%,[S:79%,D:16%],F:2% | 48,259 | C:97%[S:78%,D:19%],F:2% | GCA_904054495.1 |

Sequence statistics codes: **SZ**, total sequence length (Mb); **NN**, number of sequences; **N50**, N50 scaffold length (kb); **L50**, N50 index; **AU**, expected scaffold size (area under 'Nx' curve, kb).
Gaps: Undetermined bases (Ns) introduced during scaffolding.
BUSCO score based on eukaryote set ( $n = 303$ ); BUSCO codes: **C**, complete; **S**, complete and single copy; **D**, complete and duplicated; **F**, fragmented.
Abbreviations: GC, guanine + cytosine; BUSCO, Benchmarking Universal Single-Copy Orthologs.

**Fig 1.**
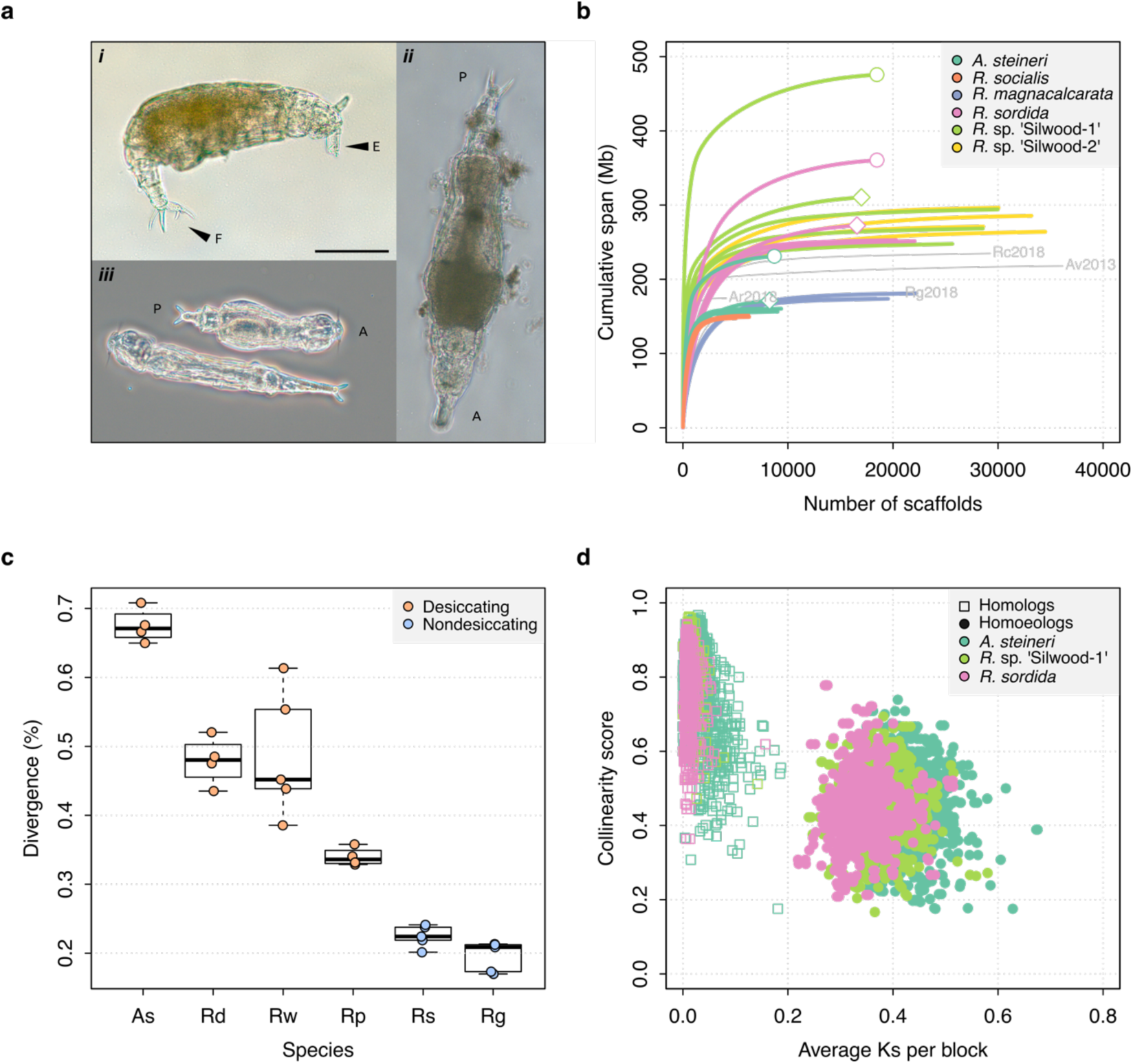
Genome properties of sequenced rotifers. **(a)** Bdelloid rotifer morphology; scale bar indicates 100 µm. (*i*) Individual from an undescribed species of *Rotaria* (*R*. sp. ‘Silwood-1’), showing eyes (E) and foot (F) with two spurs and three toes. (*ii*) Further image of *R*. sp. ‘Silwood-1’ with anterior–posterior (A–P) axis marked. (*iii*) Two individuals of *A. steineri* in phase contrast. **(b)** Cumulative assembly span for six bdelloid species with population genomics data (*n* > 2). 10x Genomics haploid (‘pseudohap’) and diploid (‘megabubbles’) assemblies for *A. steineri, R. sordida* and *R*. sp. ‘Silwood-1’ are indicated with diamond and circle symbols, respectively. The four previously published genomes for *A. vaga* (‘Av2013’, GenBank accession GCA_000513175.1) and *A. ricciae* (‘Ar2018’, GCA_900240375.1), *R. macrura* (‘Rc2018’, GCA_900239685.1) and *R. magnacalcarata* (‘Rg2018’, GCA_900239745.1) are indicated in grey, for comparison. **(c)** Intragenomic divergence, measured as the number of SNPs detected in coding regions (CDS). Boxplots show the median (band), interquartile range (box) and minimum/maximum values (whiskers). Underlying data are shown as jittered points. Desiccation-tolerant species are in orange, intolerant species in blue. Species abbreviations: **As**, *A. steineri*; **Rd**, *R. sordida*; **Rw**, *R*. sp. ‘Silwood-1’; **Rp**, *Rotaria* sp. ‘Silwood-2’; **Rg**, *R. magnacalcarata*; **Rs**, *R. socialis*. **(d)** Genome structure in *A. steineri, R. sordida* and *R*. sp. ‘Silwood-1’ haplotype-resolved (‘megabubbles’) assemblies. Each point represents a collinear block of genes, plotted by average pairwise synonymous (*K*S, *X*-axis) and collinearity score (see Materials and methods and S1 note) on the *Y*-axis. Separation into two distinct clusters representing homologous (squares) and homoeologous (circles) relationships among gene copies is consistent with ancestral tetraploidy, with homoeologous copies derived from a putative ancient genome duplication.

Reference genomes showed an expected scaffold size (AU, see Materials and methods) ranging from 21.1 kb (*Didymodactylos carnosus*) to 702.3 kb (*R*. sp. ‘Silwood-1’) and BUSCO scores that indicated 89–98% of 303 core eukaryote genes were completely recovered, increasing to 96– 99% if fragmented copies are included (**Table 1**). General genome characteristics such as genome size (assembly span), the proportion of G + C nucleotides (GC%), the number of coding genes (CDS), and the level of homologous divergence (number of SNPs identified within CDS) were within the range expected from previous analyses of bdelloid genomes [44,46] (**Fig 1b–c**; **Table 1**; S1 Fig). Intragenomic collinearity and synonymous divergence of coding regions in the *A. steineri, R. sordida* and *R*. sp. ‘Silwood-1’ maximum haplotype assemblies reveal the characteristic signature of degenerate tetraploidy that has been found in all bdelloid species examined to date (**Fig 1d**).

Compared to the reference set, maxhap assemblies generally showed increased span (mean increase = 17.9 Mb ± 21.5 standard deviation [SD]) and were substantially more fragmented, as expected (S1 Table). Nonetheless, BUSCO completeness scores remained high, with 76% to 98% of genes completely recovered (increasing to 95–98% if fragmented copies are included), indicating that the majority of core genes are successfully captured (S1 Table). The BUSCO duplication metric (‘D’) does not increase greatly between reference and maxhap assemblies, which shows that the additional sequences retained in the maxhap assemblies do not contain complete extra copies of core genes. Thus, the maxhap assemblies are not fully haplotype-resolved representations of the genome, except in the case of the three 10x assemblies.

To these new data, we added published genomes for seven monogonont taxa (one from the *Brachionus calyciflorus* species complex [65] and six from four species of the *Brachionus plicatilis* species complex, namely *B. asplanchnoidis, B. plicatilis* sensu stricto (HYR1), *B. rotundiformis*, and *B*. sp. ‘Tiscar’ [66,67]) and four bdelloids (*A. vaga, Adineta ricciae, Rotaria magnacalcarata* and *Rotaria macrura* [44,46]), yielding a total of 42 rotifer genomes. Of these, 11 samples belong to nondesiccating bdelloid species (five individuals each from *R. magnacalcarata* and *Rotaria socialis*, and the previously published genome of *R. macrura*).

### Abundant and diverse TEs in bdelloid genomes

To ascertain the repeat content of bdelloid genomes relative to other taxa in a consistent manner, we used the RepeatModeler and RepeatMasker pipelines to identify and classify repeats across genomes. The total proportion of the genome classified as repetitive ranged from ∼19% to 45% across bdelloid genera, with variation within and between species (**Fig 2a**; S2 Fig; S3 Fig; S2 Data; S3 Data). Most of these are simple or unclassified repeats that do not belong to major TE superfamilies. While the precise nature of these unclassified repeats is not elucidated, an appreciable fraction (∼7–27%, mean = 17%) are also annotated as protein-coding and thus may be derived from gene expansions or other duplications, while a further small fraction (< 1%) are accounted for by an additional survey of small, nonautonomous class II TEs called miniature inverted-repeats (MITEs) (S2 Data). The proportion of the genome accounted for by known TEs was much smaller, ranging from 2.4% to 7.3% (mean = 4.9% ± 1.2 standard deviations [SD], median = 5.1%) in bdelloids. Broken down by class and superfamily, the mean values are: class I total = 2.09% ± 0.75 (PLEs = 0.59% ± 0.14; LTRs = 0.68% ± 0.26; and LINEs = 0.82% ± 0.47); class II total = 2.79 ± 0.8 (DNA transposons = 2.49% ± 0.77; rolling circles = 0.30 ± 0.11). These results are in broad agreement with previous estimates of TE content in bdelloids [44,46,47,59].

**Fig 2.**
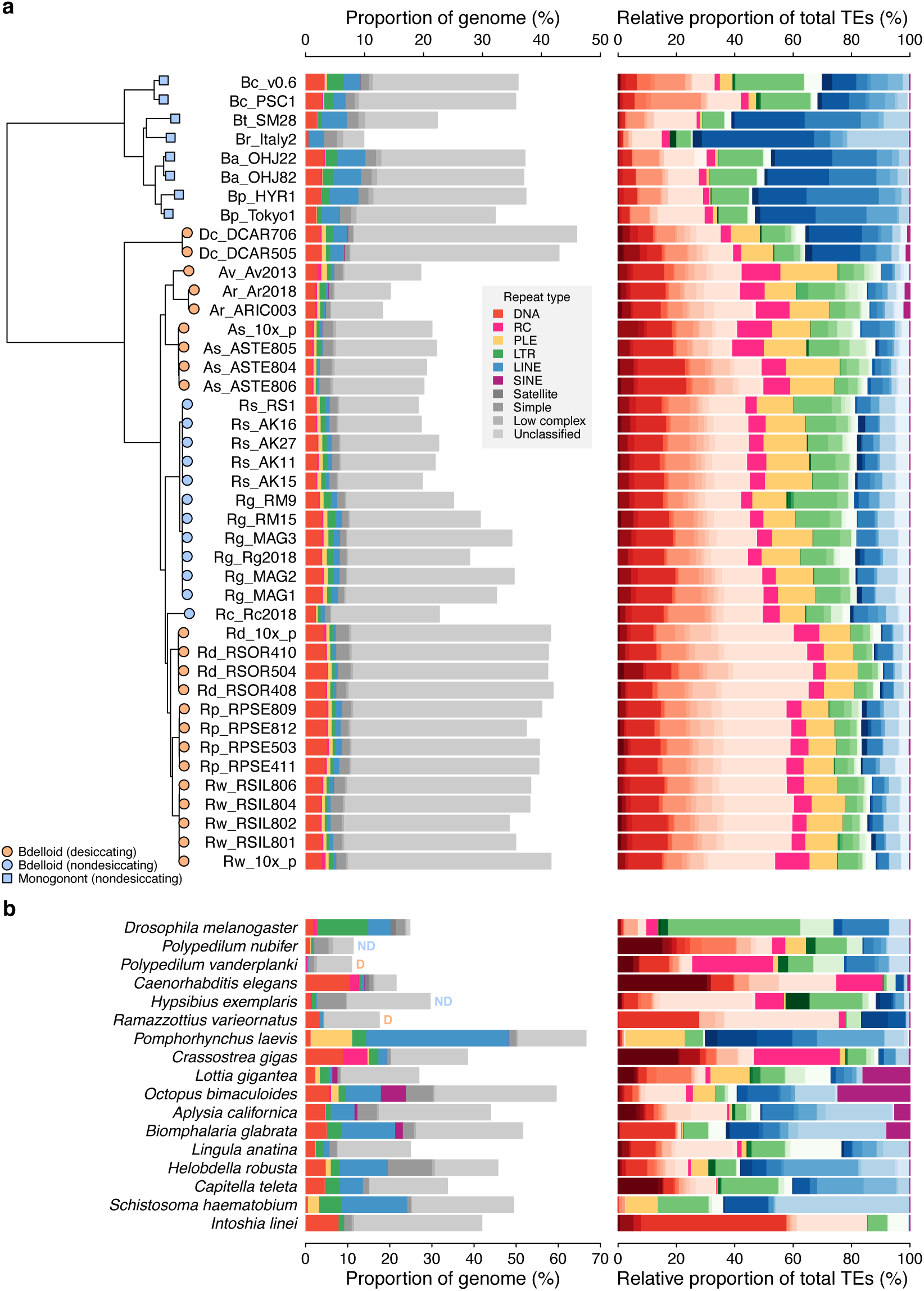
Repeat content and diversity in rotifer genomes. **(a)** Maximum likelihood phylogeny of eight monogonont (square symbols on tips) and 34 bdelloid (circles) genomes based on the concatenated alignment of a subset of core eukaryotic (BUSCO) genes. Orange and blue tip colours indicate desiccating and nondesiccating taxa, respectively. Species codes in tip names are: **Bc**, *Brachionus calyciflorus*; **Br**, *B. rotundiformis*; **Bt**, *B*. sp. ‘Tiscar’; **Bp**, *B. plicatilis* HYR1; **Ba**, *B. asplanchnoidis*; **Dc**, *Didymodactylos carnosus*; **Av**, *Adineta vaga*; **Ar**, *A. ricciae*; **As**, *A. steineri*; **Rs**, *Rotaria socialis*; **Rg**, *R. magnacalcarata*; **Rc**, *R. macrura*; **Rd**, *R. sordida*; **Rw**, *R*. sp. ‘Silwood-1’; **Rp**, *R*. sp. ‘Silwood-2’. Repeat content is shown as the genome proportion (%) broken down by TE superfamily (middle panel), and relative proportion (%) of total known (i.e., classified) TEs (right panel), where colours represent TE superfamilies (see legend) and shades of colour represent different TE families within each superfamily. **(b)** Equivalent repeat content analysis in 17 protostome animal genomes, including the model species *D. melanogaster* and *C. elegans*, the recently published acanthocephalan rotifer *P. laevis*, and selected other species from across the protostome group. Two further examples of desiccating (orange ‘D’) and nondesiccating (blue ‘ND’) species pairs are shown: the insects *P. nubifer* and *P. vanderplanki* and the tardigrades *H. exemplaris* and *R. varieornatus*.

We first compare these values to closely related sexual lineages in the phylum Syndermata, the monogonont rotifers and acanthocephalans. The most striking difference relates to the recently published genome of the obligately sexual acanthocephalan *Pomphorhynchus laevis* [68], which encodes a substantially greater proportion of repeats than either bdelloids or monogononts. In agreement with Mauer et al., we find ∼66% of the *P. laevis* genome to be composed of repeats. The large majority are class I retrotransposons (∼71% of the total TE content) from the LINE (∼52%) and PLE (∼15%) superfamilies, and there are relatively few DNA transposons (∼1.3%) (**Fig 2b**). There is increasing evidence that acanthocephalans may be the closest relatives to the Class Bdelloidea [69–71]. However, all members of the Acanthocephala (also known as the thorny-headed worms) are obligate endoparasites and are highly differentiated in both morphological and molecular terms from other syndermatans. It is unclear whether these features may contribute to the high repeat content and other unusual genome characteristics of *P. laevis* [68].

In monogononts, the mean values for class I TEs is 5.2% ± 1.5 SD and for class II TEs it is 2.5% ± 1.0 SD. Thus, monogononts are slightly more TE-rich than bdelloids, but also substantially more variable between species (**Fig 2a**; S3 Fig). Repeat content differs between bdelloids and monogononts in two main ways, both in regard to the composition of class I retrotransposons. First, monogononts encode substantially more LINE-like retroelements than bdelloids, making up (on average) approximately 50% and 16% of the total TE content in each clade respectively. The frequency of LINEs is of particular interest, because this class of TEs is thought to be least likely to undergo horizontal transfer and thus the most dependent on sex for transmission [5,33,34]. Second, *Penelope*-like elements (PLEs) have increased in proportion in all bdelloids relative to monogononts, from ∼1% in monogononts to ∼12% in bdelloids on average.

Interestingly, a high proportion of PLEs is also seen in the acanthocephalan *P. laevis*, while a high proportion of LINEs is found in both *D. carnosus* isolates (∼35% of total TE content), a deeply branching lineage sister to all other bdelloid taxa included in the analyses. Thus, assuming that acanthocephalans are the closest relatives to bdelloids, the most parsimonious explanation for these broad-scale patterns is that the expansion of PLEs occurred in the ancestor to bdelloids and acanthocephalans, whereas the contraction of LINEs has occurred more recently, confined to a subset of bdelloid genera.

To put these differences in TE content among rotifers in a broader context, we applied the same repeat-finding pipeline to animals from a range of more distantly related protostome phyla: three insects, a nematode, two tardigrades, five molluscs, two annelids, a brachiopod, platyhelminth and orthonectid [72–83] (S2 Table). As expected, both the abundance and diversity of TEs varied widely across taxa (**Fig 2b**). Total TE content ranged from 0.8% (the insect *Polypedilum vanderplanki*) to ∼24% (octopus and platyhelminth). Although the acanthocephalan genome appears to be particularly rich in TEs, bdelloids (except for *D. carnosus*) have modest amounts of TEs, including class I TEs specifically, while in monogononts these amounts vary greatly among species. All bdelloids encode relatively more TEs than both *Polypedilum* species but fewer than *Drosophila melanogaster, Caenorhabditis elegans*, annelid worms and some molluscs, and are intermediate with respect to other taxa. Note that TE proportion in molluscs, nematodes and flatworms is known to be highly variable (e.g. [18]), while bdelloids display much less variability barring the early-branching *D. carnosus*.

These results show that bdelloid species encode an abundant diversity of both class I and II TEs, and, set against the repeat content of other rotifers and animals, do not appear particularly deficient or unusual with regard to the proportion of nucleotides encoding transposable elements. Although the bdelloids do have lower frequencies of class I TEs (∼16% of total TE content) than the monogononts (∼50%) or the acanthocephalan (∼52%), as predicted by theory for elements dependent on vertical transmission, they still possess them in numbers that are comparable to sexual organisms (e.g. *C. elegans*). Furthermore, the basally divergent bdelloid *D. carnosus* did not show the same magnitude of decrease in LINEs (∼35%), indicating that different dynamics may be at play in different lineages. Thus, it is clear that the most simplistic expectations of TE evolution under the hypothesis of long-term asexuality (i.e., either runaway proliferation or complete elimination) are not met, necessitating an evaluation of possible explanations that may align theory with observation.

### TE transposition is recent and ongoing

One possible explanation is that TEs in bdelloid genomes do not replicate autonomously or are otherwise inactivated or ‘fossilised’ within their host genome. To investigate this, we first generated divergence ‘landscapes’ for identified TE copies within each genome, using the de novo RepeatMasker results. TE landscapes measure the amount of sequence divergence between each TE copy and a consensus derived from all copies in its family [84]. Histograms of the resulting Kimura distances (*K*-values [85]) provide insights into the evolutionary history of TE activity [16,86,87].

TE landscapes for the three diploid (10x Genomics) assemblies of *A. steineri, R*. sp. ‘Silwood-1’ and *R. sordida* show that TE divergence is bimodal but strongly zero-inflated (**Fig 3**). A large number of TE copies have very low or no divergence from the consensus (*K*-value ≤ 1%). Assuming a molecular clock for nucleotide substitutions within duplicated TEs, such elements represent recent duplications that are highly similar to their progenitor copy, consistent with recent transposition of an active element. In proportion, most of these belong to class II DNA transposon superfamilies (in red), but the spike of zero divergence is also present for class I retrotransposons (in blue and green). An older, broader mode is seen around a *K*-value of 20– 30% that probably reflects historical TE transpositions and/or a signal from the tetraploid genome structure present in all bdelloids sequenced to date. The same pattern was observed in the haplotype-resolved assemblies of *A. vaga* [44] and *A. ricciae* [46] (**Fig 3b**; S4 Fig), and was generally present but less pronounced in the other ‘maxhap’ assemblies depending on the repeat pipeline applied (S4 Fig; S5 Fig).

**Fig 3.**
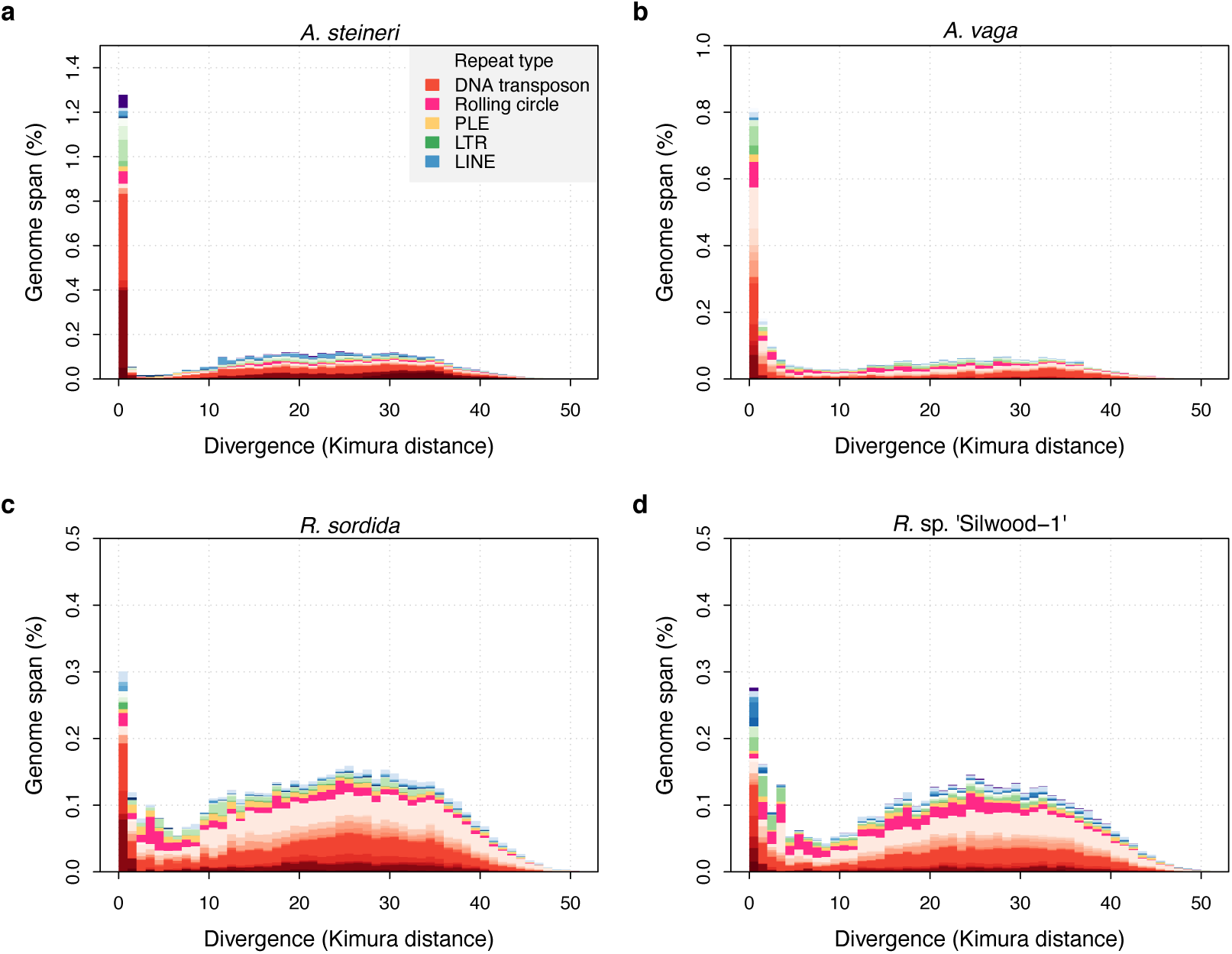
TE divergence landscapes for selected genomes. The *X*-axes show the level of divergence (Kimura substitution level, CpG adjusted) between each identified TE copy and the consensus sequence for that TE family (the inferred ancestral copy). Thus, if newly arising TE copies evolve neutrally, the amount of divergence is a proxy for the time since its duplication, with older copies accumulating more substitutions and appearing further to the right. The *Y*-axis shows the proportion of the genome occupied by each bin. Colours represent TE superfamilies (see legend) and shades of colour represent different TE families within each superfamily. Data are shown for the 10x Genomics diploid (‘megabubbles’) assemblies of *A. steineri, R*. sp. ‘Silwood-1’ and *R. sordida* compared to the published genome of *A. vaga*. Note different scales on some *Y*-axes.

To evaluate recent TE activity further, we developed a simple method to identify insertion sites for LTR retrotransposons (LTR-Rs) and assess their presence or absence in related individuals. Because most LTR-Rs insert into random genome locations [5,8], the neighbouring genome sequence provides a unique marker for a given insertion event [88]. We constructed a library of such insertion markers (‘LTR-tags’) for all ‘full-length’ LTR-Rs (i.e. those with long-terminal repeats present at both the 5’ and 3’ ends of the element) detected in our genomes, and then searched for their presence or absence in the other samples. For a given LTR-tag identified in genome *A*, the presence of a contiguous alignment in genome *B* indicates that the same insertion is shared between *A* and *B*.

For a set of 161 high-confidence and non-redundant LTR-Rs identified in the single-individual samples, alignment contiguity for each LTR-tag versus each of the other genomes was scored using a read-mapping approach (see Materials and methods), resulting in a pairwise matrix of presence/absence scores (**Fig 4a**; S4 Data). High scores for LTR insertion-site presence correlated strongly with the phylogeny, resulting in an average score of ∼0.9 within species compared to < 0.1 between species and a clear visual signal along the diagonal of Fig 4a. Very few LTR insertion sites were shared between bdelloid species. While some absences could reflect loss rather than gain, the restriction of nearly all LTR insertion sites to single species indicates that they have been gained during the separate evolutionary history of that species.

**Fig 4.**
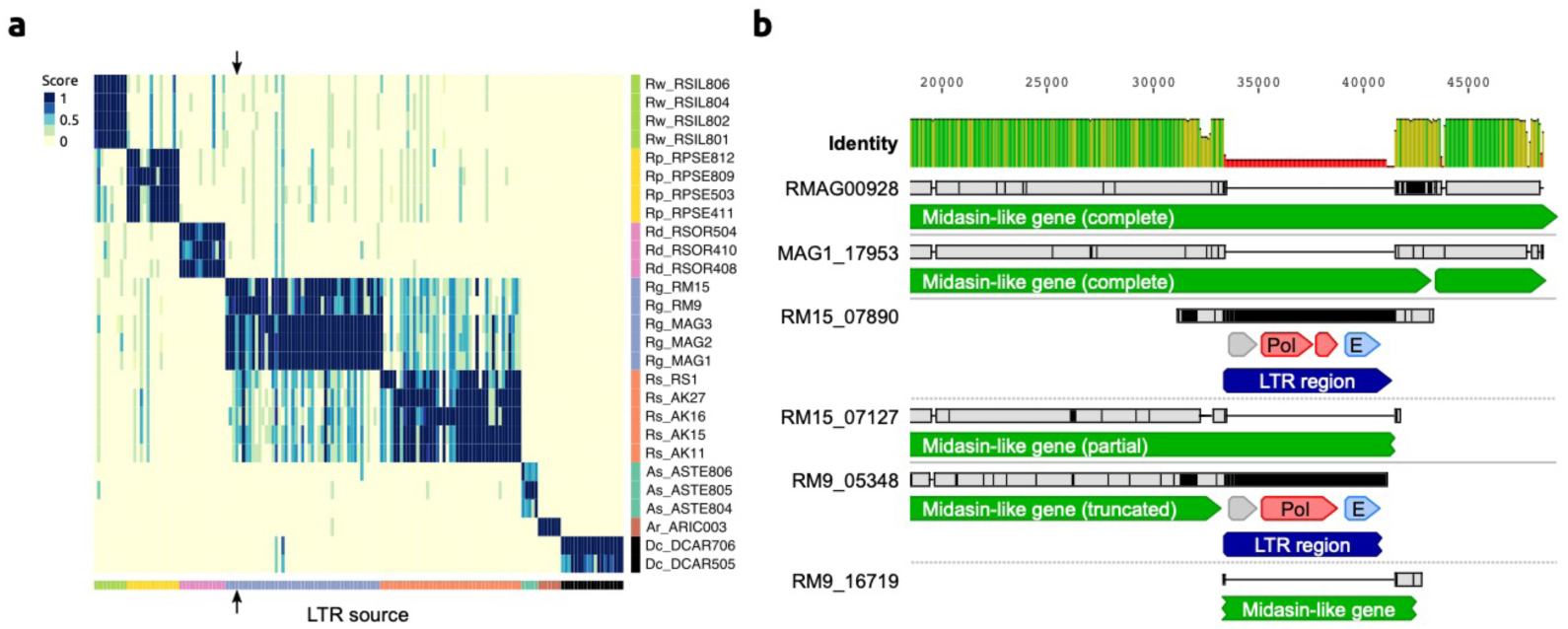
LTR insertion-site polymorphism in bdelloid species. **(a)** Columns represent 161 LTR-Rs identified across bdelloid samples, arranged by genome of origin (see colours at bottom and side). Support for the presence of a given LTR-R at a specific insertion site in each genome is scored from 0 (absent, yellow) to 1 (present, dark blue), where a score < 0.5 is strong evidence for absence (see Materials and methods for details). Arrows demark the location of the LTR-R example shown in b. **(b)** Nucleotide alignment of region around an LTR-R insertion (blue) identified in RM9 (scaffold 05348) and RM15 (scaffold 07890), alongside their putative homologous scaffolds (scaffolds 16719 and 07127 respectively) that do not show the insertion. Scaffolds from Rg2018 (RMAG00928) and MAG1 are also shown for comparison. Predicted CDS with similarity to Pol and Env proteins are shown in red and light blue. The LTR-R is most likely a member of the *TelKA* family, based on sequence similarity.

LTR-R insertions also vary between individuals within the same species, indicating recent transposition events and the potential for ongoing fitness consequences for the host. One case-study is illustrated for *R. magnacalcarata* (**Fig 4b**; S6 Fig). The individuals RM9 and RM15 share an LTR-R insertion that is not present in conspecifics. Aligning the regions of the genome assemblies containing these LTR-tags indicates that an 8.1 kb LTR-R has inserted into a protein-coding sequence in the lineage leading to RM9 and RM15. It has introduced a premature stop codon to a gene that encodes a protein (7,479 residues) of unknown function but with partial similarity to midasin, an ATPase essential to ribosome biosynthesis in several model eukaryotes [89,90]. In RM9 and RM15, the predicted product is substantially truncated (to 6,025 residues) by the element insertion. Despite the potential fitness consequences, RM9 and RM15 have evidently persisted for some time since, because they differ at approximately 0.5% of single-nucleotide sites across the 8.1 kb LTR element itself. A possible explanation is that both the RM9 and RM15 assemblies also contain a scaffold with an empty insertion site, indicating an intact version of the coding sequence spanned by the LTR insertion (represented in Fig. 4b by the partial matches on scaffolds 16719 and 07127, respectively). If the insertion is hemizygous, an uninterrupted homologous copy of the affected gene might mask or reduce the effect of the mutation. Given the degenerate tetraploid structure of bdelloid genomes, further homoeologous copies (i.e. derived from whole-genome duplication) might provide additional functional redundancy and help perform critical functions of interrupted genes.

Thus, these data contradict the idea that bdelloid TEs are inactive. All TE superfamilies in other bdelloids show a substantial fraction of copies at low-divergence, indicative of recent proliferation. Moreover, there are multiple cases of insertion-site polymorphism within species, and at least one case where a recent retroelement insertion into a protein-coding sequence seems likely to have potential fitness consequences.

### No evidence that cryptic recombination helps to limit the spread of LTR-Rs

Another possible explanation for the apparent discrepancy between bdelloid TE profiles and theory is that bdelloids in fact possess cryptic inter-individual recombination, either through undetected sex or some alternative form of gene transfer. We therefore tested for a signature of recombination among polymorphic LTR-R insertion sites within species. Under strict clonality, the pattern of presence and absence across LTR-R loci should be nested and compatible with only mutational gain and loss at each site. In contrast, in a sexual, outcrossing population, variation should be shuffled among loci. LTR-Rs provide a powerful test of these predictions because random insertion makes independent origins of the same LTR-R insertion site highly unlikely.

In every species with multiple samples, we found that variation in polymorphic TEs is perfectly nested, with a consistency index in parsimony reconstruction of 1. Furthermore, in the two species with multiple parsimony-informative characters, *R. socialis* and *R. magnacalcarata*, we found a significantly positive index of association of presence and absences among LTR-R insertion sites, as expected with clonal inheritance (S3 Table; S6 Fig). Approximate Bayesian Computation with simulations of expected patterns under varying frequencies of sexual reproduction showed that strictly clonal evolution could not be rejected (S7 Fig). While this test uses a restricted set of markers, and so should not be viewed as a test of recombination for the whole genome or species, it does support clonal inheritance of LTR-R loci and finds no evidence that inter-individual recombination helps to limit their spread. Nevertheless, local LTR-LTR recombination within genomes, leading to solo LTR formation, may act to bring the copy number down [44].

### No evidence for lower TE loads in desiccating bdelloids

The desiccation hypothesis posits that TE numbers may be kept in check via the action of DSB-repair processes during recovery from desiccation. Our study includes 11 nondesiccating bdelloid samples encompassing three obligately aquatic species (*R. macrura, R. magnacalcarata* and *R. socialis*), while the remaining samples were isolated from ephemeral ponds or moss and must undergo frequent cycles of desiccation and rehydration to survive. Contrary to the prediction that TE load should be reduced in desiccating species, there is little overall difference in TE proportions between desiccating and nondesiccating lineages (mean = 4.8% ± 1.3% *vs*. 5.0% ± 0.9% respectively). Broken down by TE superfamily, desiccating taxa have relatively more DNA transposons, simple, low complexity and unclassified repeats, and relatively fewer PLE, LTR and LINE-like retroelements (S3 Fig; S8a Fig), with the biggest differences seen between *Rotaria* lineages (S8b Fig). However, perhaps unsurprisingly given only two independent shifts in desiccation ability within our sample (see phylogeny in Fig 2a), results from a Bayesian mixed-effects modelling approach that controlled for phylogenetic relationships showed no significant correlations between desiccation ability and TE load, for either overall proportion or for any individual TE superfamily (S4 Table). For most TE superfamilies the strength of the phylogenetic signal (*λ*) was close to 1 (S9 Fig), consistent with a high fit of the data to the phylogeny under a Brownian motion model as would be expected if TE load evolves neutrally along branches of the phylogeny.

Two further comparisons of desiccating versus nondesiccating species among our wider sample of animals also present contrasting results. In chironomid midges, the desiccation-tolerant *P. vanderplanki* encodes substantially fewer TEs than its nondesiccating sister species *P. nubifer*, as predicted (0.8% and 2.2% respectively, although this rises to ∼11% in both species when all repeats are included). In tardigrades, however, the desiccation tolerant *Ramazzottius varieornatus* encodes a greater proportion of TEs than *Hypsibius exemplaris* (4.3% and 2.8%, respectively), which does not survive desiccation without extensive conditioning [81], although the trend is reversed when all repeats are included due to a large fraction of simple repeats in *H. exemplaris*. We therefore find no consistent evidence for the hypothesised link between anhydrobiosis and TE load in bdelloids or beyond. The overall effect of desiccation on TEs might be dual: while repair of a DSB within a TE via non-homologous end-joining would likely result in its inactivation (thus acting to reduce TE load), an efficient DSB repair system would enhance repair of DSBs that arise during transposition of cut-and-paste DNA TEs that leave a DSB behind upon excision (thus allowing an increased TE load).

### Bdelloids experience similar selective constraints on TEs as do other species

A fourth hypothesis is that the selective environment for TEs is different in bdelloids than in other animals, thereby shifting their TE profiles compared to simple theory. For instance, bdelloids might tolerate insertions within genes unusually well, owing to redundancy arising from tetraploidy or multiple gene copies [45,91,92]. First, we explored the genomic ‘environment’ of TE insertions and their potential effects on genome function. Differences in the location of TE insertions might reveal differential costs and benefits compared to other taxa. To do this, we first compiled a high-confidence list of class I retrotransposons by searching for proteins with significant similarity to the reverse transcriptase (RT) domain found in all retrotransposons. Phylogenies of the resulting alignments showed a diverse array of RTs in all species, most of them full-length (in terms of conserved subdomain presence) and clustered within the three primary retrotransposon superfamilies of PLEs, LTRs and LINEs (**Fig 5a**; S10 Fig; S5 Data). Many (but not all) clustered within families previously identified in *A. vaga*. The elevated LINE content in *D. carnosus* in comparison to other bdelloids is mostly due to the expanded Soliton clade and to the presence of CR1-Zenon and Tad/I/Outcast clades, the latter being characterized by the presence of the RNase H domain.

**Fig 5.**
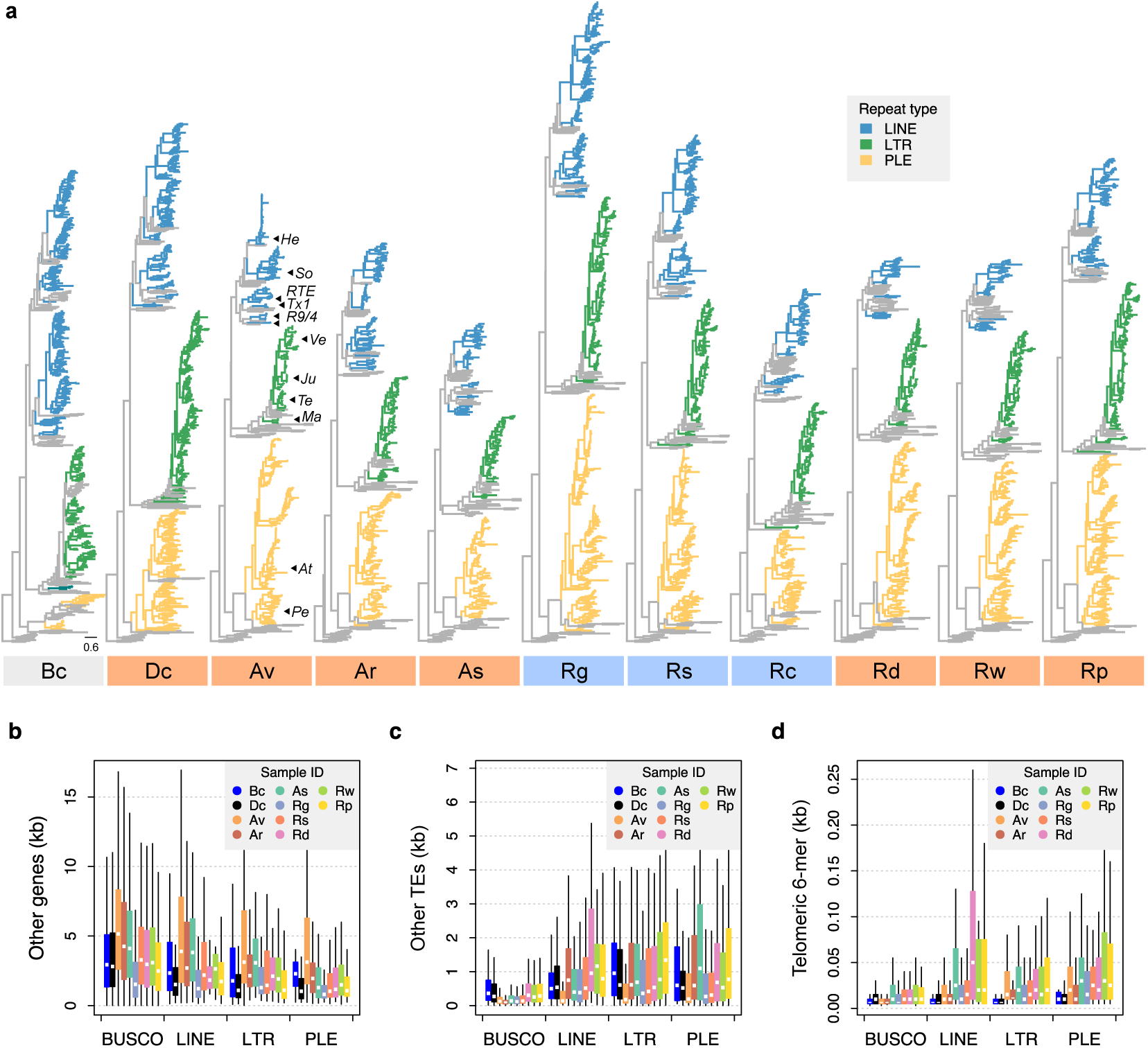
Phylogenetic diversity and genomic context of reverse transcriptase genes. **(a)** For each phylogeny, coloured branches represent identified rotifer-encoded RT copies and grey branches represent the core RT sequences from which the HMM was built (see Materials and methods and S9 Fig for core RT tree details). Colours indicated the major superfamilies. Previously characterised retrotransposons are indicated on the *A. vaga* tree (**He**, *Hebe*; **So**, *Soliton*; **RTE**, *RTE*; **Tx1**, *Tx1*; **R9/4**, *R9* and *R4*; **Ve**, *Vesta*; **Ju**, *Juno*; **Te**, *TelKA*; **Ma**, *Mag*; **At**, *Athena*; **Pe**, *Penelope*). All phylogenies are rooted on the branch separating the bacterial retrons. Scale bar represents 0.6 amino acid substitutions per site. Desiccating and nondesiccating species are indicated with orange and blue, as previously. Species codes: **Bc**, *B. calyciflorus* PSC1; **Dc**, *D. carnosus* DCAR706, **Av**, *A. vaga* Av2013; **Ar**, *A. ricciae* ARIC003; **As**, *A. steineri* ASTE805; **Rg**, *R. magnacalcarata* MAG3; **Rs**, *R. socialis* AK11; **Rc**, *R. macrura* Rc2018; **Rd**, *R. sordida* RSOR408; **Rw**, *R*. sp. ‘Silwood-1’ RSIL806; **Rp**, *R*. sp. ‘Silwood-2’ RPSE503. The genomic context in which RT genes reside is then described based on proximity to three other features: **(b)** ‘other’ genes (that do not overlap with any TE annotation), **(c)** other TEs, and **(d)** telomeric repeats (“TGTGGG”; that do not overlap with any coding region) as identified in *A. vaga*. For each plot, a 25 kb window is drawn around the focal RT gene and the total span (kb) of each feature within the window is counted, broken down per sample ID (coloured boxes) per TE superfamily (*X*-axis groups). Boxplots show the median (band), interquartile range (box) and minimum/maximum values (whiskers; outliers not plotted). The equivalent data for BUSCO genes (metazoan set) are also shown for comparison. The same representative samples are used in (b–d) as for (a).

We then characterised surrounding genome features for these TEs. In 50 kb windows surrounding each class I TE identified above, we counted the occurrence and span of three features of interest: other (non-TE) genes, other (non-focal) TEs, and the telomeric repeat “TGTGGG” (identified from *A. vaga* [58] and supported in other rotifers, see S2 Note). Relative to a set of core metazoan (BUSCO) genes, the regions surrounding PLE, LINE and LTR TEs all showed significant decreases in gene density, but significant increases in both TE and sub-telomeric repeat density (**Fig 5b–d**; S5 Table). In contrast, using a linear mixed-effects modelling approach (see Materials and methods), there were no significant differences between the monogonont *B. calyciflorus* and bdelloids, or between desiccating and nondesiccating bdelloid species (S5 Table). These results are consistent with previous findings that TEs are mainly confined to sub-telomeric regions of bdelloid genomes [56], a bias that is presumably due to either selection against insertions at or near functioning genes and/or strong specificity in TE insertion site. Thus, it appears that most TE insertions are costly in bdelloid rotifers, as in other taxa, and that selection leads to their concentration outside of gene-rich regions.

As a second source of selective constraints, we tested for evidence of selection against ectopic recombination (ER). ER is argued to be a major cost of TEs in sexual taxa, but its effects derive from chromosomal abnormalities during meiosis, which should be lacking in bdelloids. Because the rate of ER increases with both the number of elements and their length [12], the strength of selection is expected to be strongest against longer TEs at higher copy number [9,24,93]. Two testable predictions arise: first, that bdelloids should have longer TEs than sexual taxa (under the hypothesis that ER is absent in bdelloids because of a lack of meiosis), and second, that nondesiccating bdelloids should have longer TEs than desiccating bdelloids (under the hypothesis that ER may still occur when chromosomes pair during the repair of DSBs). However, comparisons of TE length distributions provide no evidence for these predictions— there was no significant difference between monogononts and bdelloids, or between desiccating and nondesiccating bdelloids (**Fig 6a**; S6 Table; S6 Data). Thus, while the precise estimation of TE lengths will no doubt improve with increasing assembly contiguity, our current data do not show any indication of changes in TE length linked to asexuality (when compared to monogononts) or desiccation ability within bdelloids. It is interesting to note that this finding also applies to the PLEs, which include the *Athena* elements that comprise the unusually large and complex *Terminon* elements found at bdelloid telomeres [59,94]. In this case, it is possible that selection may still act on individual *Athena* elements but does not affect *Terminons* per se because their structural diversity and genomic location does not make them targets of ER.

**Fig 6.**
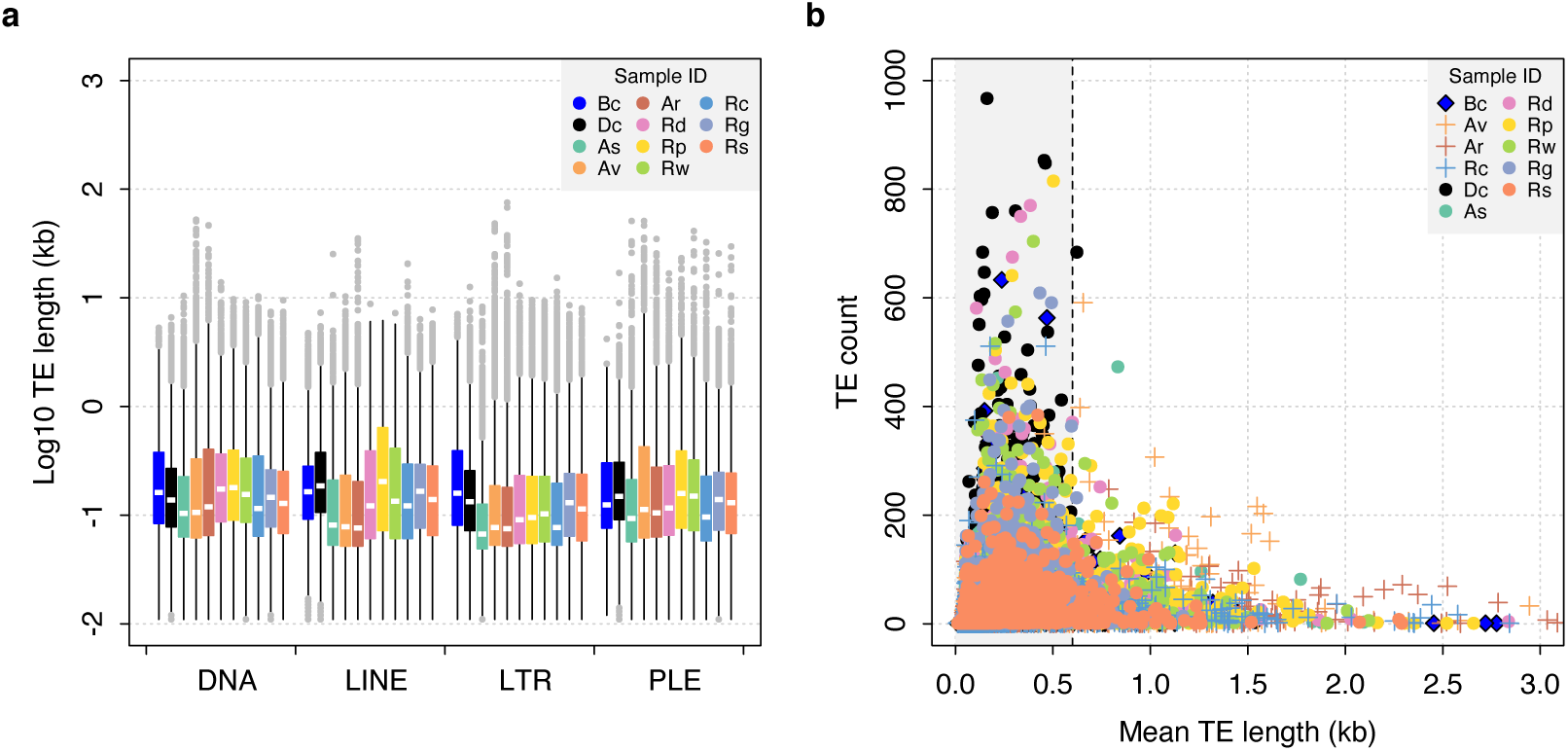
TE length dynamics. **(a)** Distribution of TE length for selected syndermatan samples decomposed into the major TE superfamilies (DNA transposons, LINE-like, LTR and PLE retrotransposons). Boxplots show the median (band), interquartile range (box) and minimum/maximum values (whiskers; outliers are shown in grey). Species codes: **Bc**, *B. calyciflorus* PSC1 (monogonont); **Dc**, *D. carnosus* DCAR706; **As**, *A. steineri* ASTE804; **Av**, *A. vaga* Av2013; **Ar**, *A. ricciae* Ar2018; **Rd**, *R. sordida* RSOR408; **Rp**, *R*. sp. ‘Silwood-2’ RPSE411; **Rw**, *R*. sp. ‘Silwood-1’ RSIL801 (desiccating bdelloids); **Rc**, *R. macrura* Rc2018; **Rg**, *R. magnacalcarata* MAG1; **Rs**, *R. socialis* AK11 (nondesiccating bdelloids). An equivalent plot including the acanthocephalan *P. laevis* is shown in S10 Fig. **(b)** Relationship between mean TE length per TE family (*X*-axis) and copy number (i.e., the number of TEs identified within each family; *Y*-axis). The same set of individuals are shown as for (a). A dashed line is drawn at 0.6 kb, given as the length threshold under which the rate of homologous ectopic recombination is negligible in mice.

A final prediction of selection against ER is that there should be a negative correlation between TE frequency and length, as is observed in *Drosophila* [24] and humans [93]. For both monogononts and bdelloids, the majority of identified TEs are short (< 1 kb), which are presumably partial matches or degraded copies. Nonetheless, we observe a sharp decline in copy number as mean TE length increases above ∼0.5 kb, and a distinct lack of longer elements at higher copy numbers (**Fig 6b**). In vertebrates, previous work has suggested a lower threshold of ∼0.6–1 kb under which ectopic recombination does not operate [93,95]. Thus, the observed patterns in rotifers are consistent with the hypothesis that longer elements above a certain length threshold are selected against more strongly due to the deleterious effects of ectopic recombination. This finding is contrary to a similar analysis performed on TEs in nematodes, which did not recover the expected relationship [18]. However, the pattern is the same in both desiccating and nondesiccating bdelloid representatives as well as the monogonont *B. calyciflorus*, and to some extent the acanthocephalan *P. laevis* (S11 Fig), suggesting that selection against longer TEs at higher copy number is a general feature in Syndermata, regardless of desiccation ability.

### Large expansion of TE silencing pathways in bdelloids

Having rejected a range of hypotheses to reconcile theory and observation of TE levels in bdelloid rotifers, we finally looked for expansions and/or diversifications in the molecular pathways that defend against TEs. We characterised copy number variation for three well-known gene families with direct roles in TE suppression via RNA interference (RNAi). (1) Argonaute proteins of both the Ago and Piwi subfamilies, the core effectors of RNAi gene-silencing that form complexes with various classes of small RNA [96,97]; (2) Dicer, an RNase III–family protein that cleaves double-stranded RNA (dsRNA) molecules from ‘target’ genes into shorter fragments that are subsequently incorporated into Argonaute complexes [98,99]; and (3) RNA-dependent RNA polymerase (RdRP), an RNA replicase that synthesises secondary small interfering RNAs (siRNAs) that amplify the silencing response [98,100].

Based on hidden Markov model (HMM) matches of key domains to the predicted proteomes of the Illumina maxhap assemblies (in which homologous copies are largely collapsed), we detect an average of 21.5 putative Argonaute copies, 3.9 Dicer copies and 37.3 RdRP copies in bdelloid genomes (**Fig 7a**; S4 Data). These expansions are substantially larger than previously uncovered from the diploid assembly of *A. vaga* (eight Dicers, 23 Ago/Piwi, 20 RdRP per diploid genome) [44], particularly after correcting for the assembly resolution (i.e. diploid vs haploid assembly; see Materials and methods), perhaps due to the increased sensitivity of the HMM-based approach, or to different degrees of pseudogenization and/or homolog collapse. Phylogenies of identified copies revealed a number of divergent clades, particularly in the Argonaute and RdRP families (**Fig 7b–d**; S12 Fig; S7 Data; S8 Data), that might indicate both expansion and diversification of these proteins. Even accounting for degenerate tetraploidy in bdelloids, the RdRP domain in particular is significantly expanded relative to monogononts and other eukaryotes (**Fig 7e–g**; S9 Data). This expansion is not found in the monogononts *B. calyciflorus* or *B. plicatilis* HYR1, nor is there evidence for it in the (unannotated) acanthocephalan genome (S4 Data), suggesting that the expansions seen in Ago and (particularly) RdRP are unique to bdelloids, including the basally divergent *D. carnosus*.

**Fig 7.**
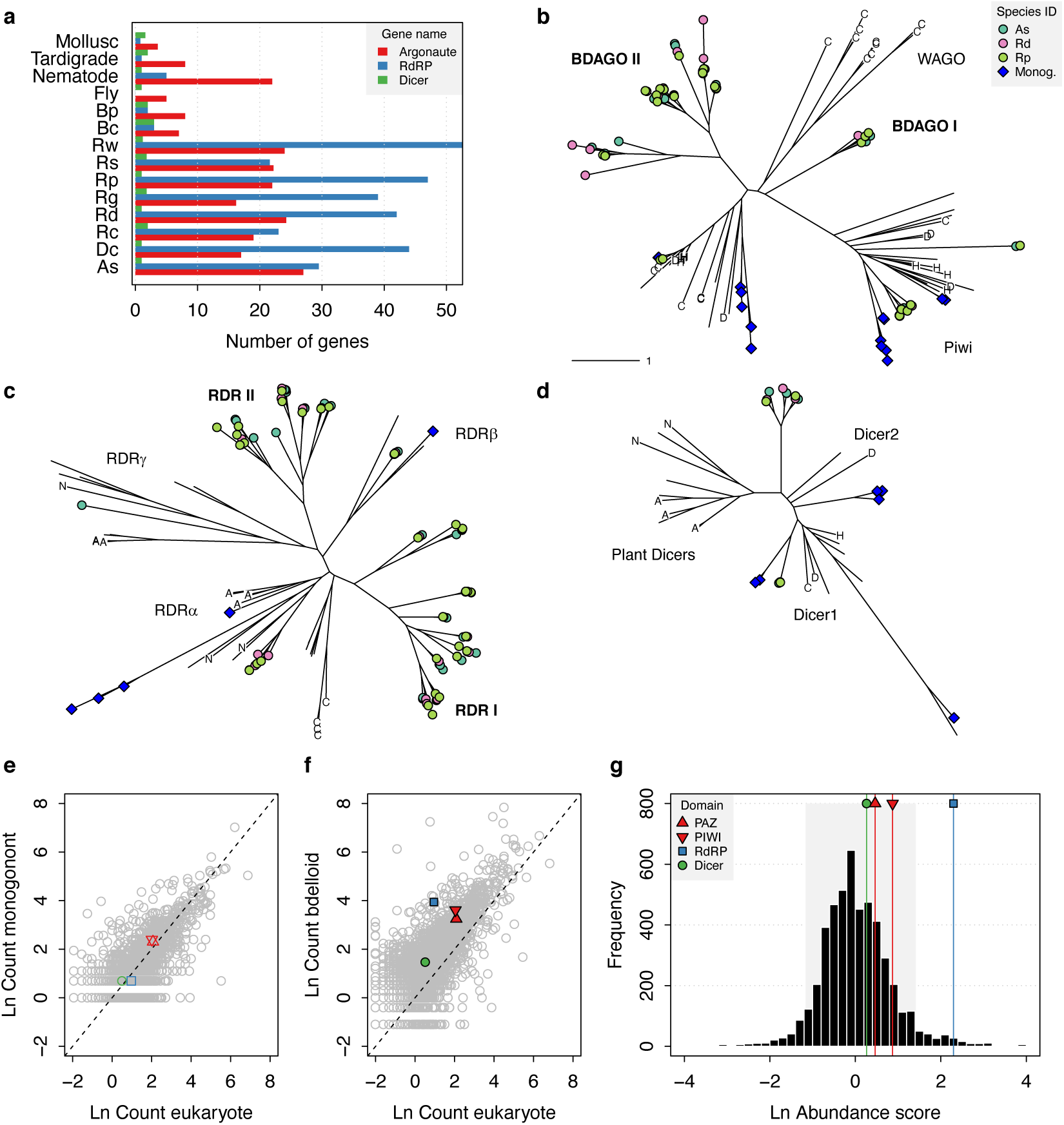
Expansion of TE silencing pathways in bdelloid rotifers. **(a)** Copy number variation for RNAi gene families Argonaute (Ago/Piwi, red), RNA-dependent RNA polymerase (RdRP, blue) and Dicer (green) in bdelloids compared to other protostome groups. Proteins are identified based on the presence of key identifying domains (see Materials and methods). Species codes for rotifers: **Bc**, *B. calyciflorus*; **Bp**, *B. plicatilis* HYR1; **Dc**, *D. carnosus*; **As**, *A. steineri*; **Rg**, *R. magnacalcarata*; **Rs**, *R. socialis*; **Rc**, *R. macrura*; **Rd**, *R. sordida*; **Rw**, *R*. sp. ‘Silwood-1’; **Rp**, *R*. sp. ‘Silwood-2’. Maximum likelihood unrooted phylogenies are then shown for **(b)** Argonaute, **(c)** RdRP and **(d)** Dicer gene copies identified in *A. steineri, R. sordida* and *R*. sp. ‘Silwood-1’ 10x haploid (‘pseudohap’) assemblies, aligned with orthologs from representative species from across the eukaryotes. Blue symbols indicate copies identified in the monogonont *B. plicatilis*, and letters on tips show selected reference species to aid visual orientation: ‘C’, *C. elegans*; ‘H’, human; ‘D’, *D. melanogaster*; ‘N’ *N. crassa*; ‘A’, *A. thaliana*. Some clade names are also shown where relevant; ‘WAGO’ indicates the worm-specific cluster of Ago genes in the Argonaute phylogeny. ‘BDAGO’ I and II and ‘RDR’ I and II indicate putative bdelloid-specific clades of Argonaute and RdRP proteins, respectively. **(e)** Comparative protein-domain abundance plot. Each point represents a Pfam domain ID, with (log) average abundance (i.e., count) in the reference eukaryote set shown on the *X*-axis and (log) abundance in the monogonont *B. plicatilis* on the *Y*-axis (see Materials and methods). The positions of the PAZ, PIWI (red up/down triangles), RdRP (blue square) and Dicer (green circle) domains are highlighted. Dashed line indicates the 1-1 relationship. **(f)** Equivalent plot for bdelloids, where the *Y*-axis shows the (log) average abundance for the *A. steineri, R. sordida* and *R*. sp. ‘Silwood-1’ 10x haploid assemblies. Note that the average abundance for all Pfam entries is shifted above the 1-1 line due to the ancient genome duplication in all bdelloids, such that many genes are found in double copy (i.e., homoeologs) even in a ‘haploid’ representation. **(g)** Comparative protein–domain abundance plot for bdelloids versus eukaryotes (see Materials and methods). Entries to the right of the distribution are overrepresented in bdelloids with respect to eukaryotes. The shaded area represents the 5% and 95% quantiles of the distribution, and the scores for the PAZ, PIWI, Dicer and RdRP domains are indicated (see legend).

The expansion of bdelloid-specific Ago genes (denoted ‘BDAGOs’ in **Fig 7**) is comparable to the ‘worm-specific’ Ago clade (WAGOs) found in nematodes [101–103]. Intriguingly, the majority of nematode species (although not *C. elegans*) have apparently lost their Piwi-like orthologue, the branch of the Argonaute family that is usually involved in TE suppression in most other animals [104], but instead mediate TE silencing using a Dicer/RdRP pathway [105]. The primary function of RdRP is to amplify RNAi responses via the production of small interfering RNAs (siRNAs) [98], but this family is found only rarely in animal genomes [100]. In *C. elegans*, RdRP-generated siRNAs (known as ‘22G-RNAs’) also play important roles in the recognition of ‘self’ versus ‘non-self’ RNA and multigenerational (i.e., inherited) epigenetic memory [103,106]. However, an analysis of TE variation across the nematode phylum found no effect of RNAi pathway differences (in terms of presence/absence of RNAi genes) among species, concluding that TE content is mediated primarily by the stochastic action of genetic drift [18].

Why do bdelloids possess such a marked expansion of gene silencing machinery? One explanation may be that it was required as part of the successful long-term transition to asexuality, if other mechanisms usually acting in sexual populations were no longer operating. Intriguingly, it has been shown in *A. vaga* that piwi-interacting small RNAs (piRNAs) target both TEs and putatively ‘foreign’ genes (i.e. non-metazoan genes gained via horizontal gene transfer [HGT]) [107], the latter of which are unusually frequent in bdelloid genomes [44,46,108–110]. Thus, it may be the case that bdelloids have an enhanced RNAi system to defend against invasion from horizontally transferred TEs, particularly if the level of exposure or rate of import is higher relative to other animals. Furthermore, the ability of some RNAi pathways to distinguish self from non-self may be needed to maintain genome integrity over longer timescales, if such mechanisms are operating. Further work on the precise functions of the divergent Ago and RdRP clades is required to explore these possibilities.

## Conclusions

We show that all bdelloids encode a rich diversity of TEs from both class I (retroelements) and class II (DNA transposons), and thus reject the idea that bdelloids are deficient or unusual in their TE content or diversity. Moreover, a substantial fraction of these elements has been active relatively recently within populations. This finding is at odds with the original predictions of population genetic theory for TEs in asexuals. One possible resolution is that theory is missing some component or assumption. It is possible that parameter space exists that permits intermediate levels of TEs in an asexual population, perhaps sustained by high rates of horizontal transfer, which there is evidence for in bdelloids. The HGT idea has some support from the increased prevalence of class II DNA transposons in bdelloids, given their greater propensity for horizontal transfer, but we found that the basally divergent lineage leading to *D. carnosus* had a profile more like the monogonont rotifers with regard to non-LTR retrotransposon abundance, despite sharing other features with the rest of bdelloids. Alternatively, some TEs might have been co-opted to provide beneficial functions, which is hypothesised to explain the unusually large and complex *Terminon* repeats. Other TEs may have evolved strong site-specificity to neutral genome regions to mitigate negative effects of transposition. This idea is supported by the preference shown for insertions into gene-poor regions that are probably at or near the telomeres, although it seems unlikely that the full complement of bdelloid TEs have accumulated in this way.

We also ask what forces may be acting to suppress TE activity in bdelloid populations. A major finding is that an abundance and diversity of RNAi gene silencing pathways, characterised by a large expansion of Argonaute and RdRP genes, appears to be a unique feature of bdelloid genomes. The precise origins and functions of these divergent Ago and RdRP clades are yet to be elucidated, but it seems likely that such an extended arsenal of TE defence genes offers enhanced protection against the deleterious effects of TE activity, particularly if bdelloid populations cannot keep TEs in check through sexual processes but are still exposed to new invasions via HGT. This enhanced RNAi system may then provide a more deterministic action against TEs, as opposed to stochastic forces (such as genetic drift) that may predominate in other animal groups and may potentially explain the greater uniformity of TE proportions in bdelloids relative to monogononts.

An alternative resolution is that the assumption of no recombination and strict clonality is not met in bdelloids. Previous work, for example, proposed that intra-individual recombination during the repair of DSBs caused by desiccation could provide a mechanism to keep TE numbers in check. We found no evidence here that overall TE loads were lower in desiccating species than nondesiccating species, a finding not limited to bdelloids, or for differences in activity or rate of turnover between them. It remains possible that an equivalent mechanism, such as mitotic recombination, or unusual DNA repair mechanisms operating in nondesiccating species as well, could still act to limit TE proliferation and facilitate elimination of inserted copies, maintaining an equilibrium between TE spread and removal by excision and selection. Finally, there could be some hidden mechanism of inter-individual recombination that facilitates TE removal. We found no evidence for its action here, but further work is needed for a final answer on the conundrum of bdelloid asexuality.

## Materials and methods

### Rotifer sampling and culture

For most samples, individual rotifers were collected from permanent and temporary freshwater habitats around Imperial College London’s Silwood Park campus, Ascot, UK between May 2015 and February 2019. Three samples (*R. magnacalcarata* RM9 and RM15, and *R. socialis* RS1) were collected from a freshwater spring in Fontaneto d’Agogna, Italy in 2016 (see S1 Data for details). Although we focused on the genera *Adineta* and *Rotaria*, we also included two individuals from the desiccation-tolerant species *Didymodactylos carnosus*. Preliminary phylogenetic data had identified this as a distant outgroup to the focal genera, useful in rooting phylogenetic trees and as a further independent datapoint to test the generality of conclusions about bdelloids. A total of 26 samples were submitted for single-individual, whole genome sequencing; for these, DNA was extracted using either a Chelex preparation (Bio-Rad InstaGene Matrix) or a QIAamp DNA Micro Kit (Qiagen), and whole-genome amplified using a REPLI-g Single Cell kit (Qiagen) before sequencing on either Illumina NextSeq500 at the Department of Biochemistry, University of Cambridge (Cambridge, UK), or Illumina HiSeq X at Edinburgh Genomics, University of Edinburgh (Edinburgh, UK). For *A. ricciae* ARIC003, DNA was extracted from ∼200 animals descended from a single individual before whole-genome amplification. For *B. calyciflorus* PSC1, individuals for DNA extractions were derived from an individual isolate from a laboratory stock population previously isolated from field-collected resting eggs [111]. DNA was extracted from ∼5000 starved individuals using a phenol-chloroform protocol and sequenced on the Illumina NextSeq500 at the Max Planck Institute for Evolutionary Biology. Three 10x Genomics Chromium ‘linked reads’ libraries were generated for *A. steineri, Rotaria* sp. ‘Silwood-1’ and *R. sordida*; for these, high molecular weight DNA was extracted from thousands of animals reared clonally from a single wild-caught animal, without whole-genome amplification, using the Chromium Demonstrated Protocol “HMW gDNA Extraction from Single Insects” (https://support.10xgenomics.com/permalink/7HBJeZucc80CwkMAmA4oQ2). Linked-read libraries were constructed at the Centre for Genomics Research, Liverpool, UK, before sequencing on the HiSeq X at Edinburgh Genomics. Further details on rotifer sampling, DNA extraction and sequencing are provided in S1 Data.

### Biological replicates

To check the repeatability of the whole-genome amplification (WGA), sequencing, assembly and analysis pipelines, we included several samples that were either biological replicates of the same rotifer clone, or where high-quality genomes were available for the same clone from unamplified source material. Specifically, for *Rotaria* sp. ‘Silwood-2’ we isolated two consecutive offspring from the same wild-caught mother and conducted WGA, sequencing, assembly and analysis for these sisters independently (as Rp_RPSE411 and Rp_RPSE503). From the same clonal laboratory line of *Rotaria* sp. ‘Silwood-1’ that was used for 10x Genomics DNA preparation, we isolated two more individuals and processed each independently using the WGA workflow (as Rw_RSIL801 and Rw_RSIL802). Finally, we applied the WGA method to DNA from *A. ricciae*, for which a previous assembly was available from unamplified DNA [46] on the same clonal culture and included this replicate in downstream analyses alongside the earlier reference assembly.

### Data filtering and genome assembly

We generated two assembly versions for each of the single-individual rotifer samples. The ‘reference’ assemblies were scaffolded and polished to result in haploid assemblies with improved contiguity. The ‘maximum haplotig’ (‘maxhap’) assemblies instead retained highly similar contigs that might otherwise be removed during assembly polishing. Our pipeline is outlined as follows.

For the Illumina libraries, raw sequence data were filtered for low quality bases and adapter sequence using BBTools v38.73 ‘bbduk’ (https://sourceforge.net/projects/bbmap/), and error corrected using BBTools ‘tadpole’. Data quality was inspected manually using FastQC v0.11.5 [112] aided by MultiQC [113] visualisation. For the *A. steineri, R*. sp. ‘Silwood-1’ and *R. sordida* linked-read libraries, data were assembled into haploid (‘pseudohap’) and diploid (‘megabubbles’) genome representations using the 10x Genomics proprietary software Supernova v2.1.1 [114] and further scaffolded with ARKS v1.0.4 [115]. All raw sequencing data are deposited in the relevant International Nucleotide Sequence Database Collaboration (INSDC) databases under the Study ID PRJEB39843 (see S1 Data for run accessions and counts for raw and filtered data).

For the single-individual samples, an initial assembly was generated using SPAdes v3.13.0 [116] with default settings. Contaminating reads from non–target organisms, identified based on aberrant GC content, read coverage and/or taxonomic annotation, were then identified and removed using BlobTools v1.1.1 [117,118]. For *R. magnacalcarata* and *R. socialis* samples, resultant haplotigs were then collapsed using Redundans [119] with default settings before scaffolding and gap filling with SSPACE v3.0 and GapCloser v1.12 respectively [120,121]. For *A. steineri, R*. sp. ‘Silwood-1’ and *R. sordida* single-individual samples, the scaffolding step was performed with RaGOO v1.1 [122,123], using the matching 10x Genomics ‘pseudohap’ assembly as a reference (contigs from *R*. sp. ‘Silwood-2’ were scaffolded using the *R*. sp. ‘Silwood-1’ 10x reference), specifying the ‘-C’ parameter to prevent concatenation of unaligned contigs. Scaffolded assemblies were subjected to further rounds of BlobTools to remove any additional sequences derived from non-target organisms. These assemblies were designated the reference set described above.

For the maxhap assemblies, filtered fastq files were first generated by mapping the original (trimmed and error-corrected) sequencing reads to each reference genome, using the ‘outm=filtered_R#.fq’ functionality of BBTools ‘bbmap’, and then reassembled with SPAdes, increasing the final kmer value to 121. Assembly metrics were summarised using ‘calN50.js’ (https://github.com/lh3/calN50), which reports the ‘expected scaffold size’ (AU) as an alternative metric of assembly contiguity that is less biased than N50 (defined as the area under the cumulative genome span versus contig length graph, equivalent to the expected scaffold size for a randomly chosen assembly location [124]). Gene-completeness scores for core eukaryotic (*n* = 303) and metazoan (*n* = 978) genes were calculated for all assemblies using BUSCO v3.0.2 [125] with default settings. Reference and maxhap assemblies for *B. calyciflorus* PSC1 and *D. carnosus* are the same, due to a lack of appropriate data for scaffolding.

### Gene prediction

Gene prediction was performed on reference assemblies using one of three approaches, depending on the availability of RNA-seq data. For *B. calyciflorus, A. ricciae*, and all *R. magnacalcarata, R. socialis*, and *R. sordida* assemblies, published RNA-seq data [109,110,126] were downloaded from NCBI Sequence Read Archive (SRA), quality-trimmed using BBTools ‘bbduk’ with default settings and aligned to the genomic scaffolds using STAR v2.7.3a [127] with the option ‘--twoPassMode Basic’. Aligned BAM files were then provided to BRAKER v2.1.2 [128–132] with default settings for gene prediction. For *A. steineri, R*. sp. ‘Silwood-1’ and *R*. sp. ‘Silwood-2’ assemblies, RNA-seq data from a related species (*A. ricciae* and *R. magnacalcarata* respectively) were used instead, aligned using BBTools ‘bbmap’ with the options ‘maxindel=200k minid=0.5’, before gene prediction with BRAKER as above. Finally, for the distantly related *D. carnosus*, BRAKER was run using gene-model parameters estimated from BUSCO analysis of the genomic scaffolds. The quality of predicted proteins was assessed using BUSCO in protein mode. Intragenomic divergence between homologous gene copies and collinearity was calculated as for Nowell *et al*. (2018) and described in S1 Note. Genome assemblies and gene predictions were converted to EMBL format using EMBLmyGFF3 v2 [133], and are deposited at DDBJ/ENA/GenBank under the Study ID PRJEB39843 (see **Table 1** and S1 Table for individual GenBank accessions).

### Rotifer phylogeny

Evolutionary relationships among new genomes and published genomes of rotifers were determined using a core-genome phylogenomics approach based on the BUSCO eukaryotic gene set. For genomes from species with very high intragenomic homologous divergence (*A. ricciae* and *A. vaga*), redundancy was removed by selecting the copy with the highest BUSCO score for all BUSCO genes with multiple copies, using the script BUSCO_collapse_multicopy.pl’ (https://github.com/reubwn/scripts). One-to-one co-orthologs found in at least 95% of the samples were then identified using the script ‘BUSCO_phylogenomics.py’ (https://github.com/jamiemcg/BUSCO_phylogenomics). Protein sequences were aligned using Clustalo [134] and concatenated in Geneious R9 [135]. The full alignment was checked by eye and sections with ambiguous alignment within the bdelloid clade were removed across all sequences to avoid aligning potential paralogs or homoeologs. Translation errors arising from annotation issues in specific bdelloid genomes were identified by obvious mismatches to the consensus of closely related genomes, and the affected residues were deleted in the affected genome only. Potential alignment issues within the monogonont clade were less obvious owing to the substantial genetic divergence from bdelloids and the smaller number of genomes and replicates, so corrections were less stringent. A maximum-likelihood phylogeny was then estimated using IQ-TREE v1.6.12, with automatic model selection (VT+F+I+G4) [136,137]. Branching support was assessed using SH-aLRT and ultrafast bootstrap sampling (‘-alrt 1000 -bb 5000’) [138,139].

### Repeat annotation and TE dynamics

TEs and other repeats were identified using the RepeatModeler and RepeatMasker pipelines. For each sample, a *de novo* repeat library was generated directly from the assembled nucleotides using RepeatModeler2 [140] and combined with a database of 12,662 protostome repeats from Repbase v23.08 [141] and 278 additional TEs manually curated from the *A. vaga* genome [44]. Repeats and TEs were then detected and classified using RepeatMasker v4.1.0 [84], and resultant outputs were post-processed using the ‘One code to find them all’ Perl script [142]. The breakdown of TE superfamilies in the final database was 4,145 DNA transposons (including 300 rolling circles), 5,523 LTRs, 2,583 LINEs (including SINEs), 227 PLEs, and 165 simple or low-complexity repeats. TE content (expressed as a proportion of genome size) was mapped onto the phylogeny using ‘contMap’ in the Phytools v0.6-99 package in R v3.6.0 [143,144]. There is no module for the detection of class II MITEs in RepeatMasker; for these, the separate program Generic Repeat Finder (GRF) was run using default parameters. TE dynamics were investigated by constructing Kimura 2-parameter divergence [85] landscapes using the utility scripts in the RepeatMasker package and plotted using custom scripts (see below). Selected assemblies were also submitted to the REPET v2.5 ‘TEdenovo’ [145,146] TE detection and annotation pipeline with default parameters, for comparison. In addition, for *D. carnosus* and *R. sordida* (using 10x Genomics) reference assemblies, we increased the parameter ‘minNbSeqPerGroup’ from 3 to 5 to evaluate contribution from tetraploid genes, which was judged to be negligible. Although REPET*denovo* TE consensus sequences are automatically classified using Wicker’s TE classification [147], RepeatMasker was additionally applied for further TE classification, detection and landscape divergence plot building.

The presence or absence of specific LTR retrotransposon (LTR-R) insertions in our population data was inferred using a read-mapping approach. Specifically, the presence of a given insertion was scored based on the alignment score of the ‘best’ read that mapped continuously and contiguously across the LTR-genome boundary. First, full-length LTR-Rs (i.e. those with annotated 5’ and 3’ LTR regions) were identified from each reference assembly using LTR_retriever v2.8 [148]. Three filters were then applied to remove false positives. Candidates that showed an overlap with a predicted gene in the 5’ or 3’ LTR itself or an ‘N’ base within 150 bases upstream or downstream of its genomic location that might indicate local mis-assembly were removed. Candidates also required supporting evidence of LTR homology from a separate RepeatMasker annotation of the reference assembly. For each remaining LTR-R, a library of ‘LTR-tags’ was then generated by extracting a 100 bp sequence that spanned 50 bases into the genomic (i.e., non-TE) region of the insertion site from both the 5’ and 3’ terminal repeated regions. Thus, each pair of ‘LTR-tags’ represents an insertion of a particular LTR into a specific location in the focal genome, and a score is calculated based on the alignment information contained in the CIGAR string of the ‘best’ read (i.e. with the highest number of alignment matches) from the SAM mapping file: *Si* = ((*MLi* - *XLi*) + (*MRi* - *XRi*))/200, where *MLi* is the number of alignment matches for the left-hand tag for LTR *i*, penalised by the number of mismatches *XLi*, with equivalent scoring for the right-hand tag. Since tag length is 100 bases, the maximum score for a perfect alignment is 200, or 1 after normalisation. The number of mapped reads is also recorded to provide an estimate of coverage (but note that *Si* is taken from the best read only). Sequencing reads from all single-individual rotifer samples were aligned to the filtered LTR-tag set using BBTools ‘bbmap’ with the parameters ‘minid=0.5 local=t’ and scored using the above system. Because orthologous LTR-Rs may be identified from searches started in different genomes, we identified these cases by reconstructing the phylogeny of the LTR-tags and any with pairwise sequence divergence less than 0.1 were collapsed to yield a condensed final matrix.

The LTR-tag case-study in Fig. 4b was selected for closer investigation in the draft assemblies after consideration of several examples, because it illustrates variability for an element insertion site within a species and indicates that Class I TEs can insert in coding regions, with potential fitness consequences. The LTR-tags were mapped to the RM15 draft assembly using Geneious Prime v2020.1.2 [135], and were found to match an element annotated by LTR_retriever, containing four predicted genes. In the RM9 draft assembly, only the left-hand tag was mapped, as the scaffold ended before the inserted element was fully assembled. For the same reason, the element in RM9 had not been annotated as such by LTR_retriever, but the sequence is nearly identical (99.7%) to the insertion in RM15 along its aligned length (except that the annotations predicted three element-associated genes rather than four). The scaffolds were trimmed to the focal gene and aligned, and the region was used as a BLASTn query against local databases for two other *R. magnacalcarata* reference genomes where the LTR-tag was absent: MAG1 and Rg2018. In each case, this provided the location of a closely similar but uninterrupted copy of the focal gene, although the annotations of the gene’s structure differed slightly among genomes. These scaffolds were trimmed and aligned against the copies from RM9 and RM15, using the Geneious alignment tool with default settings, except that the gap extension penalty was reduced from 3 to 0.2 to enable the algorithm to handle the element insertion. Local features were manually reannotated to illustrate the interpretation provided in the text. To investigate the potential function of the interrupted gene, the copy from MAG1 (g37061) was translated and used as a BLASTp query against the NCBI RefSeq Protein Database [149]. A region of approximately 1000 residues was found to have weak similarity (∼25% pairwise identity) to proteins annotated as midasins, from a range of eukaryotes. As a final step, the intact gene from MAG1 was used as a BLASTn query against the full draft genomes of RM9 and RM15, which revealed a separate scaffold in each case, containing a partial copy of the gene in which the coding sequence was intact across the junction spanned by the LTR-tag, and the element insertion was absent.

### Recombination analyses

We tested sexual versus clonal patterns of variation in LTR presence and absences. First, we calculated consistency indices (CI) with parsimony reconstruction of the binary matrix. LTR-tags with scores > 0.875 were coded as present (i.e. no more than half of the genome context or LTR region from both left and right LTR-tags was missing) and < 0.875 coded as absent (alternative thresholds led to the same qualitative results). A CI = 1 indicates perfect nesting with no homoplasy, whereas a score less than 1 is expected if variation is shuffled among loci and not tree-like. Next, we calculated the index of association and ran permutations to test for significant linkage disequilibrium of the LTR-tag data relative to a null model of random shuffling (expected in a fully outcrossing sexual population). We used the modified index of association by Agapow and Burt (2001) [150] that corrects for an effect of the number of loci on the index, and ran permutations using the ‘ia’ function in the Poppr v2.8.5 library [151] in R. Data were coded as diploid and codominant presence/absence data (because of the lack of diploid assemblies in the population-level data). Finally, for *R. magnacalcarata* and *R. socialis* we ran simulations with the FacSexCoalescent simulator of Hartfield et al. (2018) [152] to generate 50000 datasets with the same number of individuals and sampled binary loci as observed, but with frequencies of sexual versus asexual reproduction within the populations varying from 10^−7^ (i.e. negligible) to 1 (i.e. obligate sexual). We estimate the posterior distribution of the frequency of sex for our observed samples using Approximate Bayesian Computation on the simulated datasets implemented in the abc package in R.

### Reverse transcriptase survey

A hidden Markov model (HMM) approach was used to survey the predicted rotifer proteomes for proteins encoding the reverse transcriptase (RT) domain (Pfam ID PF00078). First, a HMM was constructed from an alignment of 51 RT domains from across the tree of life [57], supplemented with 67 bdelloid-specific retroelements [44,57,58,61,63,64] (S9 Fig). Alternative transcripts were first removed from predicted proteomes and proteins with a significant match (*E*-value ≤ 1e-5) were identified and inserted into the core RT alignment using HMMER v3.2.1 ‘hmmsearch’ and ‘hmmalign’, respectively (http://hmmer.org/). Maximum likelihood phylogenies were then constructed using IQ-TREE as above, specifying the root of the phylogeny to be on the branch leading to the bacterial retrons [57]. Trees were manipulated using FigTree v1.4.4 (https://github.com/rambaut/figtree), colouring the identified RT-encoding rotifer proteins based on their phylogenetic position. The span accounted for by genome ‘features’ (other genes, other TEs and telomeric repeats) in a 25 kb window around each identified RT-containing protein was summarised using BEDTools v2.29.2 ‘intersect’ and ‘groupby’ [153]. Other genes were counted as predicted coding regions that did not overlap with any TE annotation. Genomic locations of the telomeric hexamer ‘TGTGGG’ [58] were identified using EMBOSS ‘fuzznuc’ [154], excluding any hexamer that overlapped with a predicted CDS. Note that the telomeric repeat for *Brachionus* is not known, but the sequence above was among the most frequent G-rich hexamers identified in the PSC1 genome (see S2 Note).

### TE silencing machinery survey

A similar HMM based approach was used to characterise gene copy evolution of three key pathways involved in RNAi gene-silencing. Putative Argonaute proteins were identified based on the presence of both the PAZ and PIWI domains (Pfam IDs PF02170 and PF02171 respectively), putative Dicer proteins were identified based on the presence of both PAZ and Dicer (PF03368) domains, and putative RdRP proteins were identified based on the presence of the RdRP domain (PF05183). Stockholm files were downloaded from Pfam [155] and aligned to the proteomes using HMMER (*E*-value ≤ 1e-5) as above. Reference proteomes from a selection of eukaryotic species to represent the diversity and distribution of Argonaute, Dicer and RdRP proteins were downloaded (June 2020) from UniProt and subjected to the same procedure: *Arabidopsis thaliana* (UP000006548), *Oryza sativa* (UP000007015), *Neurospora crassa* (UP000001805), *Schizosaccharomyces pombe* (UP000002485), *Laccaria bicolor* (UP000001194), *Dictyostelium discoideum* (UP000002195), *D. melanogaster* (UP000000803), *C. elegans* (UP000001940), *H. exemplaris* (UP000192578), *H. robusta* (UP000015101), *L. gigantea* (UP000030746), *S. haematobium* (UP000054474), *B. plicatilis* (UP000276133), *Branchiostoma floridae* (UP000001554) and *Homo sapiens* UP000005640. Proteins were aligned using either ‘hmmalign’ from the HMMER package or Clustalo, and ML phylogenies were constructed using IQ-TREE as above.

Comparative protein–domain abundance plots were constructed using counts of Pfam entries parsed directly from InterProScan5 [156] annotation of predicted proteomes. The ‘abundance score’ was computed as the (log) ratio of domain counts in bdelloids divided by the domain counts in eukaryotes, corrected for inflation in bdelloids due to the ancient whole-genome duplication by dividing the former by two. This correction is likely to be conservative, since many loci have lost one branch of the ancient duplication (i.e. tetraploidy is degenerate). To check that the putative RdRP expansion was indeed eukaryotic in origin, rather than viral, the HMMs for four viral RdRP families (PF00680, PF00978, PF00998 and PF02123) were downloaded from Pfam and submitted to the same search protocol, with zero hits to bdelloid proteomes recorded.

### Statistical analyses

To assess differences in TE content between desiccating and non-desiccating rotifer species, we ran Bayesian linear mixed-effects models of TE content (as a percentage of genome span) including desiccation as a two-level fixed factor and sample ID as a random intercept term. The BUSCO gene phylogeny was used to account for non-independence among species. A separate model was run for each TE superfamily (including ‘unclassified’ TEs) as well as for all TEs combined. Inverse-Wishart priors were used for the random and residual variances, and models were run for 42,0000 iterations with a burn-in of 20,000 and a thinning interval of 200. This resulted in 2,000 stored samples of the posterior with minimal autocorrelation in all cases (< 0.2) [157]. Models were run using the MCMCglmm v2.29 [158] package in R. The phylogenetic signal, defined as the proportion of the total variance in TE content attributable to the phylogeny [159], was estimated from the MCMCglmm model output using the formula: λ = σ*P*^2^/(σ*P*^2^ + σ*R*^2^).

The density of genomic features surrounding BUSCO genes versus RT-containing genes were compared using linear mixed effects models with species as a random effect and gene class (BUSCO, PLE, LINE or LTR) as a fixed effect, run using lme4 [160] v1.1-21 in R. TE length distributions were compared using the same approach, with species as a random effect and binary factors specifying monogononts versus bdelloid, and desiccating versus nondesiccating species.

### Code availability

All TE analysis scripts used in this study are available at https://github.com/reubwn/te-evolution.

## Acknowledgements

Genome sequencing was performed by the UK Natural Environment Research Council (NERC) Biomolecular Analysis Facility at the Centre for Genomic Research (CGR) at the University of Liverpool (NBAF-Liverpool) and the DNA Sequencing Facility in the Biochemistry Department at the University of Cambridge. The authors wish to thank the following: Christiane Hertz-Fowler, Pia Koldkjær and John Kenny (CGR), Shilo Dickens and Nataliya Scott (Cambridge). Matthew Arno and Colin Sharp (Edinburgh), and Steven Van Belleghem (University of Puerto Rico) for support with the planning and execution of various aspects of genome sequencing and/or assembly, Tom Smith and Anita Kristiansen for rotifer sampling, and Mike Tristem for helpful discussions on detecting LTR polymorphisms.

