## Supplementary Information for "Evolutionary dynamics of transposable elements in bdelloid rotifers"

### Table of contents

| | |
|---|---|
| **S1 Data** | Sample information and data counts (XLSX) |
| **S2 Data** | Repeat content data for rotifers and other animals (XLSX) |
| **S3 Data** | Reference and maxhap assembly RepeatMasker output files (TGZ) |
| **S4 Data** | LTR-tag fasta file and mapping data (TGZ) |
| **S5 Data** | Reverse-transcriptase alignments and phylogenies (TGZ) |
| **S6 Data** | TE length data (TGZ) |
| **S7 Data** | Gene family expansion data (XLSX) |
| **S8 Data** | Argonaute, Dicer and RdRP alignments and phylogenies (TGZ) |
| **S9 Data** | Pfam counts (GZ) |

**Reviewer link to supplementary data:** (1.7 Gb TGZ archive)
https://www.dropbox.com/s/j6oam4ycm8m3gky/Supplementary.v0.8.7.tgz?dl=0

| | |
|---|---|
| **S1 Fig** | Genome characteristics for bdelloid samples |
| **S2 Fig** | Repeat content and diversity in 42 rotifer and 17 protostome animal genomes |
| **S3 Fig** | TE content mapped as continuous trait onto rotifer phylogeny |
| **S4 Fig** | TE divergence landscapes for individual rotifer and acanthocephalan genomes |
| **S5 Fig** | TE divergence landscapes for selected species constructed using REPET. |
| **S6 Fig** | LTR-R presence/absence matrix for *R. magnacalcarata* and *R. socialis* |
| **S7 Fig** | The median and 95% Highest Posterior Density interval of the frequency of sexual recombination affecting the presence/absence of LTR polymorphisms in *R. magnacalcarata* and *R. socialis* |
| **S8 Fig** | Repeat content in desiccating and nondesiccating bdelloids |
| **S9 Fig** | Phylogenetic signal in the effect of desiccation on TE content |
| **S10 Fig** | Maximum likelihood phylogeny for diverse reverse transcriptase domains from across the tree of life |
| **S11 Fig** | Distribution of TE lengths including the acanthocephalan *P. laevis* |
| **S12 Fig** | Argonaute, RdRP and Dicer phylogeny details |

| | |
|---|---|
| **S1 Note** | Intragenomic divergence and collinearity |
| **S2 Note** | Identification of putative telomeric repeats in rotifer genomes |

| | |
|---|---|
| **S1 Table** | Assembly statistics for 30 maximum-haplotig assemblies presented in this study |
| **S2 Table** | Genome properties of 16 protostome animals included in the study |
| **S3 Table** | Analysis of recombination in LTR-tag presence/absence data |
| **S4 Table** | Posterior mean and 95% credible intervals for the effect of desiccation ability on TE load |
| **S5 Table** | Fixed and random effects estimates from a linear mixed-effects model for the genomic surroundings of class I TEs |
| **S6 Table** | Fixed and random effects estimates from a linear mixed-effects model of TE lengths |

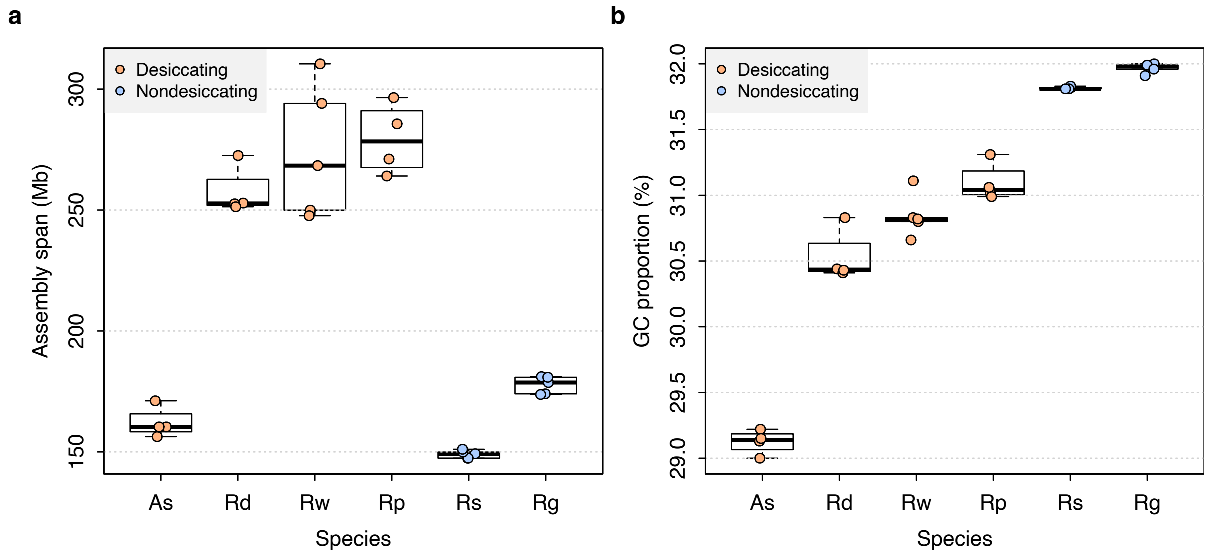

**S1 Fig. Genome characteristics for bdelloid samples.** Boxplots are shown for (**a**) genome assembly span and (**b**) GC proportion (percent G + C nucleotides). Data shown for species with $n \geq 3$ individuals. Boxplots show the median (band), interquartile range (box) and minimum/maximum values (whiskers). Underlying data are shown as jittered points. Species codes: **As**, *A. steineri*; **Rd**, *R. sordida*; **Rw**, *R.* sp. 'Silwood-1'; **Rp**, *R.* sp. 'Silwood-2'; **Rs**, *R. socialis*; **Rg**, *R. magnacalcarata*.

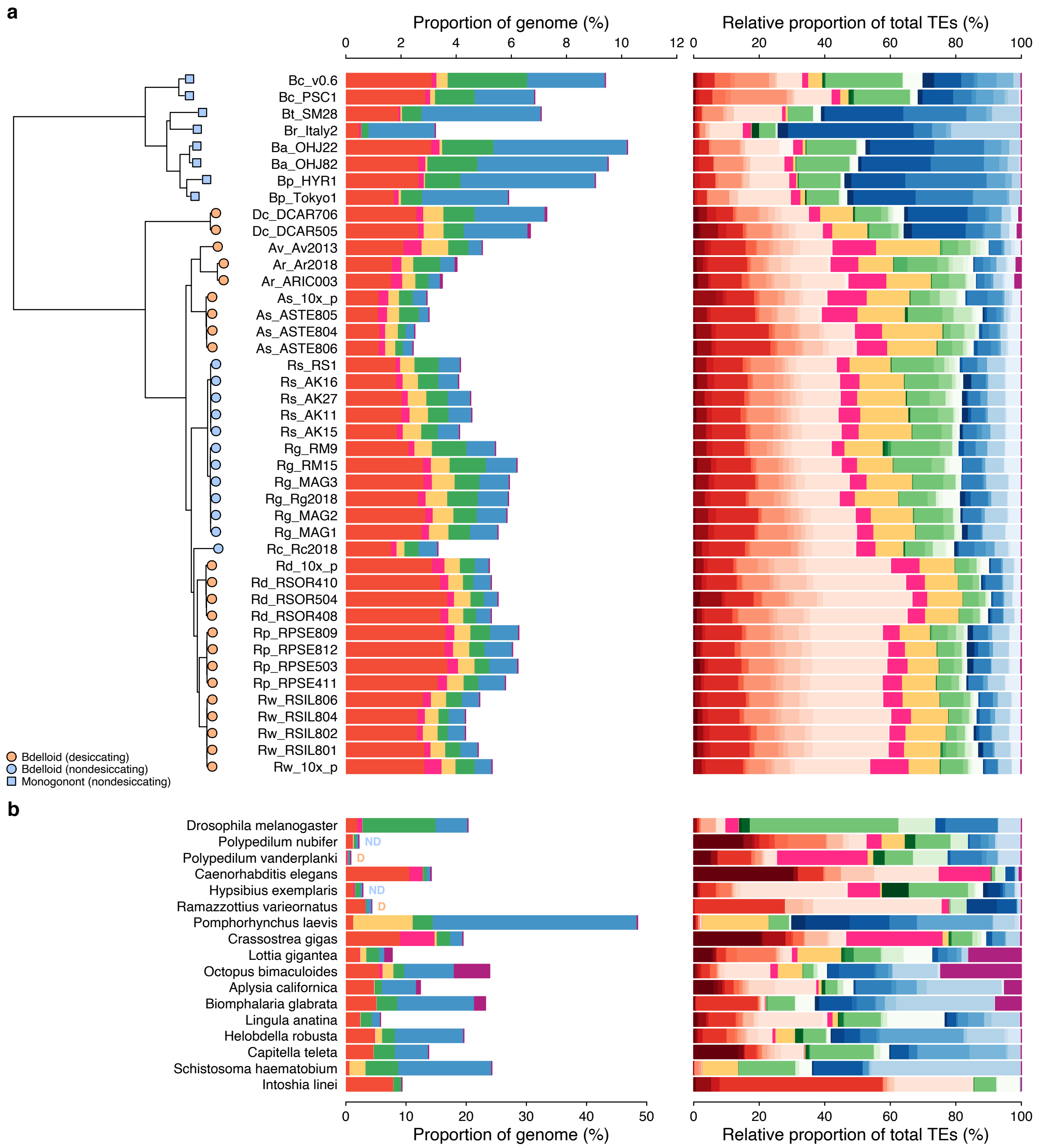

**S2 Fig. Repeat content and diversity in 42 rotifer and 17 protostome animal genomes.** Plot is the same as in main text but data for satellite, simple, low complexity and unclassified repeats are omitted.

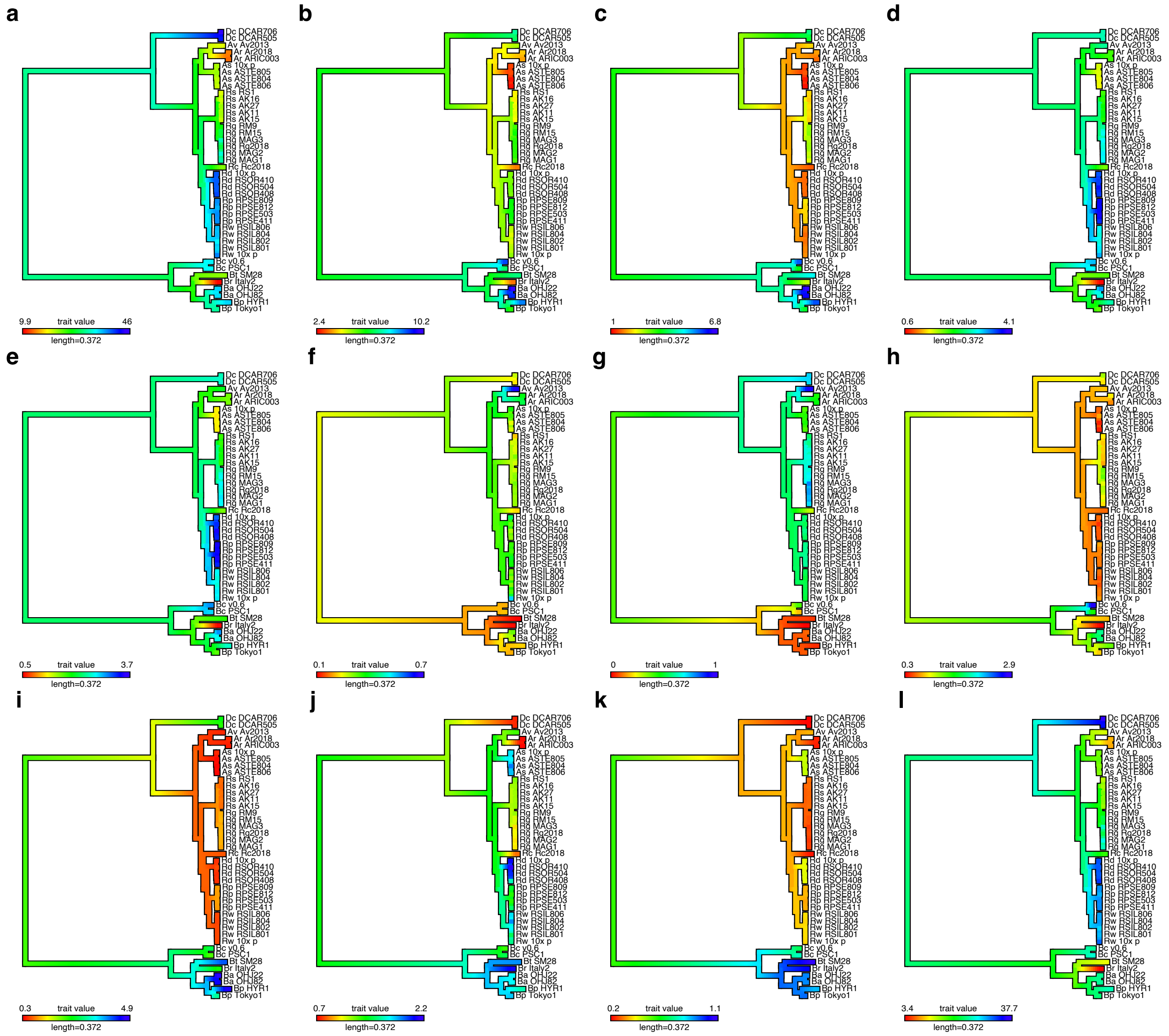

**S3 Fig. TE content mapped as continuous trait onto rotifer phylogeny.** Total repeat content is decomposed into **(a)** all repeats of any kind, **(b)** known (i.e., that are classified into one of the major Class I or II superfamilies) TEs only, **(c)** all class I retroelements, **(d)** all class II transposons, **(e)** DNA transposons **(f)** rolling circles, **(g)** PLEs, **(h)** LTRs, **(i)** LINEs **(j)** simple repeats, **(k)** low complexity repeats and **(l)** unclassified (unknown) repeats. Repeat content is mapped as a percentage of genome span, with blue indicating a relative increase and red indicating a relative decrease in TE type along a given branch of the phylogeny.

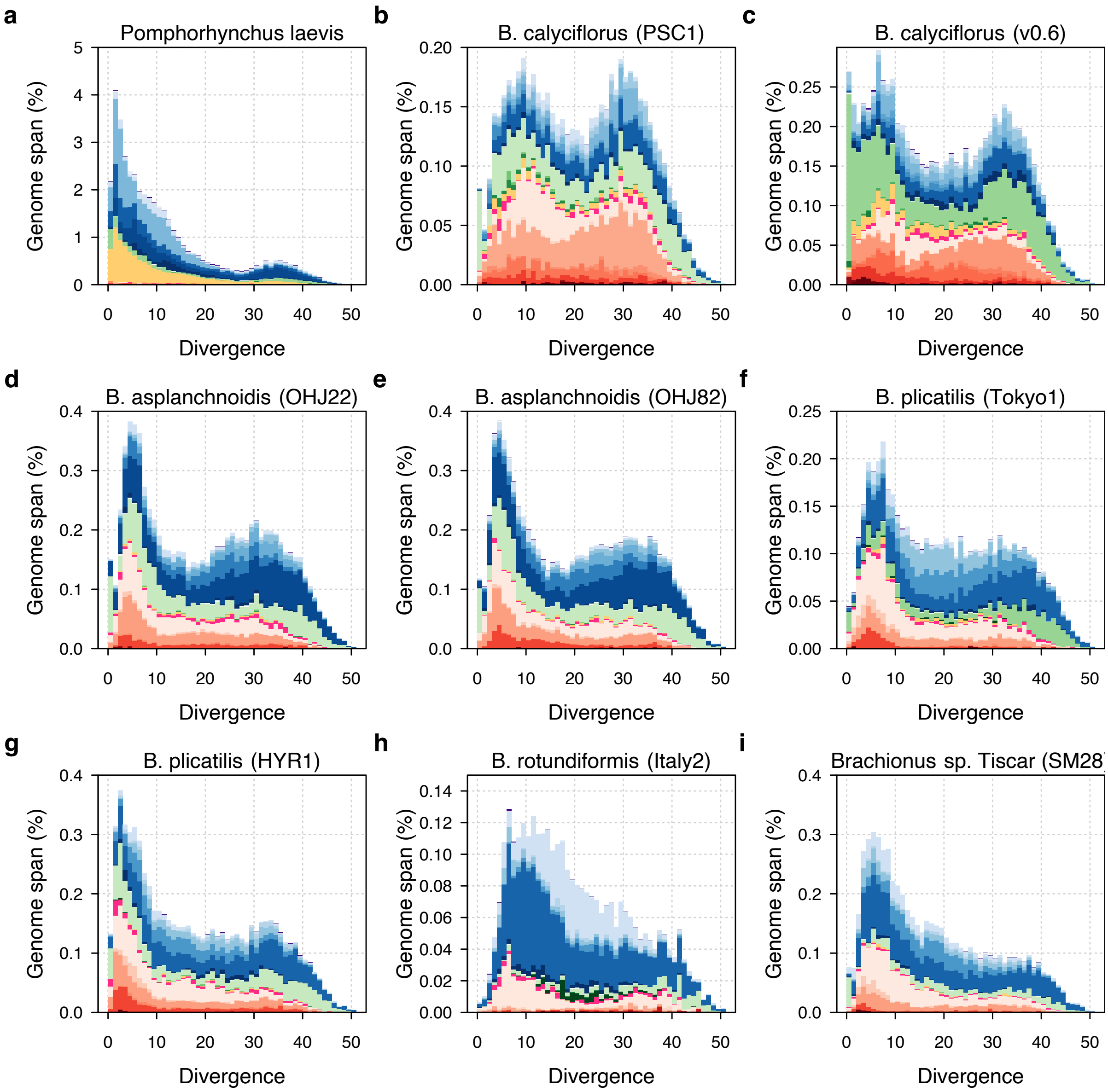

**S4 Fig. TE divergence landscapes for individual *Brachionus* genomes.** The *X*-axis shows the level of divergence (Kimura substitution level, CpG adjusted) between each identified TE copy and the computed consensus sequence for that TE family (inferred ancestral copy). Thus, if newly arising TE copies evolve neutrally, the amount of divergence is a proxy for the time since duplication for each copy, with older copies accumulating more substitutions and appearing further to the right on the *X*-axis. The *Y*-axis shows the proportion of the genome occupied by each TE divergence category, broken down by family where shades of colour indicate TE superfamily (red, DNA transposons; pink, rolling circles; yellow, PLEs; green, LTRs; blue, LINEs; purple, SINEs; see legend in main text).

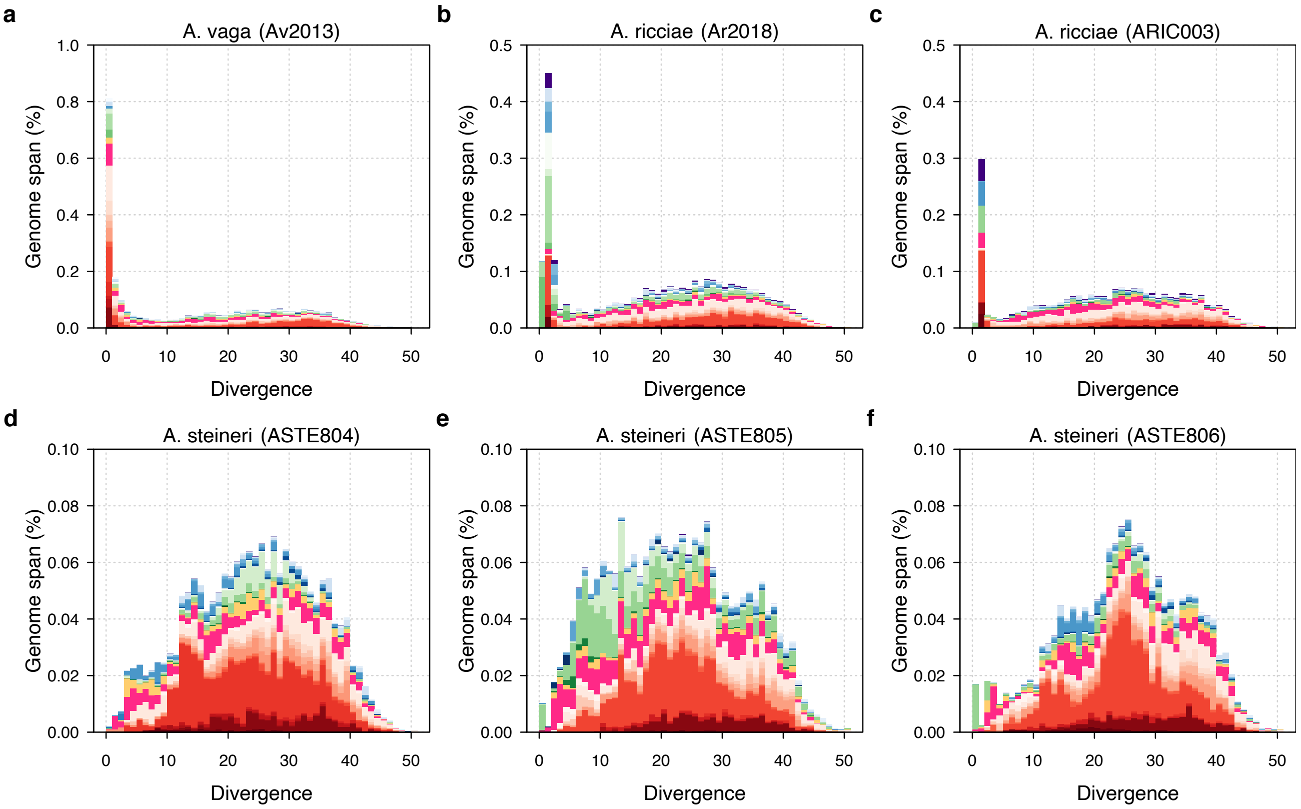

**S4 Fig—continued. TE divergence landscapes for individual *Adineta* genomes.** Plot is arranged as above. Landscape plots for the published *A. ricciae* (GCA_900240375.1) and *A. vaga* (GCA_000513175.1) assemblies are shown in **a** and **b**, respectively.

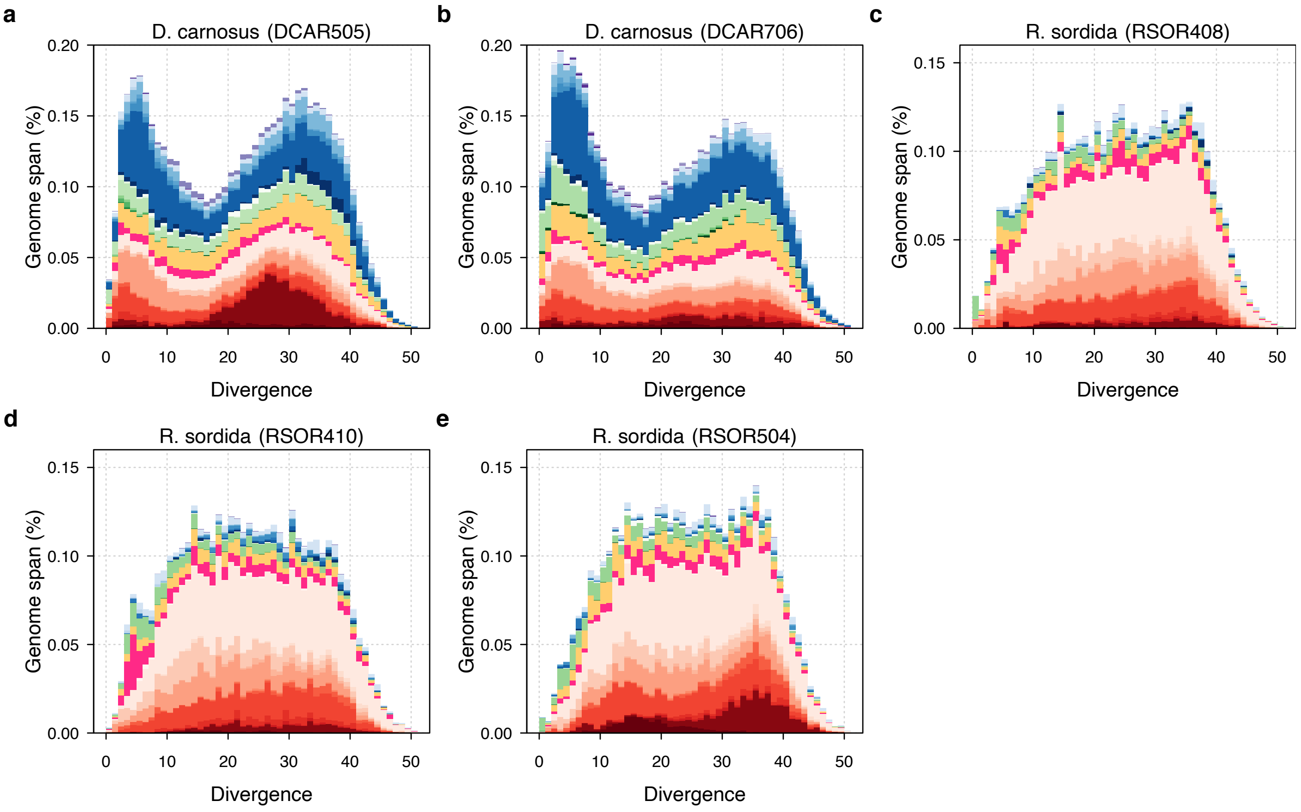

**S4 Fig—continued. TE divergence landscapes for individual *D. carnosus* and *R. sordida* genomes.** Plot is arranged as above.

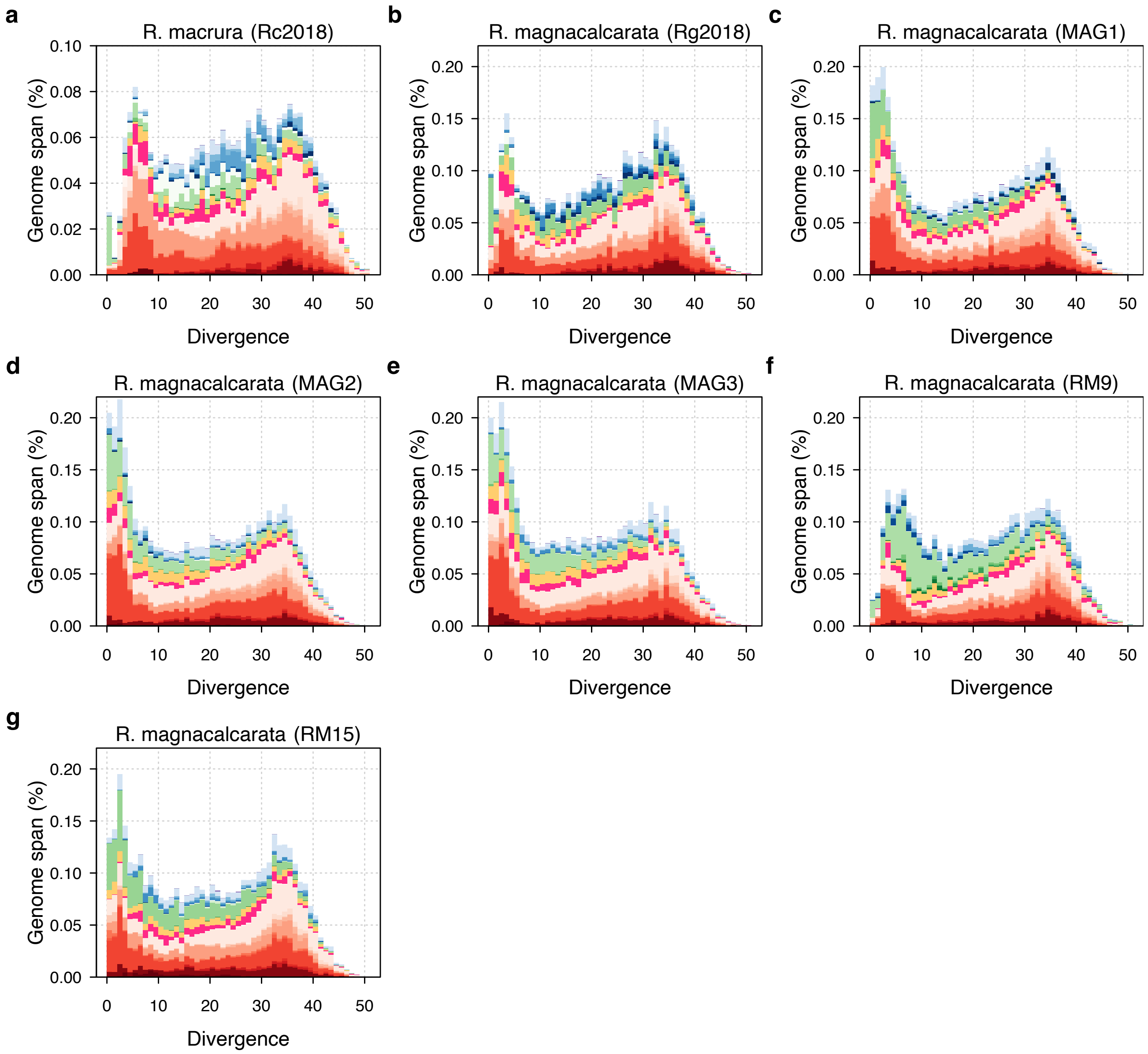

**S4 Fig—continued. TE divergence landscapes for *R. macrura* and individual *R. magnacalcarata* genomes.** Plot is arranged as above. Landscape plots for the published *R. macrura* (GCA_900239685.1) and *R. magnacalcarata* (GCA_900239745.1) assemblies are shown in **a** and **b**, respectively.

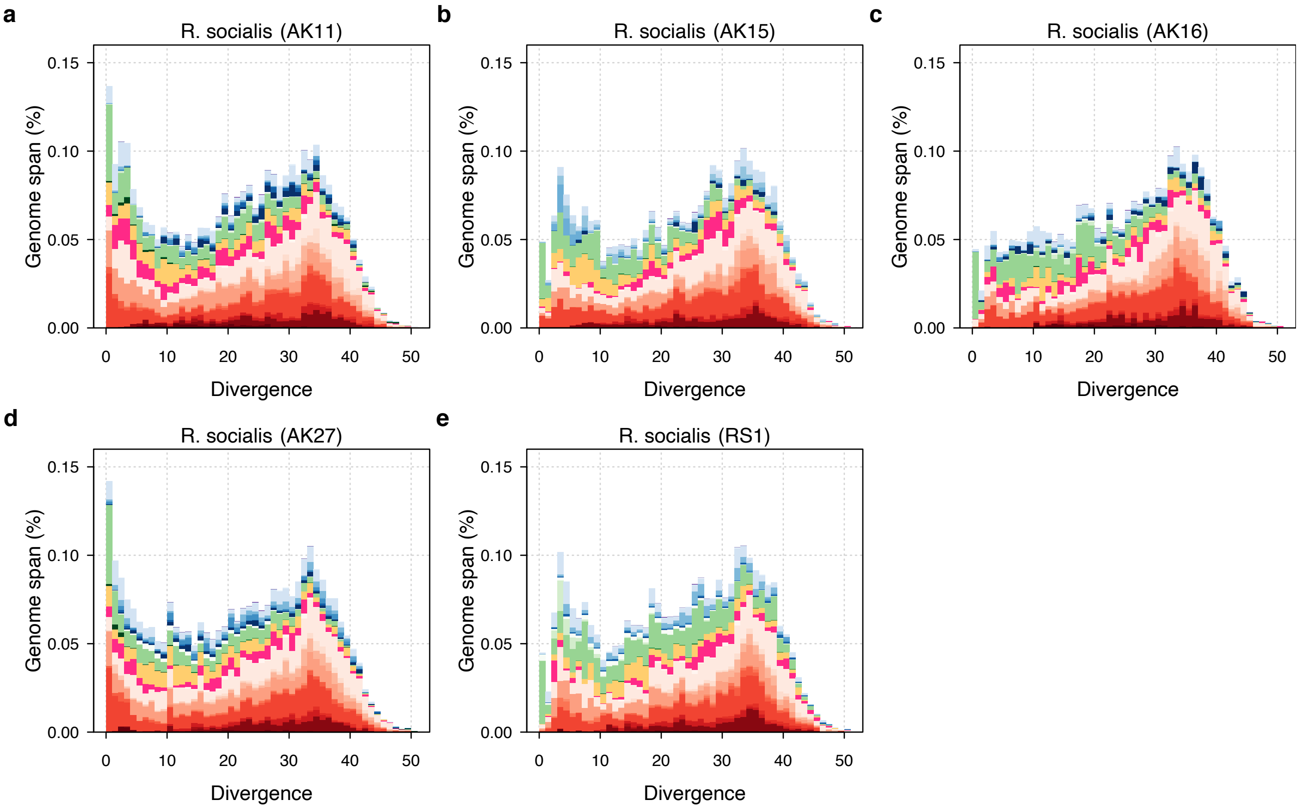

**S4 Fig—continued. TE divergence landscapes for individual *R. socialis* genomes.** Plot is arranged as above.

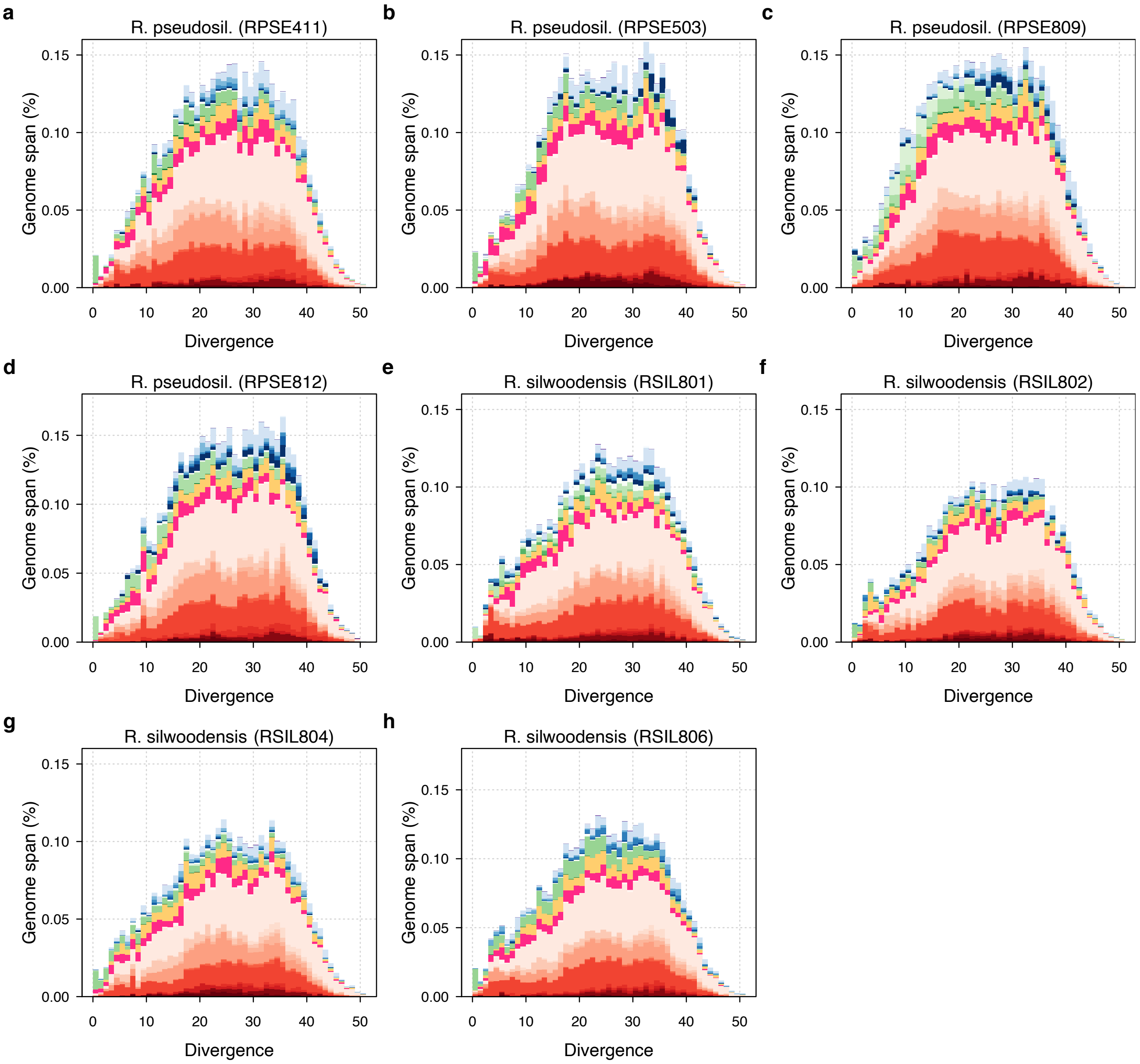

**S4 Fig—continued. TE divergence landscapes for individual *R.* sp. 'Silwood-1' and *R.* sp. 'Silwood-2' genomes.** Plot is arranged as above.

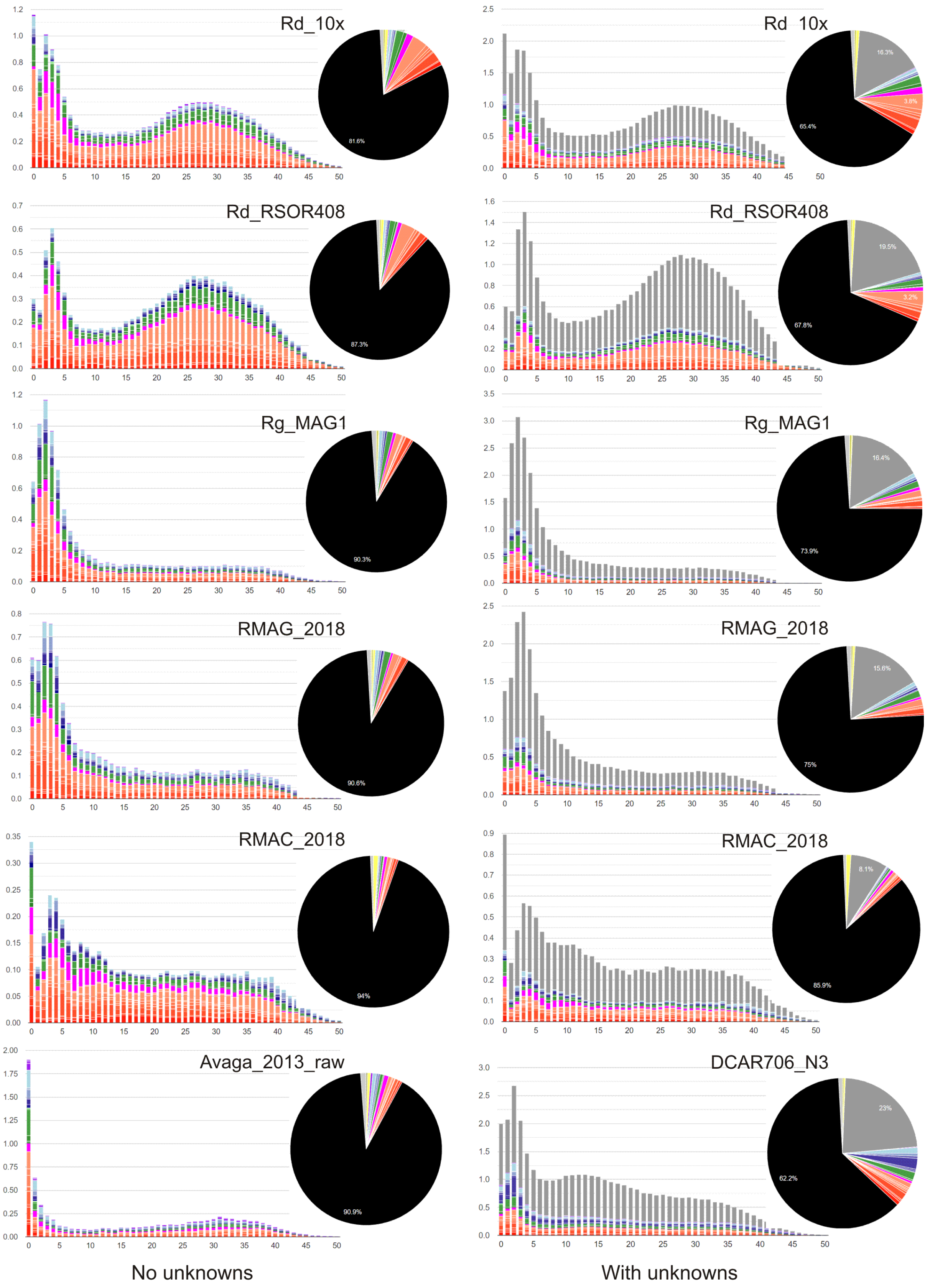

**S5 Fig. TE divergence landscapes for selected species constructed using REPET** (unfiltered, with default parameters). Landscapes and axes are arranged as previously. Pie charts show TE superfamilies occupying ≥ 2% of a given genome. Colour scheme from RepeatMasker v.4.1.0: orange, cut-and-paste DNA TEs; magenta, rolling-circle DNA TEs; green, LTR-Rs; dark blue, LINEs (non-LTR); light blue, PLEs; yellow, simple repeats; grey, unknown (unclassified). Note a greater proportion of repeats at lower divergence values (*K*-distance < 5).

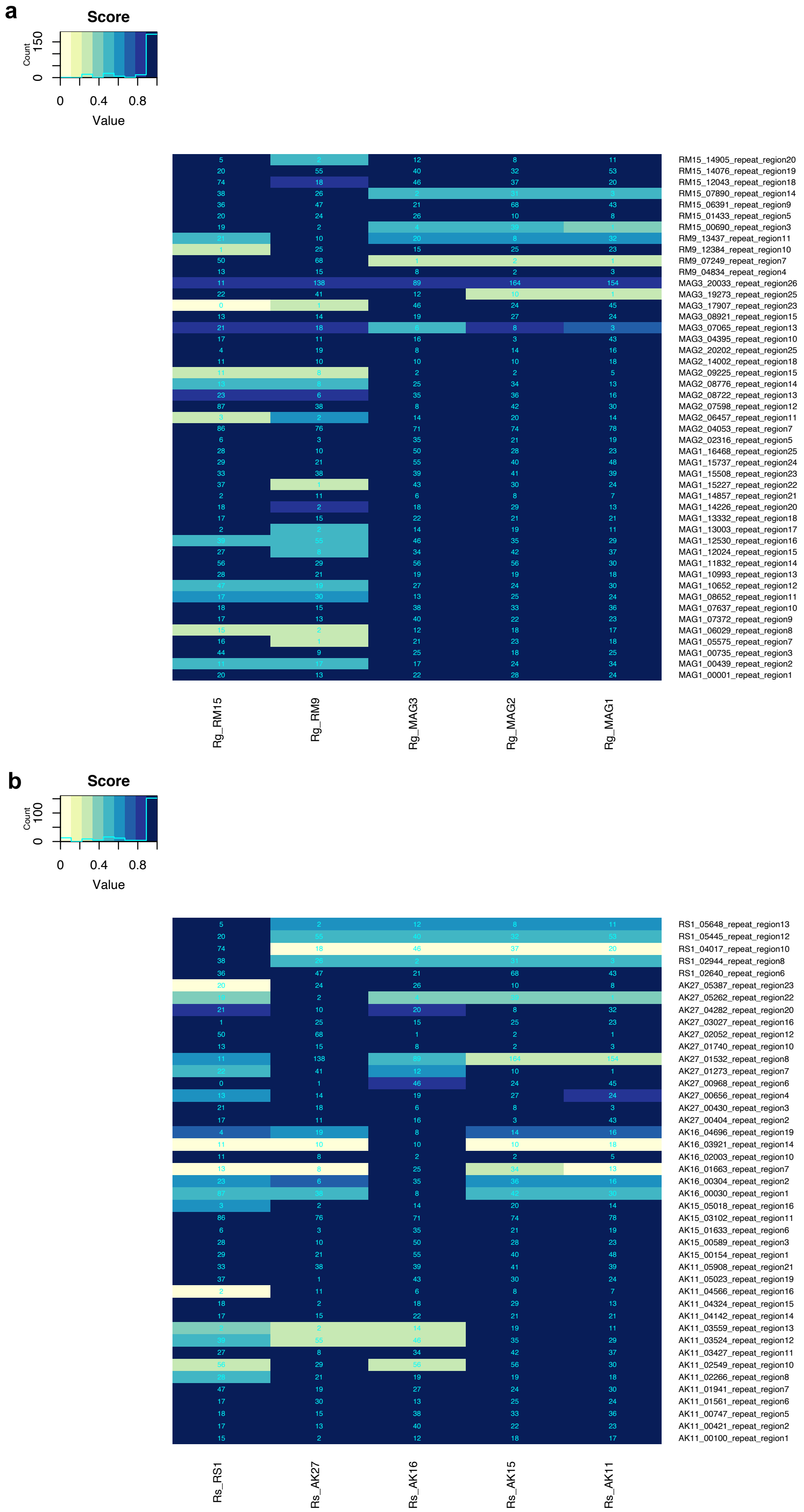

**S6 Fig. LTR-R polymorphism details.** Data are shown for **(a)** *R. magnacalcarata* and **(b)** *R. socialis*. Numbers in cells indicate the number of reads mapped to the LTR-tag location (see Materials and methods).

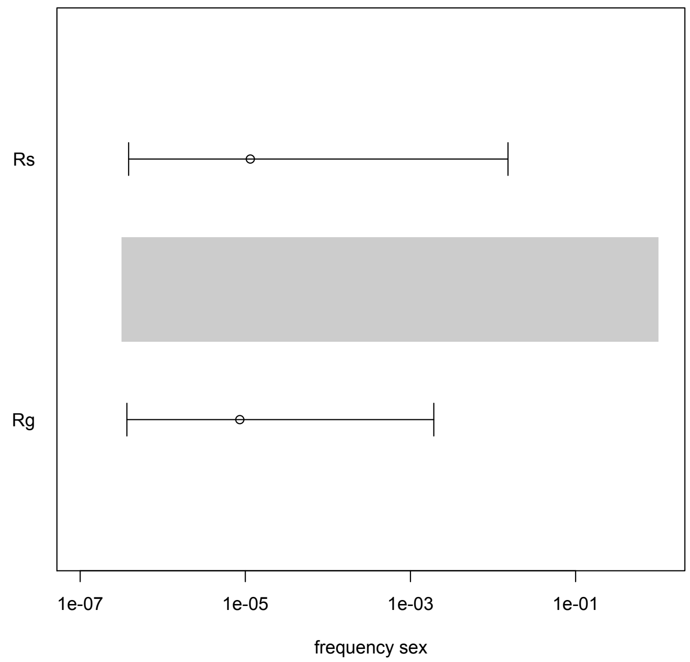

**S7 Fig. The median and 95% Highest Posterior Density interval of the frequency of sexual recombination affecting the presence/absence of LTR polymorphisms in *R. magnacalcarata* and *R. socialis*.** The grey bar shows the prior distribution used in simulations for the Approximate Bayesian Computation analysis. Confidence limits are broad but encompass the lowest range of the prior and hence cannot reject the hypothesis that recombination is absent for these loci.

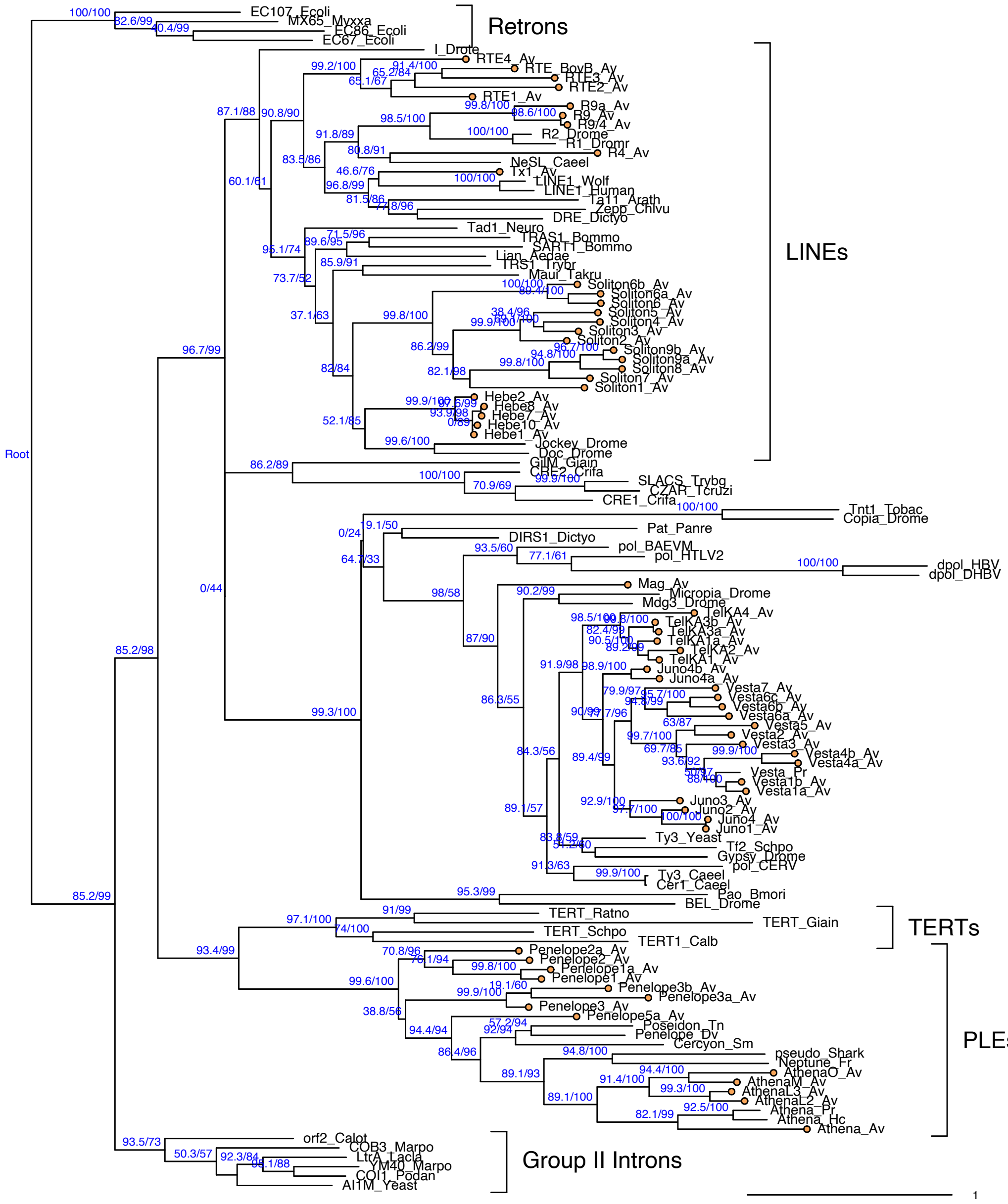

**S8 Fig. Maximum likelihood phylogeny for diverse reverse transcriptase domains from across the tree of life.** Major RT clades are indicated with labels. Sequences from the bdelloid *A. vaga* are indicated with orange tip labels and the suffix "_Av" (see [1,2] for further details). Phylogeny is rooted on branch leading to bacterial retrons. Numbers on branches represent branching support (%) based on 1,000 SH-aLRT and 5,000 UFBoot samples, respectively. Scale bar indicates 1 amino acid substitutions per site.

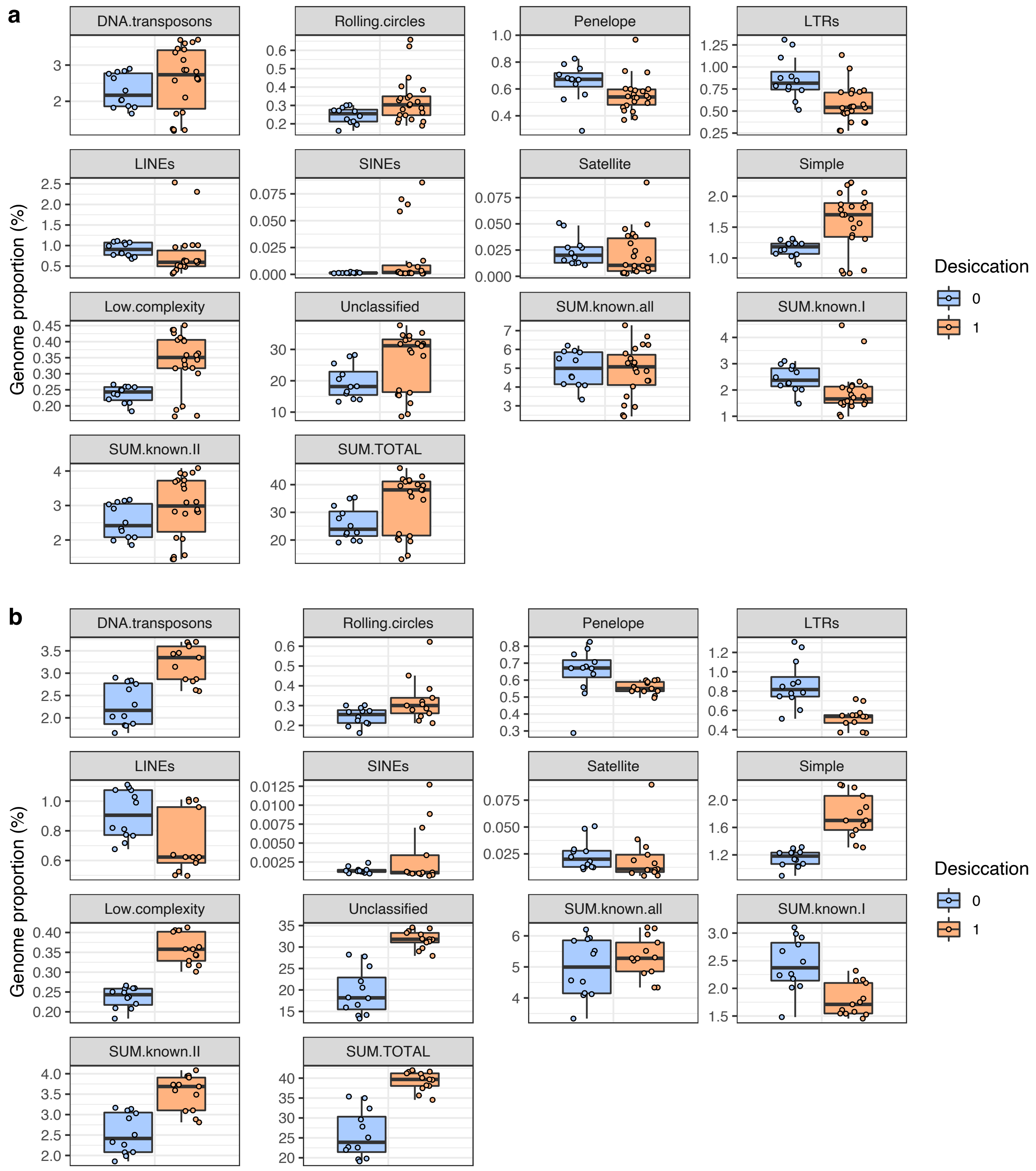

**S9 Fig. Repeat content in desiccating and nondesiccating bdelloids.** Data is shown for the proportion (%) of genome span accounted for each repeat type, plotted for **(a)** all bdelloids and **(b)** only *Rotaria* lineages. Desiccation tolerance indicated in blue (nondesiccating) and orange (desiccating). Boxplots show the median (band), interquartile range (box) and minimum/maximum values (whiskers). Underlying data are shown as jittered points. Summed categories defined as follows: '**known.all**', all classified TEs (i.e., satellite, simple, low complexity and unclassified excluded); '**known.I**', class I retrotransposon families; '**known.II**', class II DNA transposon families; '**TOTAL**', grand total of all repeats.

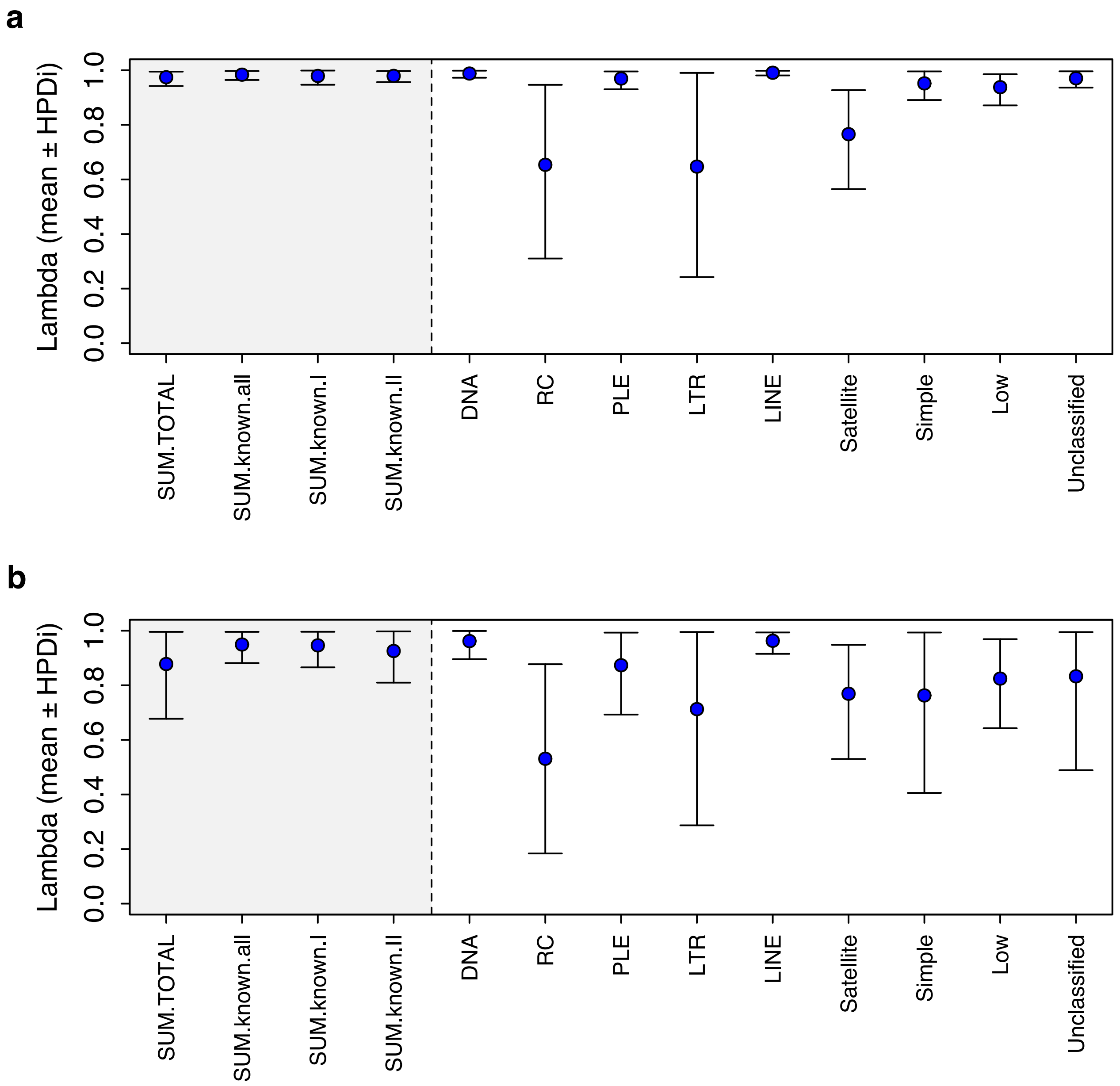

**S10 Fig. Phylogenetic signal in the effect of desiccation on TE content.** Error bars indicate the 95% credible interval around the mean value of $\lambda$ from 2,000 stored samples of the posterior. Values closer to 1 indicate a stronger phylogenetic effect. Results are shown for TE content modelled across **(a)** all bdelloids, and **(b)** only *Rotaria* samples in the phylogeny. Summed categories defined as follows: '**known.all**', all classified TEs (i.e., satellite, simple, low complexity and unclassified excluded); '**known.I**', class I retrotransposon families; '**known.II**', class II DNA transposon families; '**TOTAL**', grand total of all repeats.

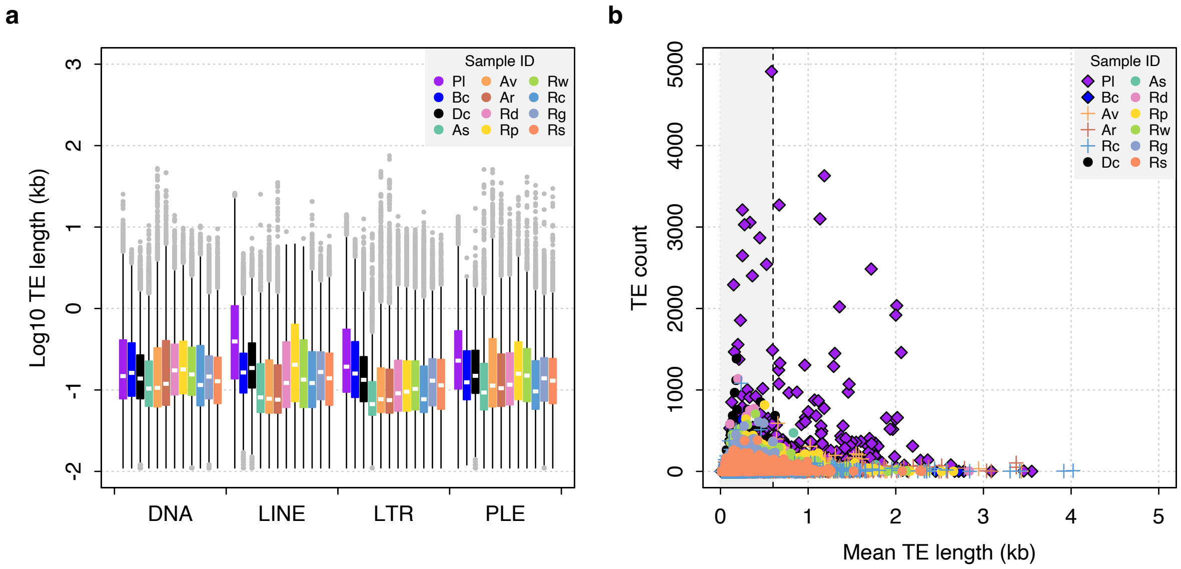

**S11 Fig. Distribution of TE lengths including the acanthocephalan *P. laevis*.** Boxplots show the median (white band), interquartile range (box) and minimum/maximum values (whiskers; outliers shown in grey).

**a**

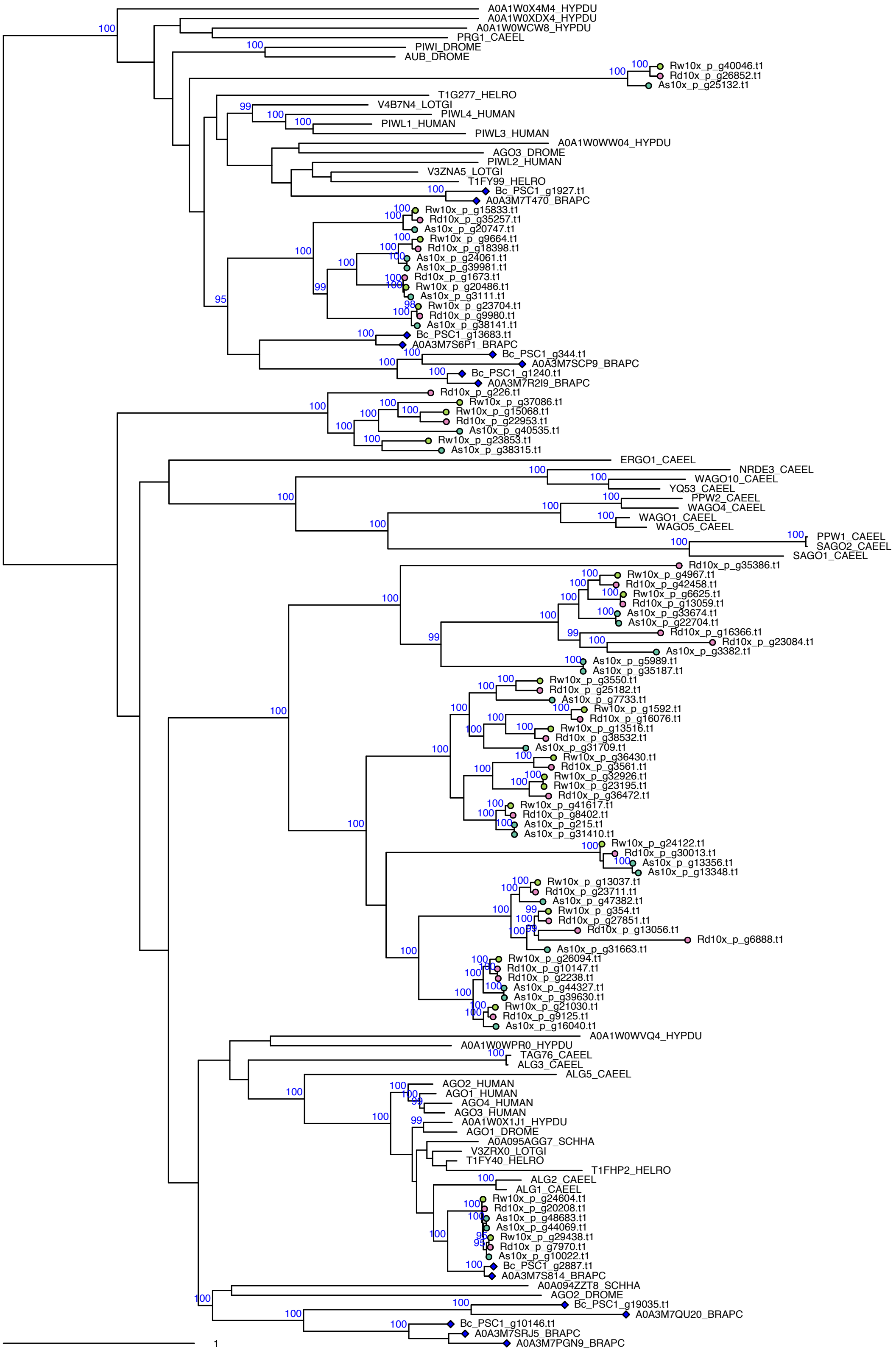
A0A1W0X4M4_HYPDU
A0A1W0XDX4_HYPDU
A0A1W0WCW8_HYPDU
PRG1_CAEEL
PIWI_DROME
AUB_DROME
Rw10x_p_g40046.t1
Rd10x_p_g26852.t1
As10x_p_g25132.t1
T1G277_HELRO
V4B7N4_LOTGI
PIWL4_HUMAN
PIWL1_HUMAN
PIWL3_HUMAN
A0A1W0WW04_HYPDU
AGO3_DROME
PIWL2_HUMAN
V3ZNA5_LOTGI
T1FY99_HELRO
Bc_PSC1_g1927.t1
A0A3M7T470_BRAPC
Rw10x_p_g15833.t1
Rd10x_p_g35257.t1
As10x_p_g20747.t1
Rw10x_p_g9664.t1
Rd10x_p_g18398.t1
As10x_p_g24061.t1
As10x_p_g39981.t1
Rd10x_p_g1673.t1
Rw10x_p_g20486.t1
As10x_p_g3111.t1
Rw10x_p_g23704.t1
Rd10x_p_g9980.t1
As10x_p_g38141.t1
Bc_PSC1_g13683.t1
A0A3M7S6P1_BRAPC
Bc_PSC1_g344.t1
A0A3M7SCP9_BRAPC
Bc_PSC1_g1240.t1
A0A3M7R2I9_BRAPC
Rd10x_p_g226.t1
Rw10x_p_g37086.t1
Rw10x_p_g15068.t1
Rd10x_p_g22953.t1
As10x_p_g40535.t1
Rw10x_p_g23853.t1
As10x_p_g38315.t1
ERGO1_CAEEL
NRDE3_CAEEL
WAGO10_CAEEL
YQ53_CAEEL
PPW2_CAEEL
WAGO4_CAEEL
WAGO1_CAEEL
WAGO5_CAEEL
PPW1_CAEEL
SAGO2_CAEEL
SAGO1_CAEEL
Rd10x_p_g35386.t1
Rw10x_p_g4967.t1
Rd10x_p_g42458.t1
Rw10x_p_g6625.t1
Rd10x_p_g13059.t1
As10x_p_g33674.t1
As10x_p_g22704.t1
Rd10x_p_g16366.t1
Rd10x_p_g23084.t1
As10x_p_g3382.t1
As10x_p_g5989.t1
As10x_p_g35187.t1
Rw10x_p_g3550.t1
Rd10x_p_g25182.t1
As10x_p_g7733.t1
Rw10x_p_g1592.t1
Rd10x_p_g16076.t1
Rw10x_p_g13516.t1
Rd10x_p_g38532.t1
As10x_p_g31709.t1
Rw10x_p_g36430.t1
Rd10x_p_g3561.t1
Rw10x_p_g32926.t1
Rw10x_p_g23195.t1
Rd10x_p_g36472.t1
Rw10x_p_g41617.t1
Rd10x_p_g8402.t1
As10x_p_g215.t1
As10x_p_g31410.t1
Rw10x_p_g24122.t1
Rd10x_p_g30013.t1
As10x_p_g13356.t1
As10x_p_g13348.t1
Rw10x_p_g13037.t1
Rd10x_p_g23711.t1
As10x_p_g47382.t1
Rw10x_p_g354.t1
Rd10x_p_g27851.t1
Rd10x_p_g13056.t1
Rd10x_p_g6888.t1
As10x_p_g31663.t1
Rw10x_p_g26094.t1
Rd10x_p_g10147.t1
Rd10x_p_g2238.t1
As10x_p_g44327.t1
As10x_p_g39630.t1
Rw10x_p_g21030.t1
Rd10x_p_g9125.t1
As10x_p_g16040.t1
A0A1W0WVQ4_HYPDU
A0A1W0WPR0_HYPDU
TAG76_CAEEL
ALG3_CAEEL
ALG5_CAEEL
AGO2_HUMAN
AGO1_HUMAN
AGO4_HUMAN
AGO3_HUMAN
A0A1W0X1J1_HYPDU
AGO1_DROME
A0A095AGG7_SCHHA
V3ZRX0_LOTGI
T1FY40_HELRO
T1FHP2_HELRO
ALG2_CAEEL
ALG1_CAEEL
Rw10x_p_g24604.t1
Rd10x_p_g20208.t1
As10x_p_g48683.t1
As10x_p_g44069.t1
Rw10x_p_g29438.t1
Rd10x_p_g7970.t1
As10x_p_g10022.t1
Bc_PSC1_g2887.t1
A0A3M7S814_BRAPC
A0A094ZZT8_SCHHA
AGO2_DROME
Bc_PSC1_g19035.t1
A0A3M7QU20_BRAPC
Bc_PSC1_g10146.t1
A0A3M7SRJ5_BRAPC
A0A3M7PGN9_BRAPC
1

**b**

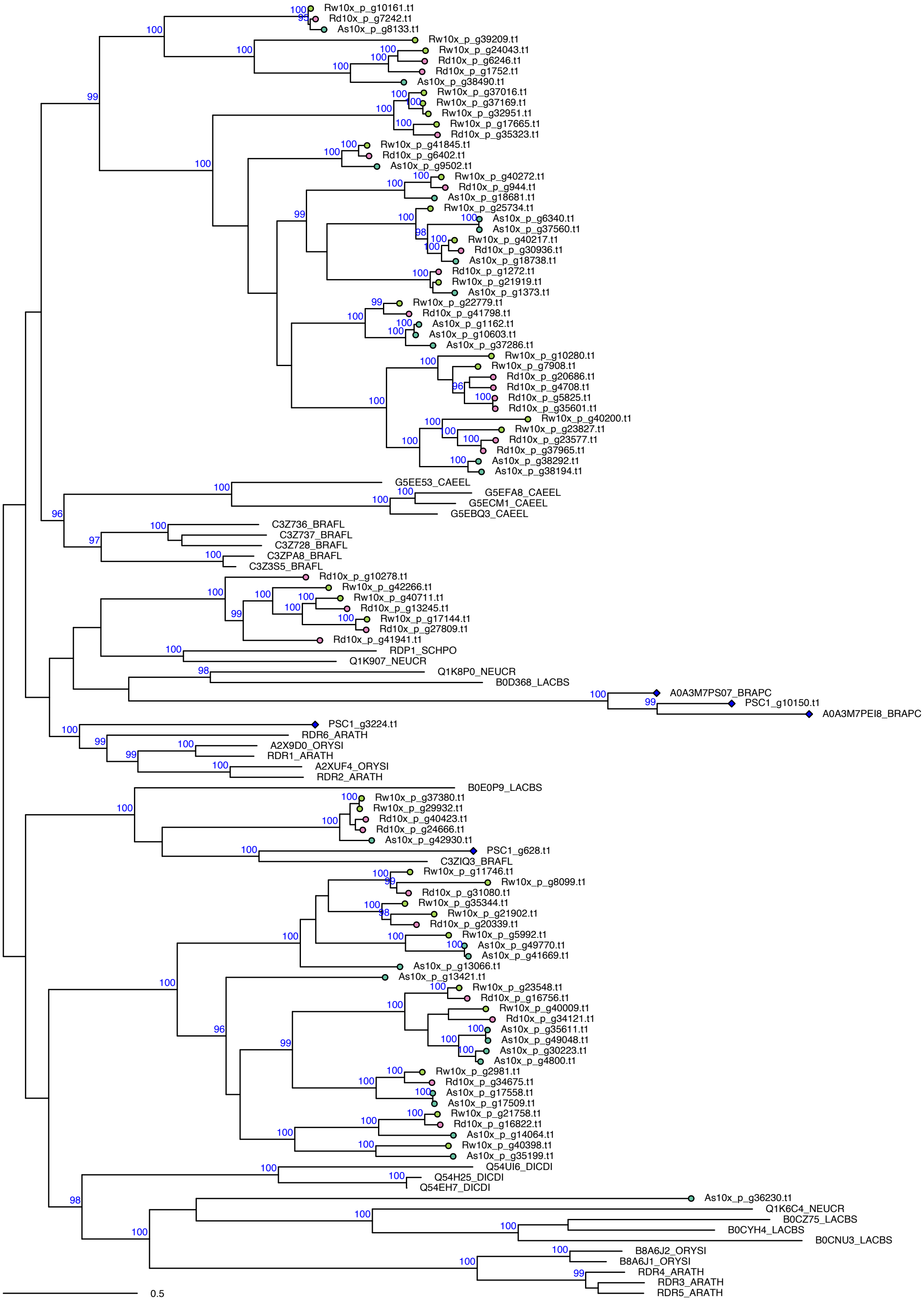

Rw10x_p_g10161.t1
Rd10x_p_g7242.t1
As10x_p_g8133.t1
Rw10x_p_g39209.t1
Rw10x_p_g24043.t1
Rd10x_p_g6246.t1
Rd10x_p_g1752.t1
As10x_p_g38490.t1
Rw10x_p_g37016.t1
Rw10x_p_g37169.t1
Rw10x_p_g32951.t1
Rw10x_p_g17665.t1
Rd10x_p_g35323.t1
Rw10x_p_g41845.t1
Rd10x_p_g6402.t1
As10x_p_g9502.t1
Rw10x_p_g40272.t1
Rd10x_p_g944.t1
As10x_p_g18681.t1
Rw10x_p_g25734.t1
As10x_p_g6340.t1
As10x_p_g37560.t1
Rw10x_p_g40217.t1
Rd10x_p_g30936.t1
As10x_p_g18738.t1
Rd10x_p_g1272.t1
Rw10x_p_g21919.t1
As10x_p_g1373.t1
Rw10x_p_g22779.t1
Rd10x_p_g41798.t1
As10x_p_g1162.t1
As10x_p_g10603.t1
As10x_p_g37286.t1
Rw10x_p_g10280.t1
Rw10x_p_g7908.t1
Rd10x_p_g20686.t1
Rd10x_p_g4708.t1
Rd10x_p_g5825.t1
Rd10x_p_g35601.t1
Rw10x_p_g40200.t1
Rw10x_p_g23827.t1
Rd10x_p_g23577.t1
Rd10x_p_g37965.t1
As10x_p_g38292.t1
As10x_p_g38194.t1
G5EE53_CAEEL
G5EFA8_CAEEL
G5ECM1_CAEEL
G5EBQ3_CAEEL
C3Z736_BRAFL
C3Z737_BRAFL
C3Z728_BRAFL
C3ZPA8_BRAFL
C3Z3S5_BRAFL
Rd10x_p_g10278.t1
Rw10x_p_g42266.t1
Rw10x_p_g40711.t1
Rd10x_p_g13245.t1
Rw10x_p_g17144.t1
Rd10x_p_g27809.t1
Rd10x_p_g41941.t1
RDP1_SCHPO
Q1K907_NEUCR
Q1K8P0_NEUCR
B0D368_LACBS
A0A3M7PS07_BRAPC
PSC1_g10150.t1
A0A3M7PEI8_BRAPC
PSC1_g3224.t1
RDR6_ARATH
A2X9D0_ORYSI
RDR1_ARATH
A2XUF4_ORYSI
RDR2_ARATH
B0E0P9_LACBS
Rw10x_p_g37380.t1
Rw10x_p_g29932.t1
Rd10x_p_g40423.t1
Rd10x_p_g24666.t1
As10x_p_g42930.t1
PSC1_g628.t1
C3ZIQ3_BRAFL
Rw10x_p_g11746.t1
Rw10x_p_g8099.t1
Rd10x_p_g31080.t1
Rw10x_p_g35344.t1
Rw10x_p_g21902.t1
Rd10x_p_g20339.t1
Rw10x_p_g5992.t1
As10x_p_g49770.t1
As10x_p_g41669.t1
As10x_p_g13066.t1
As10x_p_g13421.t1
Rw10x_p_g23548.t1
Rd10x_p_g16756.t1
Rw10x_p_g40009.t1
Rd10x_p_g34121.t1
As10x_p_g35611.t1
As10x_p_g49048.t1
As10x_p_g30223.t1
As10x_p_g4800.t1
Rw10x_p_g2981.t1
Rd10x_p_g34675.t1
As10x_p_g17558.t1
As10x_p_g17509.t1
Rw10x_p_g21758.t1
Rd10x_p_g16822.t1
As10x_p_g14064.t1
Rw10x_p_g40398.t1
As10x_p_g35199.t1
Q54UI6_DICDI
Q54H25_DICDI
Q54EH7_DICDI
As10x_p_g36230.t1
Q1K6C4_NEUCR
B0CZ75_LACBS
B0CYH4_LACBS
B0CNU3_LACBS
B8A6J2_ORYSI
B8A6J1_ORYSI
RDR4_ARATH
RDR3_ARATH
RDR5_ARATH
0.5

c

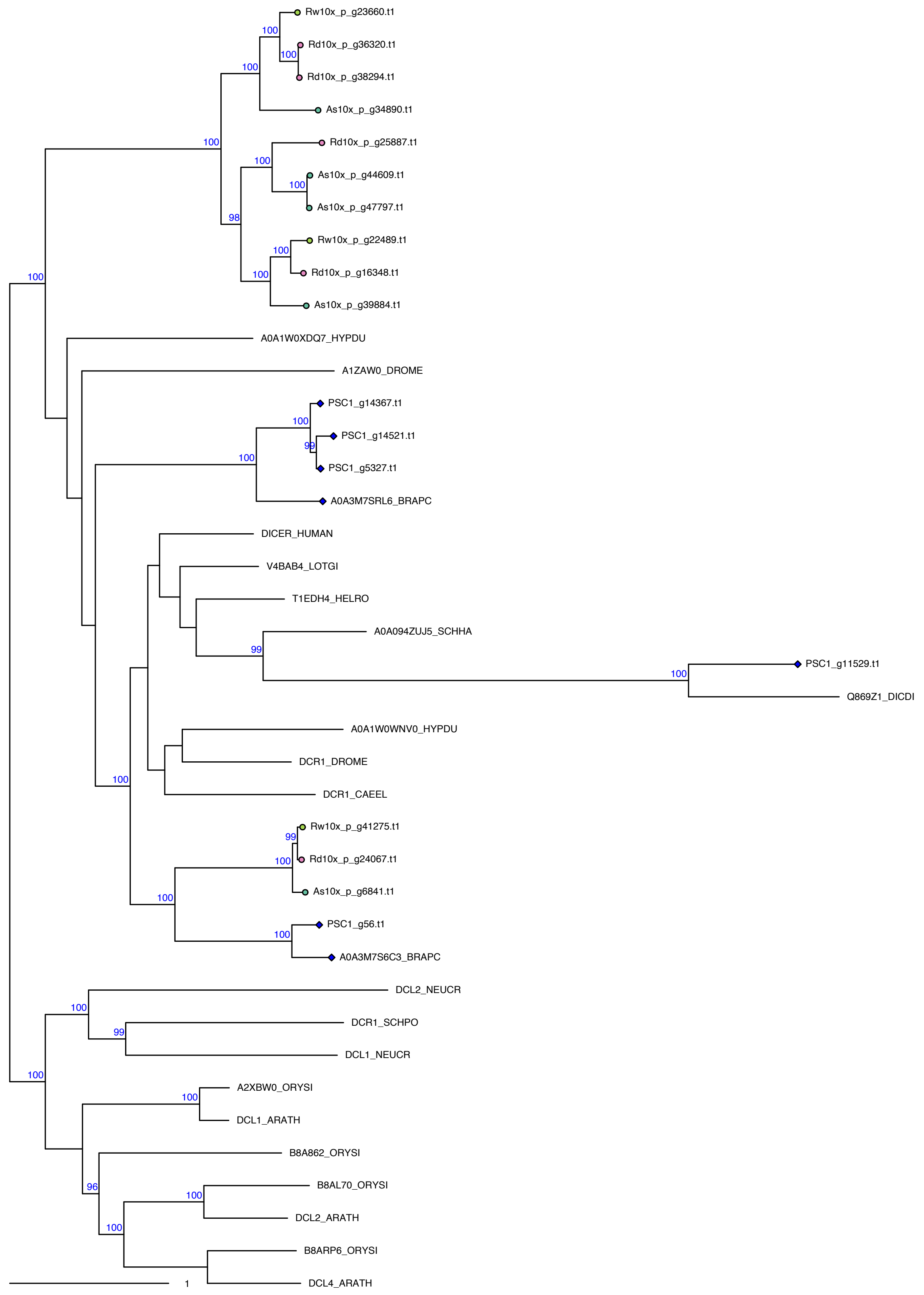

**S12 Fig. Argonaute, RdRP and Dicer phylogeny details.** Bdelloid copies are indicated with tip labels. Numbers on branches represent branching support (%) based on 5,000 UFBoot samples, respectively. Scale bar indicates 1 amino acid substitutions per site.

**S1 Note. Intragenomic divergence and collinearity.** Intragenomic divergence between homologous gene copies was estimated by counting the number of single nucleotide polymorphisms (SNPs) called inside coding regions (CDS). For each sample, read mapping (filtered data) and SNP calling was performed using BBTools 'bbmap' and 'callvariants' respectively, both with default settings. The number of high-quality SNPs intersecting with CDSs was counted using BEDTools v2.29.2 'intersect' [3] and divided by the total span of all CDSs to provide a simple measure of divergence between homologous gene copies. For the three 10x diploid ('megabubbles') genomes of *A. steineri*, *R. sordida* and *R. silwoodensis*, collinear blocks of five or more genes were identified using MCScanX [4], allowing a maximum of 10 gaps ('-m 10') in each block. Protein sequences for collinear pairs of genes were aligned using Clustalo v1.2.4 [5] and the number of synonymous ($K_S$) and nonsynonymous ($K_A$) substitutions were calculated using BioPerl [6]. A collinearity score was computed by dividing the number of collinear genes in a given block by the total number of genes in that block [2].

**S2 Note. Identification of putative telomeric repeats in rotifer genomes.** The telomeric repeat hexamer 'TGTGGG' is identified from *A. vaga* but may differ between species. For example, in the bdelloid species *Philodina roseola* it is known to be 'TGAGGG', with 'TGTGGG' found as a less-frequent variant [7]. In other cases, however, the telomeric repeat may be conserved even across large evolutionary distances (http://telomerase.asu.edu/sequences_telomere.html). The telomeric repeat in monogononts is not currently known. Thus, to test if the telomeric repeat sequence has diverged substantially in monogononts, we identified the most frequent G-rich hexamers that did not overlap with any predicted CDS (i.e. discounting any biases in hexamer frequency that may be due to amino acid composition) in the *B. calyciflorus* PSC1 genome assembly using BBTools 'kmercountexact' with the parameters 'k=6 mincount=5'. Visual inspection of the resulting output revealed the hexamers 'TGTGGG', 'TGGTGG' and 'TTGGGG' to be the most frequent and with similar occurrence in the genome (13,499, 13,076 and 16,357 exact matches to non-CDS regions, respectively; see Table below). for comparison, the equivalent counts in the *A. vaga* (Av2013) genome were 'TGTGGG' = 58,964, 'TGGTGG' = 36,269 and 'TTGGGG' = 9720, while in the distantly related bdelloid species *D. carnosus* DCAR706 the counts were 'TGTGGG' = 91,200, 'TGGTGG' = 123,684 and 'TTGGGG' = 74,904. Thus, in lieu of any substantial differences in hexamer frequency within species that might indicate a clear alternative candidate for *B. calyciflorus* (or *D. carnosus*), we decided to include only the frequency of occurrence of the known telomeric repeat 'TGTGGG' in all samples.

| Species | Genome | Frequency of repeat | | | Ratio |
|---|---|---|---|---|---|
| | | TGTGGG | TGGTGG | TTGGGG | |
| *A. vaga* | Av2013 | 58,964 | 36,269 | 9720 | 1.0:0.6:0.2 |
| *B. calyciflorus* | PSC1 | 13,499 | 13,076 | 16,357 | 1.0:1.0:1.2 |
| *D. carnosus* | DCAR706 | 91,200 | 123,684 | 74,904 | 1.0:1.4:0.8 |

**S1 Table.** Assembly statistics for 30 maximum-haplotig assemblies presented in this study.

| ID | SZ | NN | N50 | L50 | AU | GC | Gaps (kb) | Genome BUSCO | CDS | Proteome BUSCO | GenBank accession |
|---|---|---|---|---|---|---|---|---|---|---|---|
| **Ar_ARIC003** | 172.2 | 22,742 | 18.6 | 2,729 | 23.2 | 35.6 | 2 | C:91%,[S:27%,D:64%],F:5% | 64,538 | C:95%[S:14%,D:81%],F:3% | GCA_904047305.1 |
| **As_10x_m** | 231.1 | 8,705 | 199.2 | 285 | 289.3 | 28.8 | 290 | C:95%,[S:24%,D:71%],F:2% | 67,518 | C:97%[S:19%,D:78%],F:2% | GCA_904059895.1 |
| **As_ASTE804** | 163.7 | 34,092 | 20.8 | 2,068 | 29.7 | 29.1 | 6 | C:97%,[S:78%,D:19%],F:1% | 54,870 | C:99%[S:73%,D:26%],F:1% | GCA_904047245.1 |
| **As_ASTE805** | 166.6 | 39,093 | 22.4 | 1,837 | 33.6 | 29.3 | 36 | C:98%,[S:81%,D:17%],F:0% | 55,382 | C:99%[S:74%,D:25%],F:1% | GCA_904047255.1 |
| **As_ASTE806** | 163.8 | 32,429 | 22.7 | 1,846 | 32.9 | 29.1 | 18 | C:96%,[S:77%,D:19%],F:2% | 54,295 | C:97%[S:70%,D:27%],F:2% | GCA_904047275.1 |
| **Dc_DCAR505** | 344.1 | 121,494 | 7197 | 13,019 | 9,909 | 33.6 | 70 | C:86%[S:69%,D:17%],F:8% | 50,399 | C:89%[S:70%,D:18%],F:9% | GCA_904047325.1 |
| **Dc_DCAR706** | 401.6 | 165,346 | 10523 | 9,156 | 17,581 | 33.7 | 98 | C:95%[S:70%,D:25%],F:2% | 51,520 | C:95%[S:71%,D:25%],F:2% | GCA_904053135.1 |
| **Rd_10x_m** | 360.6 | 18,468 | 71.9 | 1,262 | 107.6 | 30.0 | 388 | C:93%,[S:23%,D:70%],F:1% | 61,901 | C:96%[S:21%,D:75%],F:1% | GCA_904059885.1 |
| **Rd_RSOR408** | 273.4 | 73,081 | 17.8 | 3,788 | 28 | 30.6 | 16 | C:93%,[S:76%,D:18%],F:3% | 44,351 | C:96%[S:74%,D:22%],F:3% | GCA_904053145.1 |
| **Rd_RSOR410** | 272.4 | 69,564 | 21.2 | 3,120 | 33.8 | 30.5 | 10 | C:95%,[S:81%,D:15%],F:2% | 43,594 | C:97%[S:76%,D:21%],F:2% | GCA_904053495.1 |
| **Rd_RSOR504** | 264.2 | 69,184 | 15.6 | 4,305 | 23.6 | 30.5 | 16 | C:93%,[S:81%,D:12%],F:4% | 44,209 | C:95%[S:77%,D:19%],F:4% | GCA_904047315.1 |
| **Rg_MAG1** | 241 | 293,619 | 2.8 | 17,705 | 5.6 | 31.9 | 1,785 | C:82%,[S:74%,D:8%],F:14% | 79,415 | C:89%[S:78%,D:11%],F:11% | GCA_903995865.1 |
| **Rg_MAG2** | 262 | 386,620 | 1.8 | 28,222 | 4.3 | 31.9 | 1,823 | C:76%,[S:71%,D:5%],F:19% | 88,549 | C:84%[S:74%,D:10%],F:16% | GCA_903995855.1 |
| **Rg_MAG3** | 258.7 | 374,008 | 1.9 | 26,450 | 4.6 | 31.9 | 1,822 | C:78%,[S:68%,D:10%],F:18% | 86,934 | C:85%[S:69%,D:16%],F:14% | GCA_903995845.1 |
| **Rg_RM9** | 176.6 | 38,569 | 16.8 | 2,801 | 23.2 | 31.9 | 223 | C:94%,[S:79%,D:15%],F:3% | 39,153 | C:96%[S:77%,D:20%],F:2% | GCA_903995825.1 |
| **Rg_RM15** | 198.5 | 126,446 | 10.4 | 4,693 | 15.7 | 32.0 | 766 | C:93%,[S:79%,D:14%],F:5% | 51,592 | C:97%[S:79%,D:18%],F:3% | GCA_903995885.1 |
| **Rp_RPSE411** | 308.8 | 61,655 | 18.2 | 4,528 | 25.6 | 31.1 | 8 | C:92%,[S:75%,D:17%],F:4% | 52,815 | C:93%[S:72%,D:21%],F:6% | GCA_904053095.1 |
| **Rp_RPSE503** | 296.4 | 68,725 | 13 | 6,328 | 17.4 | 31.4 | 12 | C:90%,[S:75%,D:15%],F:7% | 53,244 | C:91%[S:70%,D:21%],F:8% | GCA_904053125.1 |
| **Rp_RPSE809** | 295.1 | 107,316 | 12 | 6,719 | 16.3 | 31.2 | 29 | C:89%,[S:76%,D:13%],F:6% | 57,469 | C:92%[S:75%,D:18%],F:6% | GCA_904053085.1 |
| **Rp_RPSE812** | 274.1 | 65,539 | 11 | 6,936 | 14.7 | 31.2 | 9 | C:87%,[S:74%,D:13%],F:9% | 52,347 | C:89%[S:74%,D:16%],F:9% | GCA_904053105.1 |
| **Rs_AK11** | 175 | 115,181 | 11.6 | 3,687 | 17.7 | 31.8 | 968 | C:91%,[S:80%,D:11%],F:5% | 50,678 | C:94%[S:76%,D:18%],F:5% | GCA_903995835.1 |
| **Rs_AK15** | 151.3 | 21,443 | 34.3 | 1,245 | 44.7 | 31.8 | 77 | C:96%,[S:79%,D:18%],F:1% | 34,893 | C:98%[S:77%,D:21%],F:1% | GCA_903995795.1 |
| **Rs_AK16** | 151.3 | 18,665 | 40.3 | 1,049 | 53.4 | 31.9 | 78 | C:96%,[S:79%,D:17%],F:3% | 34,390 | C:98%[S:78%,D:20%],F:2% | GCA_903995805.1 |
| **Rs_AK27** | 174.5 | 109,049 | 13 | 3,377 | 19.1 | 31.8 | 837 | C:91%,[S:78%,D:14%],F:5% | 49,012 | C:95%[S:78%,D:17%],F:5% | GCA_903995815.1 |
| **Rs_RS1** | 154.6 | 20,987 | 35 | 1,224 | 46.2 | 31.8 | 96 | C:97%,[S:80%,D:17%],F:0% | 34,499 | C:99%[S:79%,D:20%],F:1% | GCA_903995875.1 |
| **Rw_10x_m** | 475.8 | 18,466 | 298.9 | 220 | 142 | 30.5 | 674 | C:94%,[S:16%,D:79%],F:1% | 70,338 | C:97%[S:9%,D:89%],F:1% | GCA_904059905.1 |
| **Rw_RSIL801** | 278.1 | 73,499 | 14.3 | 5,005 | 21.3 | 30.8 | 6 | C:90%,[S:79%,D:11%],F:6% | 47,030 | C:93%[S:75%,D:19%],F:5% | GCA_904053155.1 |
| **Rw_RSIL802** | 265.8 | 69,095 | 14.2 | 4,767 | 21.2 | 30.7 | 7 | C:87%,[S:75%,D:12%],F:9% | 46,049 | C:90%[S:73%,D:17%],F:8% | GCA_904053505.1 |
| **Rw_RSIL804** | 275.7 | 146,178 | 8.9 | 7,948 | 13 | 31.1 | 33 | C:87%,[S:75%,D:12%],F:8% | 51,958 | C:90%[S:71%,D:19%],F:8% | GCA_904053115.1 |
| **Rw_RSIL806** | 301.9 | 87,188 | 17.9 | 4,046 | 29.4 | 30.9 | 14 | C:94%,[S:79%,D:15%],F:4% | 48,353 | C:96%[S:76%,D:20%],F:3% | GCA_904054495.1 |

Sequence statistics: **SZ**, total sequence length (Mb); **NN**, number of sequences; **N50**, N50 scaffold length (kb); **L50**, N50 index; **AU**, expected scaffold size (area under Nx curve) (kb).
Gaps: Undetermined bases (Ns) introduced during scaffolding.
BUSCO score based on eukaryote set (n = 303); BUSCO codes: **C**, complete; **S**, complete and single-copy; **D**, complete and duplicated; **F**, fragmented.
Abbreviations: GC, guanine + cytosine; BUSCO, Benchmarking Universal Single-Copy Orthologs.

**S2 Table.** Genome properties of 16 protostome animals included in the study.

| Species | Phyla | Genome size | Assembly level | N50 | Genome BUSCO | GenBank Accession | Reference |
|---|---|---|---|---|---|---|---|
| *Drosophila melanogaster* | Arthropoda | 143.7 Mb | Chromosome | 21.5 Mb | C:99.7%[S:97.7%,D:2.0%],F:0.0% | GCF_000001215.4 | Adams et al. 2000 [8] |
| *Polypedilum nubifer* | Arthropoda | 115.7 Mb | Scaffold | 25.9 kb | C:97.0%[S:95.7%,D:1.3%],F:1.0% | PRJDB2914 | Gusev et al. 2014 [9] |
| *P. vanderplanki* | Arthropoda | 127.4 Mb | Scaffold | 228.6 kb | C:97.3%[S:94.7%,D:2.6%],F:1.7% | PRJDB1558 | Gusev et al. 2014 [9] |
| *Caenorhabditis elegans* | Nematoda | 100.3 Mb | Complete | 17.5 Mb | C:96.4%[S:94.7%,D:1.7%],F:0.0% | GCF_000002985.6 | C. elegans Sequencing Consortium 1998 [10] |
| *Hypsibius exemplaris* | Tardigrada | 104.2 Mb | Scaffold | 76.8 kb | C:93.0%[S:91.7%,D:1.3%],F:1.7% | GCA_002082055.1 | Hashimoto et al. 2016 [11] |
| *Ramazzottius varieornatus* | Tardigrada | 55.8 Mb | Scaffold | 129.7 kb | C:95.7%[S:94.4%,D:1.3%],F:0.7% | GCA_001949185.1 | Yoshida et al. 2017 [12] |
| *Octopus bimaculoides* | Mollusca | 2.3 Gb | Scaffold | 5.5 kb | C:86.5%[S:85.8%,D:0.7%],F:6.3% | GCF_001194135.1 | Albertin et al. 2015 [13] |
| *Crassostrea gigas* | Mollusca | 564.8 Mb | Scaffold | 42.3 kb | C:94.1%[S:89.8%,D:4.3%],F:1.7% | GCF_000297895.1 | Zhang et al. 2012 [14] |
| *Lottia gigantea* | Mollusca | 359.5 Mb | Scaffold | 96.0 kb | C:94.4%[S:92.4%,D:2.0%],F:1.3% | GCF_000327385.1 | Simakov et al. 2013 [15] |
| *Aplysia californica* | Mollusca | 927.3 Mb | Scaffold | 9.6 kb | C:92.4%[S:91.7%,D:0.7%],F:1.7% | GCF_000002075.1 | Broad Institute 2013 |
| *Biomphalaria glabrata* | Mollusca | 916.4 Mb | Scaffold | 7.3 kb | C:84.5%[S:82.8%,D:1.7%],F:7.9% | GCF_000457365.1 | Adema et al. 2017 [16] |
| *Helobdella robusta* | Annelida | 235.4 Mb | Scaffold | 52.2 kb | C:89.4%[S:88.1%,D:1.3%],F:4.3% | GCF_000326865.1 | Simakov et al. 2013 [13] |
| *Capitella teleta* | Annelida | 333.3 Mb | Scaffold | 21.9 kb | C:98.4%[S:94.4%,D:4.0%],F:0.3% | GCA_000328365.1 | Simakov et al. 2013 [13] |
| *Lingula anatina* | Brachiopoda | 406.3 Mb | Scaffold | 56.1 kb | C:93.7%[S:68.3%,D:25.4%],F:2.0% | GCF_001039355.1 | Luo et al. 2015 [17] |
| *Schistosoma haematobium* | Platyhelminthes | 376.0 Mb | Scaffold | 317.5 kb | C:72.3%[S:71.6%,D:0.7%],F:12.9% | GCF_000699445.1 | Young et al. 2012 [18] |
| *Intoshia linei* | Orthonectida | 41.6 Mb | Scaffold | 26.0 kb | C:25.1%[S:24.4%,D:0.7%],F:9.2% | GCA_001642005.1 | Mikhailov et al. 2016 [19] |

Assembly information taken from NCBI where provided.
BUSCO score based on eukaryote set (n = 303); BUSCO codes: **C**, complete; **S**, complete and single-copy; **D**, complete and duplicated; **F**, fragmented.

**S3 Table.** Analysis of recombination in LTR-tag presence/absence data.

| **Species**[1] | ***n*** | **LTR loci**[2] | $\bar{r}_d$[3] | ***p*** | **Consistency index** | **Frequency of sex**[4] | | |
|---|---|---|---|---|---|---|---|---|
| | | | | | | Median | lower 95% | upper 95% |
| **Rg** | 5 | 16 | 0.81 | 0.001 | 1.0 | $8.57 \times 10^{-06}$ | $3.68 \times 10^{-07}$ | 0.0019169 |
| **Rs** | 5 | 5 | 0.50 | 0.007 | 1.0 | $1.15 \times 10^{-05}$ | $3.86 \times 10^{-07}$ | 0.01517881 |

[1] Species codes: **Rg**, *Rotaria magnacalcarata*; **Rs**, *R. socialis*.
[2] Number of LTR-tag loci that were parsimony informative, i.e. no uniform or single variant individual within the sample.
[3] $\bar{r}_d$ is the modified index of association by Agapow and Burt (2001), $p$ is probability of obtaining a value this high in a permutated dataset, i.e. with full out-crossing and recombination, calculated with the 'ia()' function in the R package poppr.
[4] Frequency of sex was calculated by Approximate Bayesian Computation against 50,000 simulated datasets generated by the 'FacSexCoalescent' simulator of Hartfield et al. (2018). Simulation parameters: population size 10000, mutation rate 0.001, 900 sites, cross-over rate 2.224694e-06, yielding scaled cross-over rate of 40, frequency of sexual versus asexual reproduction log-uniform prior ranging from $10^{-6.5}$ to 0.999999. From the simulation output we extracted informative sites (i.e. removing uniform or sites with just a single variant individual) matching the number of informative sites in the observed data (selected to be spread out, i.e. unlinked, along the simulated chromosome). ABC computation used $\bar{r}_d$ and Consistency Index as two statistics for inferences, and tolerance 0.05 with the 'rejection' method in the 'abc()' function in the abc package in R. Posterior distribution is summarized by the median value of the frequency of sex and the upper and lower 95% highest posterior density limits.

**S4 Table.** Posterior mean and 95% credible intervals for the effect of desiccation ability (model levels: nondesiccating = 0, desiccating = 1) on TE load from a MCMCglmm Gaussian model. pMCMC is defined as twice the posterior probability that the estimate is negative or positive (whichever probability is smallest).

| Test | Mean | L-95% | U-95% | pMCMC |
|---|---|---|---|---|
| All TEs | 8.899 | -5.643 | 25.762 | 0.255 |
| All known TEs | 0.714 | -1.567 | 3.084 | 0.507 |
| Class I TEs | -0.146 | -1.563 | 1.243 | 0.834 |
| Class II TEs | 0.858 | -0.441 | 2.371 | 0.208 |
| DNA transposons | 0.743 | -0.643 | 2.149 | 0.298 |
| Rolling circles | 0.109 | -0.073 | 0.292 | 0.186 |
| PLEs | 0.092 | -0.270 | 0.465 | 0.610 |
| LTRs | -0.213 | -0.579 | 0.188 | 0.232 |
| LINEs | -0.112 | -0.758 | 0.475 | 0.719 |
| Satellite | -0.003 | -0.072 | 0.069 | 0.930 |
| Simple | 0.587 | -0.215 | 1.439 | 0.140 |
| Low complexity | 0.131 | 0.002 | 0.278 | 0.057 |
| Unclassified repeats | 7.311 | -6.853 | 22.611 | 0.285 |

**S5 Table.** Fixed and random effects estimates for a linear mixed effects model for 'other genes'.

| Random effects | | | | | |
|---|---|---|---|---|---|
| **Groups** | Variance | Std. Dev. | | | |
| **Sample ID** | 0.1149 | 0.3389 | | | |
| **Residual** | 1.0333 | 1.0165 | | | |
| **Fixed effects** | | | | | |
| | **Estimate** | **Std. Error** | **T-value** | **Pr(>\|t\|)** | |
| **(Intercept)** | 7.790 | 0.340 | 22.908 | < 0.001 | *** |
| **is.bdelloid** | 0.054 | 0.358 | 0.151 | 0.884 | |
| **is.desiccating** | 0.033 | 0.028 | 1.164 | 0.245 | |
| **type.LINE** | -0.220 | 0.037 | -6.005 | < 0.001 | *** |
| **type.LTR** | -0.420 | 0.045 | -9.373 | < 0.001 | *** |
| **type.PLE** | -0.599 | 0.041 | -14.514 | < 0.001 | *** |

The reference levels for fixed factors were as follows: type = BUSCO, is.bdelloid = 0, is.desiccation = 0. Results are based on 29,158 observations of 'other genes' span from 11 samples IDs.

**S5 Table—continued.** Fixed and random effects estimates for a linear mixed effects model for 'other TEs'.

| Random effects | | | | | |
|---|---|---|---|---|---|
| **Groups** | Variance | Std. Dev. | | | |
| **Sample ID** | 0.1149 | 0.1149 | | | |
| **Residual** | 1.6796 | 1.2960 | | | |
| **Fixed effects** | | | | | |
| | **Estimate** | **Std. Error** | **T-value** | **Pr(>\|t\|)** | |
| **(Intercept)** | 5.724 | 0.341 | 16.776 | < 0.001 | *** |
| **is.bdelloid** | -0.471 | 0.359 | -1.313 | 0.226 | |
| **is.desiccating** | 0.006 | 0.039 | 0.149 | 0.881 | |
| **type.LINE** | 0.746 | 0.035 | 21.151 | < 0.001 | *** |
| **type.LTR** | 0.772 | 0.037 | 20.718 | < 0.001 | *** |
| **type.PLE** | 0.695 | 0.033 | 20.798 | < 0.001 | *** |

The reference levels for fixed factors were as follows: type = BUSCO, is.bdelloid = 0, is.desiccation = 0. Results are based on 20,189 observations of 'other TEs' span from 11 samples IDs.

**S5 Table—continued.** Fixed and random effects estimates for a linear mixed effects model for 'telomeric repeats'.

| Random effects | | |
|---|---|---|
| **Groups** | Variance | Std. Dev. |
| **Sample ID** | 0.057 | 0.239 |
| **Residual** | 0.711 | 0.843 |

| | | | | | |
|---|---|---|---|---|---|
| **Fixed effects** | | | | | |
| | **Estimate** | **Std. Error** | **T-value** | **Pr(>\|t\|)** | |
| **(Intercept)** | 2.062 | 0.241 | 8.553 | < 0.001 | *** |
| **is.bdelloid** | 0.335 | 0.253 | 1.324 | 0.222 | |
| **is.desiccating** | -0.069 | 0.030 | -2.283 | 0.023 | * |
| **type.LINE** | 0.282 | 0.030 | 9.418 | < 0.001 | *** |
| **type.LTR** | 0.325 | 0.034 | 9.476 | < 0.001 | *** |
| **type.PLE** | 0.661 | 0.029 | 22.941 | < 0.001 | *** |

The reference levels for fixed factors were as follows: type = BUSCO, is.bdelloid = 0, is.desiccation = 0. Results are based on 16,641 observations of 'telomeric repeats' span from 11 samples IDs. For the desiccation contrast, a $P = 0.023$ is not significant after Bonferroni correction for multiple testing.

**S6 Table.** Fixed and random effects estimates from a linear mixed effects model of TE lengths.

| Random effects | | | | | |
|---|---|---|---|---|---|
| **Groups** | Variance | Std. Dev. | | | |
| **Sample ID** | 0.048 | 0.220 | | | |
| **Residual** | 1.331 | 1.154 | | | |
| **Fixed effects** | | | | | |
| | **Estimate** | **Std. Error** | **T-value** | **Pr(>\|t\|)** | |
| **(Intercept)** | 5.174 | 0.099 | 52.497 | < 0.001 | *** |
| **is.bdelloid** | -0.006 | 0.005 | -1.230 | 0.219 | |
| **is.desiccating** | -0.099 | 0.129 | -0.766 | 0.462 | |
| **type.LINE** | 0.204 | 0.004 | 50.434 | < 0.001 | *** |
| **type.LTR** | -0.199 | 0.005 | -43.472 | < 0.001 | *** |
| **type.PLE** | -0.081 | 0.005 | -17.945 | < 0.001 | *** |

The reference levels for fixed factors were as follows: type = DNA, is.bdelloid = 0, is.desiccation = 0. Results are based on 664,420 observations of TE length from 12 samples IDs.

### Supplementary information reference list
